# Molecular switching of a DNA-sliding clamp to a repressor mediates long-range gene silencing

**DOI:** 10.1101/2024.02.16.579611

**Authors:** Thomas C. McLean, Francisco Balaguer-Pérez, Joshua Chandanani, Christopher M. Thomas, Clara Aicart-Ramos, Sophia Burick, Paul Dominic B. Olinares, Giulia Gobbato, Julia E. A. Mundy, Brian T. Chait, David M. Lawson, Seth A. Darst, Elizabeth A. Campbell, Fernando Moreno-Herrero, Tung B.K. Le

**Author notes:** co-first authors (Thomas C. McLean, Francisco Balaguer-Pérez, Joshua Chandanani).

## Abstract

Long-range gene regulation is rare in bacteria and is confined to the classical DNA looping model. Here, we use a combination of biophysical approaches, including X-ray crystallography and single-molecule analysis, to show that long-range gene silencing on the plasmid RK2, a source of multidrug resistance across diverse Gram-negative bacteria, is achieved cooperatively by a DNA-sliding clamp, KorB, and a clamp-locking protein, KorA. We find that KorB is a CTPase clamp that can entrap and slide along DNA to reach distal target promoters. We resolved the tripartite crystal structure of a KorB-KorA-DNA co-complex, revealing that KorA latches KorB into a closed-clamp state. KorA thus stimulates repression by stalling KorB sliding at target promoters to occlude RNA polymerase holoenzymes. Altogether, our findings explain the mechanistic basis for KorB role-switching from a DNA-sliding clamp to a co-repressor, and provide a new paradigm for the long-range regulation of gene expression.

## INTRODUCTION

Spatiotemporal gene expression in eukaryotes is commonly coordinated by transcriptional enhancers/silencers, DNA elements that form loops over long genomic distances to contact and regulate target-gene promoters^1^. By contrast, gene regulation over kilobase distances is rare in bacteria, and it remains to be determined whether other mechanisms, beyond DNA looping, are involved.

Long-range gene regulation by the DNA-binding protein KorB is essential for stable vertical inheritance and horizontal transmission of the broad-host-range multidrug resistance plasmid RK2, as well as for the fitness of its host bacterium^2–6^. KorB is a member of the same family of proteins as ParB, which is crucial for bacterial chromosome segregation^7–10^ and the founding member of a new class of CTP-dependent molecular switches^11,12^. However, in contrast to ParB, KorB also functions as a long-range silencer, repressing the expression of plasmid genes that promote replication and conjugative transfer^13^. The best understood KorB-mediated transcriptional repression requires a 16-bp silencer sequence called *OB* (operator of KorB) and the small DNA-binding protein KorA. *OB* is positioned either 4-10 bp (*OB*-proximal) or 45 bp to >1000 bp (*OB-*distal) upstream of target promoters^13–17^. KorB-mediated long-range gene silencing has been investigated for three decades yet is not fully understood. Conflicting models propose that KorB either polymerizes on DNA to reach the target promoter from a distal *OB* site, or promotes DNA looping over a long distance to connect *OB* and the target promoter, or a combination thereof^13^. Thus, there is currently no unifying model that explains: (i) the flexibility of KorB-mediated long-range gene repression, which is insensitive to both the helical position of *OB* and the *OB*-promoter distance^13^; (ii) the requirement of KorA to tighten KorB-mediated long-range repression^14,18–20^; and (iii) the evolution of a dual-function protein, KorB, that has roles in both plasmid inheritance and gene expression regulation^5,21^.

The discovery of ParB CTPase activity^11,12^ prompted us to revisit previous data and models of KorB-mediated gene expression. By combining biochemistry, structural biology, single-molecule *in vitro* reconstitution, native mass spectrometry, and *in vivo* gene expression assays, we reveal that KorB is a DNA-dependent CTPase. We find that KorB binds CTP to form a protein clamp that can entrap and slide along DNA to mediate long-range transcriptional repression, likely by allowing KorB to reach target promoters from distal *OB* sites. Unlike canonical ParB, KorB does not bridge and condense DNA. We resolved the tripartite crystal structure of a KorB-KorA-DNA co-complex, finding that KorA is a clamp-locking protein that docks underneath the DNA-binding domain of KorB to latch the closed-clamp state. Our data suggest that the KorA-KorB interaction stalls KorB sliding at target core promoter elements and exploits an inherently unstable open RNA polymerase-promoter complex to exclude RNA polymerase holoenzymes from the promoters. KorA-KorB interaction also increases the residence time of KorAB on DNA, enhancing repression. Overall, we demonstrate how the clamp-locking protein KorA allows KorB to switch functions between sliding and stalling on DNA to act as an effective repressor. Our findings, therefore, provide unanticipated insights into long-range transcriptional repression mechanisms in bacteria.

## RESULTS

### KorB is a DNA-stimulated CTPase

As KorB harbors a widely distributed ParB/Srx domain (Figure 1A and S1A), we sought to determine if it binds and hydrolyses CTP. We performed isothermal titration calorimetry (ITC) of KorB individually with CTP, CDP, and a non-hydrolyzable CTP analog (CTPɣS) and found that KorB binds CTP with a moderate affinity (K_D_ = 4.5 ± 0.5 μM) and ∼tenfold more tightly to CTPɣS (K_D_ = 0.4 ± 0.1 μM) (Figure S1B). KorB binding to CDP was qualitatively much weaker than to CTP, but the data precluded the estimation of a binding affinity through curve fitting (Figure S1B). To gain insight into nucleotide binding, we solved a 2.3 Å resolution co-crystal structure of KorBΔN30ΔCTD with CTPɣS (Figure 1C and Figure S1C). To facilitate crystallization, the disordered N-terminal IncC-interacting peptide (30 amino acids) and the predicted flexible C-terminal domain (CTD) were removed (Figure 1A). At this resolution, it was not possible to assign the position of CTPɣS sulfur atom, likely due to variation in its placement from one ligand to the next in the crystal, leading to an averaging of the electron density. However, by modelling CTP instead of CTPɣS into the electron density, we could locate the CTP-binding pocket and identify CTP-contacting residues (Figure 1D). We validated these residues using structure-guided mutagenesis and ITC, finding that substituting residue R117 by alanine (R117A) abolished CTP binding, while the substitution of N146 by alanine (N146A) did not (Figure 1B).

**Figure 1.**
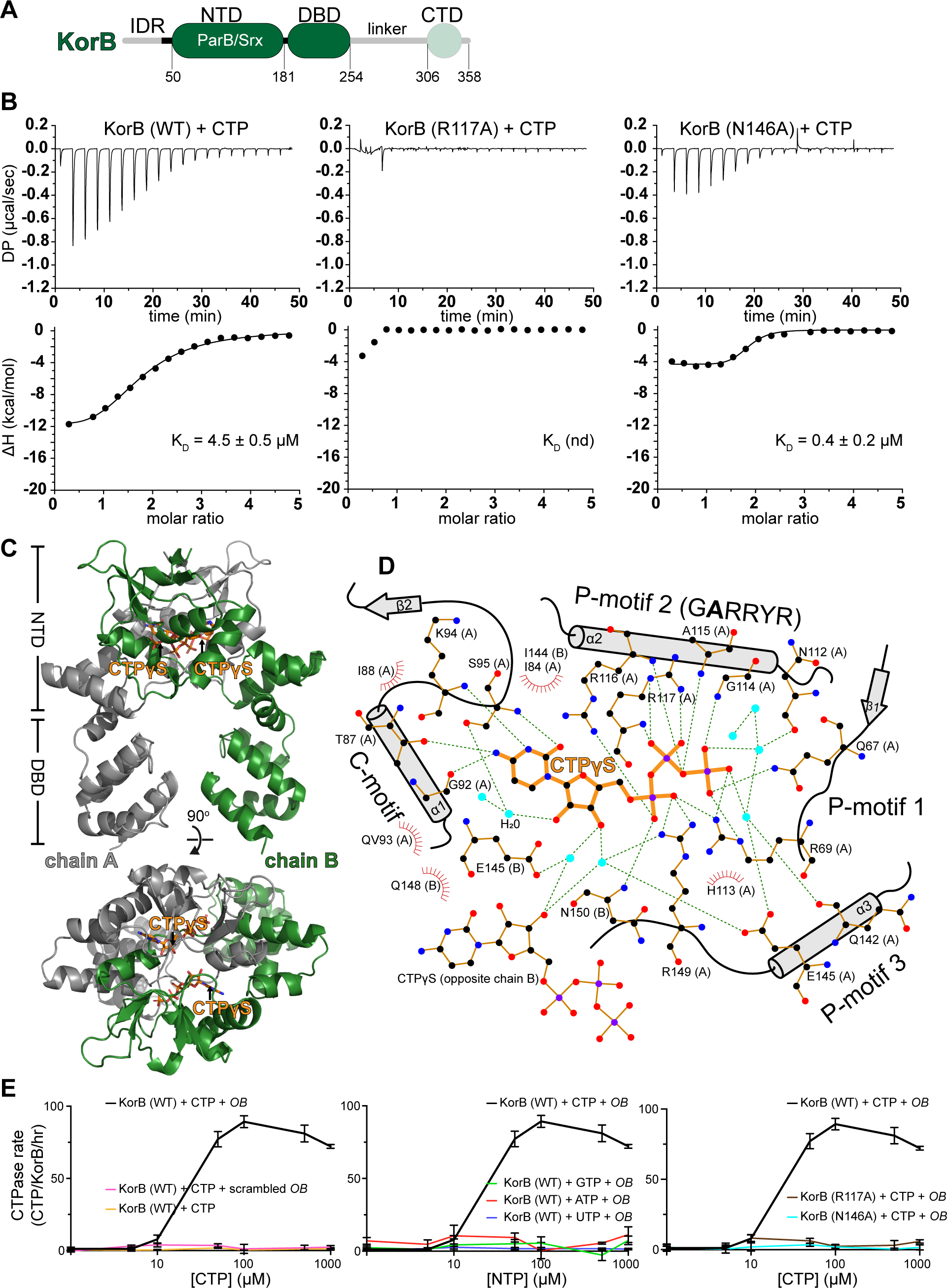
KorB binds and hydrolyses CTP in the presence of *OB* DNA. **(A)** The domain architecture of KorB: an intrinsically disordered region (IDR) IncC-interacting peptide, the N-terminal domain (NTD), a central *OB* DNA-binding domain (DBD), a predicted flexible 57-amino acid linker, and a C-terminal domain (CTD). The KorBΔN30ΔCTD variant that was used for crystallization lacks the 30 N-terminal amino acids, the linker, and the CTD (faded green). **(B)** Analysis of the interaction of KorB (WT and mutants) with CTP by ITC. Each experiment was duplicated. **(C)** Co-crystal structure of a KorBΔN30ΔCTD-CTPɣS complex reveals a CTP-binding pocket and a closed conformation at the NTD of KorB. (top panel) The front view of the co-crystal structure of KorBΔN30ΔCTD (dark green and gray) bound to a non-hydrolyzable analog CTPɣS (orange). (bottom panel) The top view of the KorBΔN30ΔCTD-CTPɣS co-crystal structure. **(D)** The protein-ligand interaction map of CTPɣS bound to KorBΔN30ΔCTD. Hydrogen bonds are shown as dashed green lines and hydrophobic interactions as red semi-circles. We did not observe electron density for Mg^2+^ in the CTP-binding pocket. **(E)** NTP hydrolysis rates of KorB (WT and variants) were measured at increasing concentrations of NTP. Experiments were triplicated, and the standard deviations (SD) of the CTPase rates were presented.

The phosphate-binding motif (GERRxR) (Figure S1A) is highly conserved in ParB, with the glutamate residue being crucial for CTPase activity but not for CTP binding^22–24^. However, KorB’s phosphate-binding motif (GARRYR) has an alanine at the equivalent position (Figure 1D and S1A), raising the possibility that KorB might not hydrolyze CTP or might do so at a slower rate than ParB. Surprisingly, KorB hydrolyzed ∼60-90 CTP molecules per KorB dimer per hour in the presence of CTP and a cognate DNA-binding site *OB* (Figure 1E and Figure S1D), a rate that is comparable to that of the *B. subtilis* ParB-like protein Noc and ∼sixfold faster than *C. crescentus* ParB (Figure S1D)^24,25^. KorB did not hydrolyze other NTPs (Figure 1E) and did not noticeably hydrolyze CTP if *OB* DNA was removed or a scrambled *OB* DNA was included (Figure 1E). Furthermore, KorB R117A was not able to hydrolyze CTP, likely owing to its inability to bind CTP (Figure 1E), while KorB N146A bound CTP but did not show detectable CTP-hydrolyzing activity (Figure 1E). Altogether, our data show that KorB is a CTPase that specifically binds and hydrolyses CTP in the presence of its cognate binding site, *OB* DNA.

### CTP and *OB* DNA promote KorB N-terminal domain engagement *in vitro*

In the presence of CTP, canonical ParB self-engages at the N-terminal domain (NTD) to create a clamp-like molecule^11,12,22,24^. In the co-crystal structure of KorBΔN30ΔCTD with CTPɣS, we similarly observed dimerization of NTDs from opposing KorB subunits (with a substantial interface area of ∼2097 Å^2^) (Figure 1C), supporting a potential clamping mechanism. To validate whether the KorB NTD mediates dimerization in solution, we tested for site-specific crosslinking of a purified KorB variant using the sulfhydryl-to-sulfhydryl crosslinker bismaleimidoethane (BMOE) (Figure 2A). Based on the KorBΔN30ΔCTD-CTPɣS co-crystal structure and sequence alignment between KorB and *C. crescentus* ParB (Figure S1A), residue S47 at the NTD was selected and substituted by cysteine on an otherwise cysteine-less KorB background (Figure S2A). This change creates a variant in which cysteines will covalently crosslink if they are within 8 Å of each other. The S47C substitution did not impact the *OB*-mediated repression function of KorB (Figure S3). In the absence of CTP, ∼20% KorB S47C could be crosslinked (lane 2), while crosslinking efficiency did not increase in the presence of 24 bp *OB* DNA or scrambled (SCR) *OB* DNA alone (lanes 3 and 4) (Figure 2A). Crosslinking efficiency reduced to ∼5-10% when CTP was included alone (lane 6) or together with scrambled *OB* DNA (lane 5) (Figure 2A), suggesting that in the absence of the *OB* DNA, CTP inhibits KorB NTD engagement. However, the crosslinking efficiency increased to ∼80% (lane 7) when both CTP and cognate *OB* DNA were added together (Figure 2A). These data suggest that both CTP and a cognate *OB* DNA are required to promote KorB NTD engagement. Consistent with this, KorB S47C R117A, which is defective for CTP-binding, did not crosslink beyond ∼20% under any tested conditions (Figure 2A). Furthermore, KorB S47C N146A, which can bind but not hydrolyze CTP, did not crosslink beyond ∼40% (Figure 2A), suggesting that the N146A substitution also impairs NTD engagement. However, this is unlikely due to the lack of CTP hydrolysis as a non-hydrolyzable analog CTPɣS could readily promote crosslinking of KorB S47C, even without DNA (Figure S2A).

**Figure 2.**
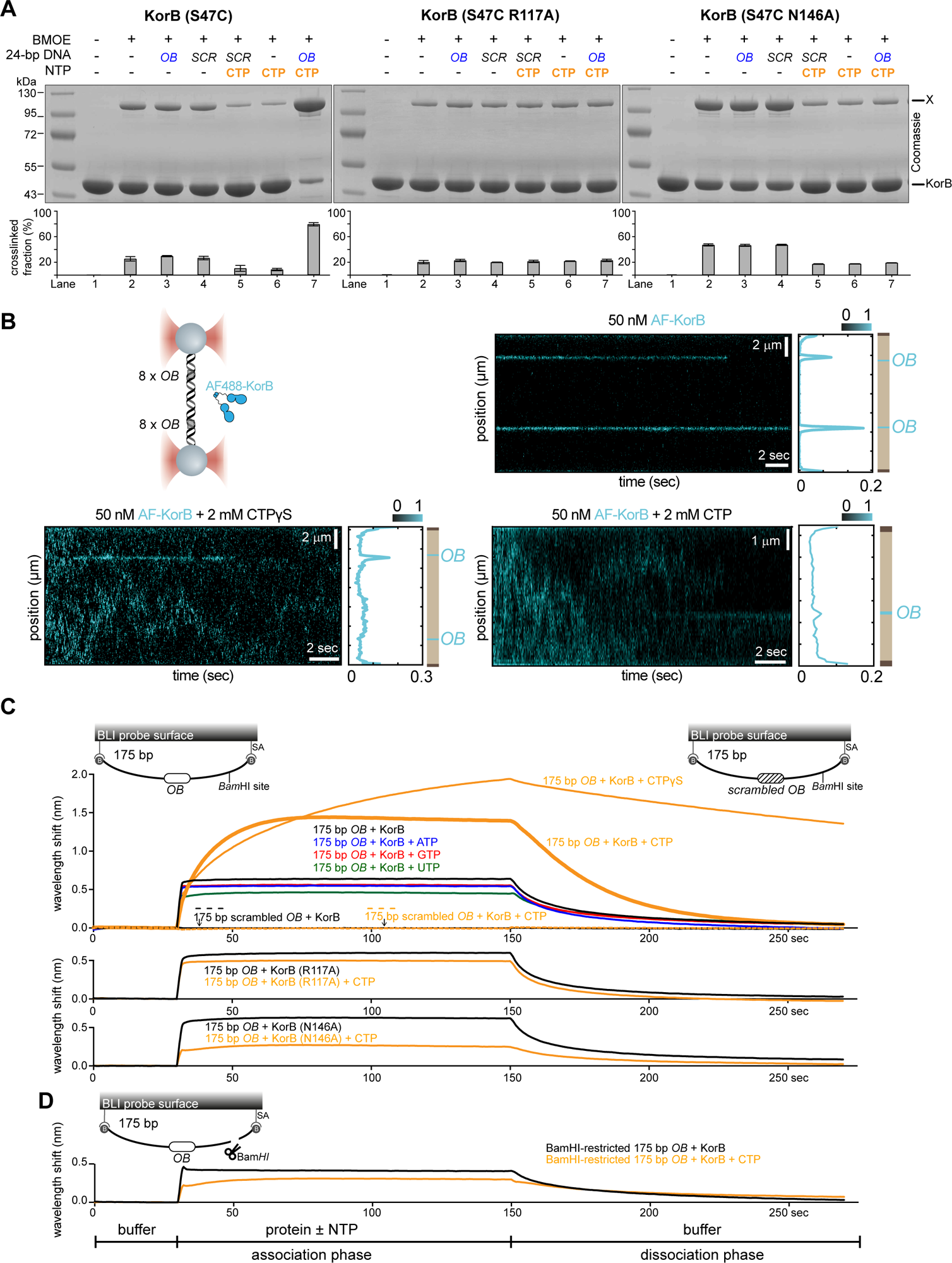
CTP and *OB* DNA promote the engagement of the N-terminal domain of KorB *in vitro*. **(A)** SDS-PAGE analysis of BMOE crosslinking products of 8 µM KorB (S47C) (and variants) ± 1 µM 24 bp *OB*/scrambled *OB* DNA ± 1 mM CTP. X indicates a crosslinked form of KorB (S47C) or (S47C R117A) or (S47C N146A). Quantification of the crosslinked fraction is shown below each representative image. Error bars represent SEM from three replicates. **(B)** CTP binding promotes the diffusion of KorB on DNA containing *OB* sites. (left panel) Schematic of the C-trap optical tweezers experiments where a 44.8-kb DNA containing two clusters of 8x *OB* sites were tethered between two beads and scanned with a confocal microscope using 488 nm illumination. A 44.8-kb DNA was constructed from two identical tandem 22.4-kb DNA, each containing 8x *OB* and 1x *OA* site (See Materials and Methods). *OA* site is omitted from the schematic picture for simplicity. (right and bottom panel) Representative kymograph showing the binding of KorB at the *OB* cluster in the presence or absence of CTPɣS or CTP. **(C)** CTP facilitates the association of KorB with a closed *OB* DNA substrate. Bio-layer interferometry (BLI) analysis of the interaction between a premix of 5 µM KorB ± 1 mM NTP and a 175-bp dual biotin-labeled DNA that contains either an *OB* or a scrambled *OB* site. Interactions between a dual biotinylated DNA and a streptavidin (SA)-coated probe created a closed DNA molecule where both ends were blocked (see also the schematic diagram of the BLI probes). Other KorB variants, KorB (R117A) and KorB (N146A) were also analyzed in the same assay. **(D)** BLI analysis of the interaction between a premix of 5 µM KorB ± 1 mM CTP and a BamHI-restricted dual biotinylated *OB* DNA. Each experiment was triplicated, and a representative sensorgram was shown.

Lastly, to investigate dimerization of the KorB CTD, we created a KorB K351C variant. This position is predicted to be crosslinkable based on symmetry-related interactions observed in the previously published crystal structure of KorB CTD (PDB: 1IGQ)^26^. KorB K351 showed a high crosslinking efficiency at ∼40% in most tested conditions (Figure S2A), consistent with the known role of the CTD in dimerization. Moreover, the presence of both CTP and *OB* DNA further increased the crosslinked fraction of KorB K351C to ∼60% (lane 7) (Figure S2A), suggesting that CTP and cognate DNA together promote KorB dimerization at both the NTD and CTD.

### CTP enables KorB diffusion and accumulation on a closed *OB* DNA substrate

To investigate the impact of CTP binding on KorB-DNA interaction, we employed dual optical tweezers combined with confocal fluorescence microscopy^27^. Here, individual 44.8-kb DNA molecules were immobilized between two polystyrene beads and extended almost to their contour length under force (Figure 2B). To amplify the signal of KorB DNA binding, eight *OB* sites were present either in one or two clusters on the DNA (Figure 2B). We first incubated DNA with 50 nM of AlexaFluor (AF) 488-labeled KorB alone and took confocal images over time to construct kymographs. A 30s kymograph showed two stable regions of high fluorescence, corresponding to the position of the two *OB* clusters (Figure 2B), consistent with the initial binding of KorB at *OB* being CTP-independent. In the presence of 2 mM CTPɣS or CTP, fluorescence signals from AF488-KorB were identified outside of the *OB* clusters (Figure 2B), suggesting that KorB-CTP was now distributed non-specifically along the DNA. Next, we measured the position of individual AF488-KorB molecules along the DNA over time to determine a diffusion constant (*D*) for KorB of ∼1.61 ± 0.12 µm^2^/s (Figure S2B), which is ∼fourfold higher than the value previously determined for *B. subtilis* ParB-CTP diffusion on DNA^27^.

To further investigate the roles of KorB NTD engagement, we followed the interaction of KorB with cognate *OB* DNA in real-time using bio-layer interferometry (BLI). We employed a 175-bp dual biotin-labeled *OB* or scrambled *OB* DNA tethered at both ends to a streptavidin-coated probe to form a closed DNA loop (Figure 2C). BLI monitors the shift in wavelength resulting from changes in probe optical thickness during the association and dissociation of KorB from the DNA loop. In the presence of 5 µM purified KorB alone, a low BLI signal was observed (Figure 2C), consistent with *OB*-specific initial binding by KorB. Premixing KorB with ATP, GTP, and UTP did not change the BLI signal markedly (Figure 2C). However, premixing with 1 mM CTP increased the BLI response ∼3-4 fold (Figure 2C), suggesting that KorB accumulates on DNA at greater levels than an initial binding event at *OB* under these conditions. Consistent with the requirements for CTP and NTD engagement, KorB R117A and N146A did not accumulate on DNA beyond initial loading at *OB* (Figure 2C). Next, we observed that non-hydrolyzable CTPɣS increased the BLI signal even further, and reduced KorB dissociation from the DNA loop ∼10 fold compared to CTP (see the dissociation phase of the BLI experiments, k_off_ = 3.5×10^-3^ ± 10^-4^ sec^-1^ vs. 3×10^-2^ ± 10^-4^ sec^-1^; Figure 2C). This result indicates that CTP hydrolysis is not required for KorB accumulation on an *OB* DNA loop but might facilitate its release from DNA.

Next, we investigated whether a DNA substrate with a free end could support high KorB association (Figure 2D). The 175-bp dual biotin-labeled DNA loop was designed with a single BamHI recognition site flanking the *OB* site. The DNA-loop probe was immersed into BamHI-containing buffer to enable DNA cutting to generate a free end. The addition of CTP no longer resulted in a high BLI signal (Figure 2D), suggesting that KorB spreads by diffusion and slides off the free DNA end.

### KorB-CTP does not condense *OB* DNA *in vitro*

Canonical ParB was previously reported to bridge and condense DNA *in vitro* in the presence of CTP^27–30^. Since the optical trap approach used above keeps DNA extended, we employed magnetic tweezers, which allow simultaneous measurement of multiple DNA molecules and application of lower forces, to test whether KorB induces DNA condensation (Figure S2C). Single DNA molecules containing a cluster of 16x *OB* sites were immobilized between a glass surface and magnetic beads (Figure S2C). Forces in 1-5 pN range were applied to the beads to stretch *OB* DNA between a pair of magnets before the addition of either buffer alone or with different concentrations of KorB in the presence of 2 mM CTP. As a control, the same experiment was performed using *Bacillus subtilis* ParB (*Bs*ParB) in the presence of 2 mM CTP. The force was then gradually lowered to 0.002 pN, which would be permissive for DNA condensation, and the extension of the tethered DNA was monitored in real-time (Figure S2C). We observed no difference in the force-extension curves in the presence or absence of KorB-CTP (Figure S2C), indicating that KorB-CTP does not condense DNA under the tested conditions. In the case of *Bs*ParB, we observed unspecific condensation at high concentrations (1-2 µM). As expected, *Bs*ParB was not able to condense at a lower concentration (500 nM) because *Bs*ParB does not recognize *OB* sites on our DNA substrate.

### CTP-dependent KorB NTD-engagement is essential for long-range transcriptional repression

KorB was previously shown to mediate long-range transcription repression^13^. To investigate whether CTP-binding and DNA-sliding influence this activity, we constructed two promoter-*xylE* transcriptional fusion reporters and assayed for catechol dioxygenase activity in the presence or absence of KorB *in vivo* (Figure 3A). In these constructs, *OB* DNA was either engineered 4 bp (*OB*-proximal) or 1.5 kb (*OB*-distal) upstream of the -35 element of the core promoter of the RK2 *trbB* gene (Figure 3A). The presence of KorB led to ∼4-5 fold transcriptional repression compared to a KorB-minus control for both proximal and distal promoter-reporter constructs, consistent with previous findings^13^ (Figure 3B). The KorB R117A and N146A variants, which can site-specifically load onto *OB* in the absence of CTP but cannot slide, are capable of repressing transcription from an *OB*-proximal promoter but were defective at repressing an *OB*-distal promoter (Figure 3B). Overall, our data suggest that the ability of KorB to bind CTP, close the clamp and slide on DNA is crucial for long-range gene silencing.

**Figure 3.**
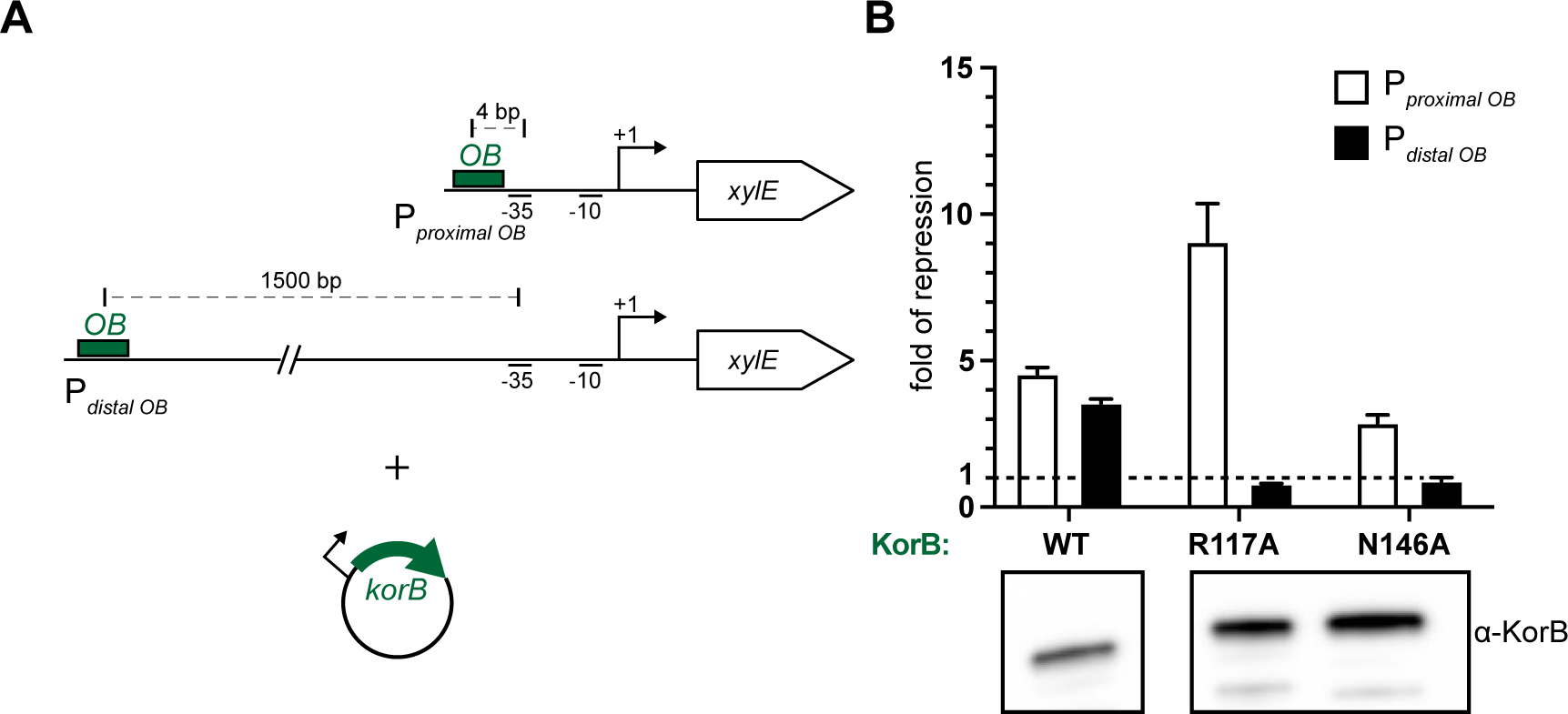
CTP-dependent N-engagement is essential for KorB to repress transcription from a distance. **(A)** Schematic diagrams of promoter-*xylE* reporter constructs. For the *OB*-proximal promoter, *OB* is positioned 4 bp upstream of the -35 core promoter element. For the *OB*-distal promoter, *OB* is 1500 bp upstream of the -35 element. *E. coli* cells were co-transformed with the reporter plasmid and an inducible plasmid expressing KorB (WT/variants or empty plasmid as a negative control). **(B)** Substitutions R117A and N146A at the NTD of KorB did not drastically affect transcriptional repression at an *OB*-proximal promoter but eliminated repression at an *OB*-distal promoter. Values shown are fold of repression, a ratio of XylE activities from cells co-harboring a reporter plasmid and a KorB-expressing plasmid to that of cells co-harboring a reporter plasmid and an empty plasmid (KorB-minus control). Experiments were triplicated, and the SDs were presented. An α-KorB immunoblot from lysates of cells used in the experiments is also shown below.

### KorA promotes the NTD-engagement of KorB independent of CTP and *OB* DNA *in vitro*

KorB has previously been reported to require KorA (Figure 4A) to strengthen transcriptional repression, especially at *OB*-distal promoters^13,19,20^. However, it remains unclear how KorA and KorB cooperate to regulate gene expression. We hypothesized that KorA might modulate the CTP-dependent activities of KorB. To investigate this possibility, we used a crosslinking approach (Figure 4B) in which purified cysteine-less KorA was pre-incubated with KorB S47C in the presence or absence of *OB* DNA and/or CTP before crosslinking by BMOE. KorB S47C crosslinked at an efficiency of ∼20% in the absence of KorA (lane 2) (Figure 4B). In the presence of KorA, the crosslinking efficiency of KorB S47C increased to ∼60%, even in the absence of CTP and *OB* DNA (lane 3) (Figure 4B) or when *OB* or scrambled *OB* DNA alone was included (lanes 4 and 5) (Figure 4B). In the presence of CTP alone, KorA increased the crosslinking efficiency of KorB S47C to ∼20% (lane 7) (Figure 4B), again higher than the ∼10% crosslinked fraction observed in the absence of KorA (lane 6) (Figure 4B). KorB S47C crosslinked maximally at ∼80% efficiency when KorA, CTP, and the cognate *OB* DNA were all included (lane 9) (Figure 4B). Notably, for both KorB R117A and N146A variants, which are defective in NTD-engagement, the presence of KorA further increases the efficiency of crosslinking in all tested conditions (Figure S4A vs. Figure 2A). Overall, our results suggest that KorA promotes engagement of the KorB NTD. We reasoned that KorA likely does so via a direct interaction with KorB^18,19^, which we confirmed by ITC (K_D_ of KorAB interaction = 4.2 ± 0.5 µM) (Figure 4C).

**Figure 4.**
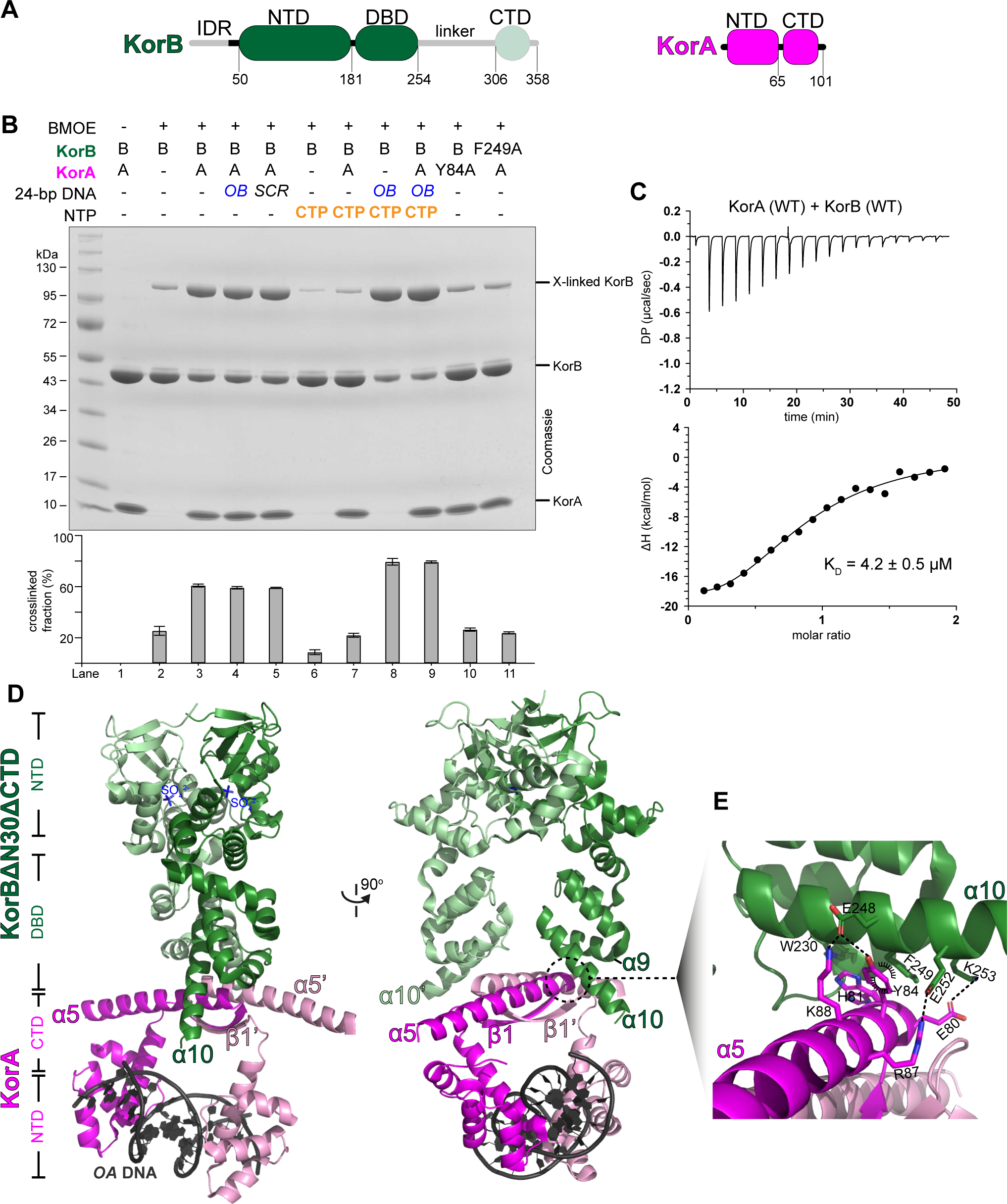
KorA promotes the N-engagement of KorB independent of CTP and *OB* DNA *in vitro*. **(A)** The domain architecture of KorB (same as Figure 1A) and KorA. The KorBΔN30ΔCTD variant that was used for crystallization lacks the 30 N-terminal amino acids, the linker, and the CTD (faded green). KorA has an N-terminal *OA* DNA-binding domain (NTD) and a C-terminal dimerization domain (CTD). **(B)** SDS-PAGE analysis of BMOE crosslinking products of 8 µM KorB (S47C) (and the variant S47C F249A) ± KorA (WT or the variant Y84A) ± 1 mM 24 bp *OB*/scrambled *OB* DNA ± 1 mM CTP. X indicates a crosslinked form of KorB. Quantification of the crosslinked fraction is shown below each representative image. Error bars represent SD from three replicates. **(C)** Analysis of the interaction of KorB with KorA by ITC. Each experiment was duplicated. **(D)** KorA-DNA docks onto the DNA-binding domain of a clamp-closed KorB in the co-crystal structure of a KorBΔN30ΔCTD-KorA-DNA complex. (left panel) The side view of the co-crystal structure of KorBΔN30ΔCTD (dark green and light green) bound to a KorA (magenta and pink) on a 14-bp *OA* DNA duplex (black). (right panel) The front view of the KorBΔN30ΔCTD-KorA-DNA co-crystal structure. **(E)** Helix α10 (dark green) of KorB interacts with helix α5 (magenta) of KorA. Hydrogen bonds are shown as dashed lines, pi-stacking interaction between the aromatic ring of Tyr84 in KorA with the Glu248-Phe249 peptide bond of KorB and between the aromatic ring of Phe249 in KorB with the Glu80-His81 peptide bond of KorB are shown as a black semi-circles (see also Figure S5C).

### KorA-DNA docks onto the DNA-binding domain locking the KorB-clamp

To determine the mechanism by which KorA induces the KorB closed-clamp conformation, we solved a 2.7 Å resolution co-crystal structure of a tripartite complex between KorBΔN30ΔCTD and KorA bound to its 14-bp cognate *OA* (operator of KorA) DNA duplex (Figure 4D and Figure S5A). The asymmetric unit contains two subunits of KorA that dimerize via the C-terminal β1 strands and α5 helices (Figure 4D). The DNA-binding domains (helices α1-4) from opposing KorA subunits dock into adjacent major grooves on the 14-bp *OA* DNA duplex (Figure 4D). The asymmetric unit also contains two subunits of KorBΔN30ΔCTD whose opposing NTDs (strands β1-3 and helices α1-4) are self-dimerizing (with an interface area of ∼1250 Å^2^) (Figure 4D). Structural alignment of the tripartite complex vs. the CTPɣS-bound KorBΔN30ΔCTD complex showed a low root mean square displacement value (RMSD), suggesting that KorBΔN30ΔCTD in the tripartite complex has already adopted an NTD-engaged conformation, even though CTPɣS was not added to the crystallization setup. A closer inspection of KorB NTD in the tripartite complex revealed that helices α3 and α4 from each subunit bundle together (bundling-in conformation) (Figure S6A), in contrast to the CTPɣS-bound KorBΔN30ΔCTD structure in which helix α3 swings outwards to pack against α4’ from the opposing subunit (swinging-out conformation) (Figure S6A). The reciprocal exchange of helices in the swinging-out conformation maintained packing at the α3-α4 protein core of the KorB NTD and was likely driven by CTPɣS-binding (Figure S6A). The bundling-in conformation of α3-4 is often associated with the NTD-disengaged open-clamp conformation of ParB/ParB-like proteins^24,25,31–33^, while the swinging-out conformation is often observed for a nucleotide-bound NTD-engaged closed-clamp conformation^11,22,24^. For canonical ParB, the closed-clamp conformation was demonstrated to be energetically favorable but the transition to the closed-clamp state is slow without CTP and cognate DNA^11,23^. We therefore reasoned that KorA-DNA may facilitate this transition or capture KorBΔN30ΔCTD in the NTD-engaged state, as observed in the tripartite structure.

The two opposing *OB* DNA-binding domains (DBD) of KorB are also closer together when bound to KorA (inter-domain distance = ∼30 Å) than when they are bound to *OB* DNA alone (inter-domain distance = ∼34 Å; PDB: 1R71)^34^ (Figure S6B), which is incompatible with specific binding to the *OB* site and suggests that KorA binding stabilizes a closed-clamp conformation that is more compatible with sliding on DNA. The DNA-bound KorA dimer docks underneath the *OB*-binding domain of KorBΔN30ΔCTD, specifically via helix α5 of KorA interacting directly and at a perpendicular angle to helix α10 of KorBΔN30ΔCTD (Figure 4D). A network of specific interactions is established at the KorA-KorB interface, including KorA Y84 and KorB F249 (Figure 4E and Figure S5B-C). To verify the observed interactions, KorA Y84A^18^ and KorB F249A variants were constructed and assayed for binding to wild type KorB or KorA, respectively. ITC data showed that these single mutations completely abolished the interaction between KorA and KorB (Figure S4B). Furthermore, in our crosslinking assay, neither KorA Y84A nor KorB S47C F249A resulted in a higher fraction of crosslinked KorB beyond ∼20% (lanes 10 and 11 vs. lane 3, Figure 4B). Overall, our results showed that KorA, via its C-terminal helix α5, directly interacts with the DBD of KorB to lock KorB in an NTD-engaged closed-clamp state.

### KorA blocks KorB diffusion on DNA and increases retention time at operator *OA*

To investigate the significance of KorAB interaction for DNA-sliding in the presence of CTP, we used fluorescently labeled KorAB in our *in vitro* optical tweezers setup. Individual DNA molecules were engineered to contain an *OA* site ∼3 kb away from the 8x *OB* sites. These molecules were immobilized between two polystyrene beads and extended to almost their contour length under force (Figure 5A). In the absence of CTP, kymographs showed that fluorescence signals from AF488-labeled KorB and AF647-labeled KorA were confined only to the corresponding positions of *OB* and *OA* sites, respectively (Figure 5A left panel). When CTP was included, the location of the fluorescence signal from AF647-KorA remained confined to *OA* sites. In contrast, AF488-KorB signals were found outside of the *OB* clusters, consistent with KorB-CTP sliding on DNA (Figure 5A right panel). However, we observed the accumulation of AF488-KorB fluorescence signal between *OB* and its proximal *OA* site (Figure 5A right panel), suggesting that *OA*-bound KorA and/or *OA*-bound KorA-KorB complex might block sliding of KorB-CTP.

**Figure 5.**
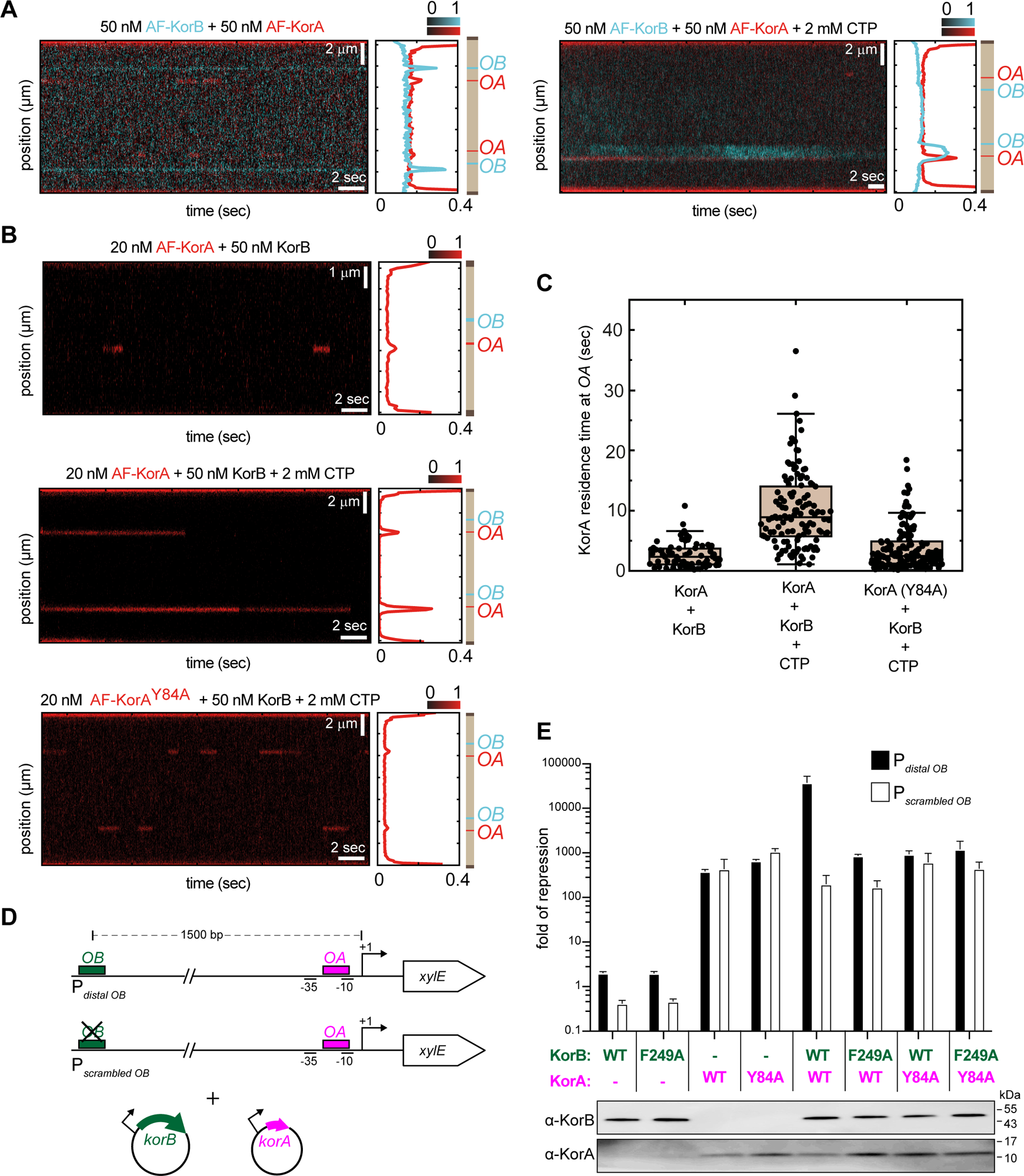
KorA can block the diffusion of KorB on DNA, and KorB increases the retention time of KorA at its operator *OA*. **(A)** KorA acts as a roadblock to KorB diffusion along the DNA to generate an accumulation of KorB between the *OA* and *OB* sites. (left panel) Representative kymograph showing the binding of AF488-labeled KorB and AF647-labeled KorA in the absence of CTP. (right panel) Representative kymograph showing the binding of AF488-labeled KorB and AF647-KorA in the presence of 2 mM CTP. The positions of the 8x *OB* and 1x *OA* clusters in the cartoon are represented to scale. There are two clusters of *OA* and *OB* in some examples, see Materials and Methods section for details on the construction of DNA substrates. All the fluorescence intensities were normalized and the scale in the intensity profiles were adjusted for better visualization. **(B)** KorB increases the retention time of KorA at *OA*. (top panel) Representative kymograph showing the binding of AF647-labeled KorA in the presence of unlabeled KorB. (middle panel) Representative kymograph showing the binding of AF647-KorA in the presence of unlabeled KorB and 2 mM CTP. (bottom panel). Representative kymograph showing the binding of AF647-KorA Y84A in the presence of unlabeled KorB and 2 mM CTP. **(C)** Box plot showing the retention times of AF647-KorA in the presence of KorB (2.61 ± 0.24 s, mean ± SEM, n= 70) or KorB-CTP (10.11 ± 0.56 s, mean ± SEM, n= 125), and the retention time of KorA (Y84A) variant in the presence of KorB-CTP (3.72 ± 0.29 s, mean ± SEM, n = 148). All the fluorescence intensities were normalized and the scale in the intensity profiles were adjusted for better visualization. **(D-E)** KorA captures the KorB-CTP clamp to heighten long-range transcriptional repression. (left panel) Schematic diagrams of promoter-*xylE* reporter constructs. For *OB*-distal promoter, *OB* is 1500 bp upstream of the -35 core promoter element and *OA* is overlapping the -10 element. *OB* site was scrambled in the scrambled-*OB* distal promoter-*xylE* reporter construct. *E. coli* cells were co-transformed with the reporter plasmid, an inducible plasmid expressing KorB (WT or variants or empty vector as a negative control), and an inducible plasmid expressing KorA (WT or variants or empty vector as a negative control). (right panel) Substitutions Y84A on KorA, F249A on KorB, or removing the upstream *OB* site eliminated high-level transcriptional repression. Values shown are fold of repression, a ratio of XylE activities from cells co-harboring a reporter plasmid and KorB/A-expressing plasmid to that of cells co-harboring a reporter plasmid and an empty plasmid (KorA/B-minus control). Experiments were triplicated, and the SDs were presented. An α-KorB and α-KorA immunoblots from lysates of cells used in the experiments are also shown below.

The AF647-KorA fluorescence signal exhibited an on-off behavior at the *OA* site in the presence of KorB but absence of CTP (Figure 5B). However, the AF647 signal (at *OA*) was more stable over time when KorB and CTP were both included (Figure 5B), suggesting that KorB-CTP might reduce KorA dissociation from *OA*. To test this possibility, we quantified the retention time of AF647-KorA at *OA* in the presence of unlabeled KorB with or without 2 mM CTP (Figure 5B). The retention time increased ∼4.5 fold in the presence of KorB-CTP compared to apo-KorB (Figure 5C). Notably, this effect was abolished when a non-interacting AF488-KorA Y84A variant was used instead (Figure 5C). Altogether, we suggest that KorA can block the diffusion of KorB on DNA, and KorB increases the retention time of KorA on DNA.

### KorA captures and locks KorB-CTP clamp converting KorB to a local co-repressor

To investigate the interplay between KorA and KorB-CTP *in vivo*, we engineered a promoter-*xylE* reporter where the *OB* site is 1.5 kb upstream of the core promoter while the *OA* site overlaps with the -10 element (Figure 5D). The positioning of *OA* and *OB* DNA mimicked KorAB-dependent promoters natively found on the RK2 plasmid. Note that *OA* is invariably found in immediate proximity to the core promoter elements of RK2 genes^6^. Inducing the production of KorB repressed transcription weakly by ∼fourfold, as previously observed (Figure 5E). This low degree of repression was also observed when KorB F249A, which does not bind KorA, was expressed alone. Both wild type KorA and the non-interacting KorA Y84A variant repressed transcription ∼200 fold when produced alone, consistent with previous findings^18^. However, when wild type KorB and KorA were co-expressed, transcriptional repression was increased to ∼38,000 fold, indicating cooperation between the two proteins^18^. This cooperation in transcriptional repression was abolished when wild type KorA and KorB F249A or KorA Y84A and wild type KorB were co-produced. Since the F249A and Y84A substitutions removed the ability of KorA to bind KorB and close the clamp, we suggest that these activities are essential for effective and efficient transcriptional repression^18^.

Since KorA could bind apo-KorB (Figure 4C), we wondered whether DNA-unbound KorB could cooperate with KorA to elicit the same high level of transcriptional repression *in vivo*. To investigate, we scrambled the distal *OB* site on the promoter-*xylE* reporter construct and measured XylE activity when wild type KorAB or their variants were co-produced (Figure 5D). In the absence of *OB*, we reasoned that only apo-KorB and CTP-bound KorB (i.e., opened-clamp states) could form inside the cells, while *OB*-stimulated DNA-entrapped KorB-CTP (i.e., closed-clamp) could not. As expected, in the absence of *OB*, no cooperative transcriptional repression was observed with any combination of KorAB or their variants (Figure 5E). Collectively, these results suggest that *OA*-bound KorA captures a DNA-entrapped sliding clamp of KorB to cooperatively repress promoters and explains the long-range transcription repression activity of KorB.

### KorB-CTP and KorA exploit unstable promoter complexes to repress transcription initiation

The molecular mechanism governing the co-repressive activity of KorAB remains elusive. The proximity of the *OA* site to the core promoter elements of RK2 genes^35,36^ suggests a classical RNAP-occlusion mechanism. However, previous work suggested a potential interaction between KorB and RNAP, indicating a repression mechanism stabilizing a closed or intermediate state of a promoter complex where the DNA bubble is incompletely formed^37^. To investigate further, we first employed *in vitro* transcription initiation assays with *E. coli* RNA polymerase (*Eco* RNAP) and a linear DNA containing a model promoter *PkorABF* (referred to as *PkorA*)^38^. *PkorA* controls the transcription of the central operons coding for *korA*, *incC*, and *korB*. *PkorA* features an *OA* site overlapping with the upstream half of its -10 element, and an *OB* site is situated directly upstream of the -35 element (Figure 6A). Both the -10 and -35 promoter elements play a crucial role in RNAP holoenzyme (Eσ^70^) promoter-dependent initiation^39^. Transcription initiation assays confirmed that KorA and KorB (with and without CTP) individually function as repressors (Figure S7A and B), while their combined activity is further augmented, consistent with our *in vivo* and single-molecule studies (Figure 5C and E). Notably, KorA emerges as the stronger repressor (*p ≤ 0.05*; unpaired Welch’s t-test; Figure S7B), aligning with prior studies and the finding that its operator site is embedded within the -10 element^40,41^. Next, to investigate the possible binding of KorA and KorB to *E. coli* Eσ^70^-promoter complexes (Eσ^70^:DNA), we used native mass spectrometry (nMS). Introducing a 2.5-fold excess of KorA or KorB to *PkorA*-bound *E. coli* Eσ^70^ (Eσ^70^:DNA) led to near complete dissociation of Eσ^70^ from DNA rather than ternary complex formation (Figure 6B and C). Minor peaks for Eσ^70^ + 2KorA and Eσ^70^:DNA + 4KorA were detected in the samples with KorA but no corresponding KorB-bound Eσ^70^ peaks were observed (Figure 6B and C) indicating low-level binding of KorA to *PkorA*-bound Eσ^70^ compared to KorB.

**Figure 6.**
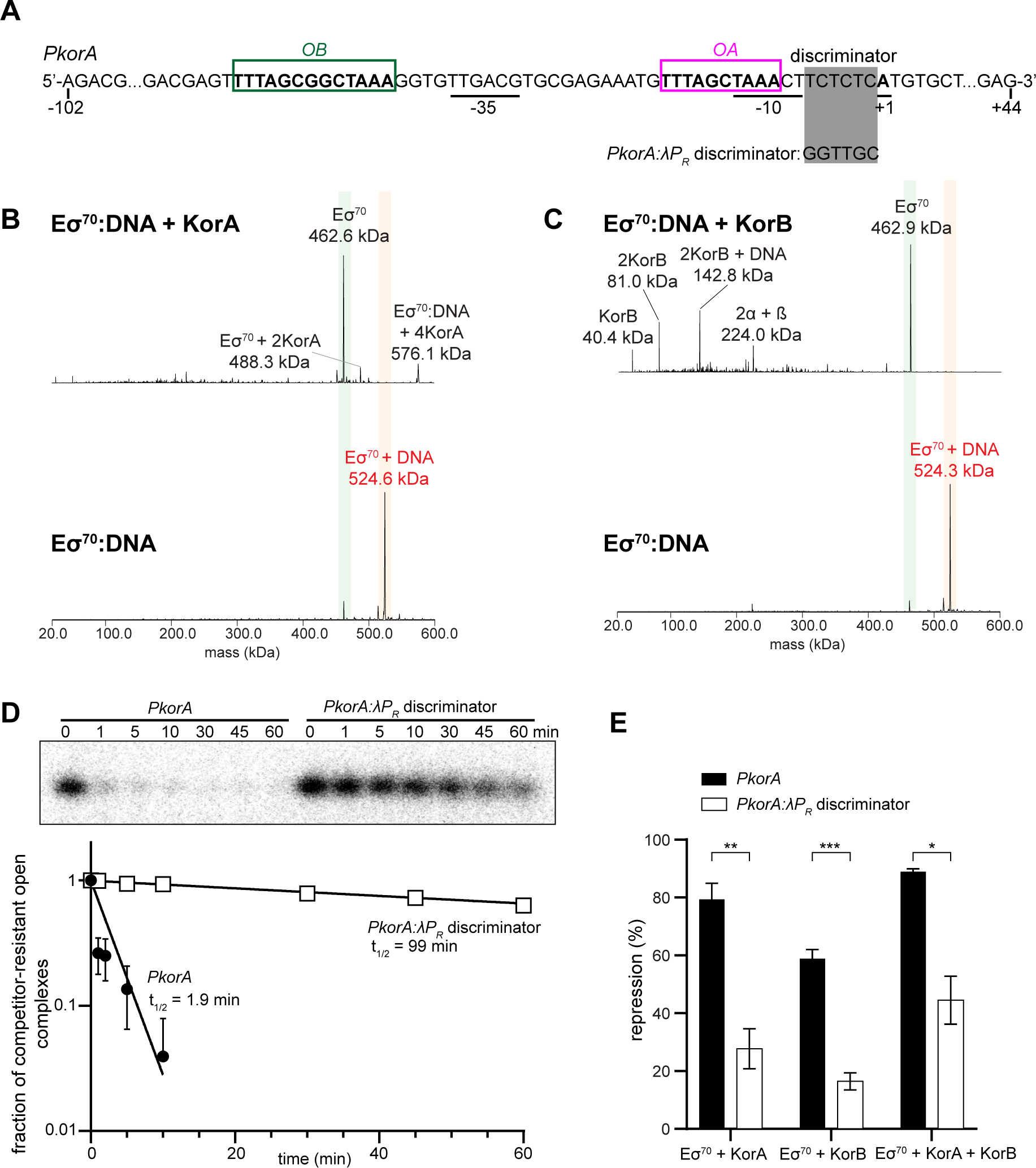
KorA and KorB exploit kinetically unstable *E. coli* RNAP:*PkorA* DNA complexes to repress transcription initiation. **(A)** Promoter scaffold from the RK2 *korABF* operon (*PkorA*) is shown with core promoter elements (underlined), *OA* (magenta box), and *OB* (green box). Differences to the *PkorA*:λ*P_R_* discriminator are depicted. **(B)** Deconvolved nMS spectra of *E. coli* Eσ^70^ holoenzyme assembled on 100-bp *PkorA* DNA (Eσ^70^:DNA) with and without 2.5-fold excess KorA dimer in 150 mM ammonium acetate pH 7.5, 0.01% Tween-20. **(C)** Deconvolved nMS spectra of Eσ^70^:DNA with and without 2.5-fold excess KorB dimer in 500 mM ammonium acetate pH 7.5, 0.01% Tween-20. **(D)** (top panel) Representative gel closeup on the abortive RNA product (5’-ApUpG-3’) transcribed by Eσ^70^ in *in vitro* abortive initiation half-life assays on the two *PkorA* linear scaffold variants. (bottom panel) Plot of fraction of competitor-resistant open complexes (from normalized abortive RNA band intensities) against time. Data points from three experimental replicates are mean values ± SEM with an exponential trendline fit. Some error bars are too small or lead to negative values, thus were omitted. Estimated half-lives are shown adjacent to exponential decay trendlines. **(E)** WT *PkorA* and *PkorA*:λ*P_R_* discriminator *in vitro* transcription repression of Eσ^70^ in the presence of 5-fold excess KorA and/or KorB. Data points from three experimental replicates are normalized to holoenzyme only control as mean values ± SEM. P-values were calculated by unpaired Welch’s t-tests; * p ≤ 0.05, ** p ≤ 0.01, *** p ≤ 0.001.

Analysis of the *PkorA* promoter sequence revealed near-consensus -10 (TAAACT; consensus is TATAAT) and -35 elements (TTGACG; consensus is TTGACA)^39,42^, an AT-rich region upstream of the -35 element (consistent with an UP element)^43,44^, and a consensus 17-bp spacer between the - 10 and -35 elements (Figure 6A). The presence of these features highly favors promoter melting by RNAP^45,46^. However, the discriminator (the sequence between the -10 element and the transcription start site) (Figure 6A), crucial for the stability of an open RNAP-promoter complex (RPo), is highly unfavorable (TCTCTC; consensus is GGGnnn)^47^. To examine the importance of the discriminator sequence in KorAB-mediated regulation of *PkorA*, we substituted the wild-type discriminator with that of bacteriophage λ *P_R_* (Figure 6A), a strong promoter forming kinetically stable complexes with a half-life (*t_1/2_*) of many hours due to a favored discriminator^48,49^. The substitution increased the half-life of *PkorA* RPo from two minutes to over 90 minutes, stabilizing the RPo over 50-fold (Figure 6D). The substituted discriminator showed a significant decrease in *in vitro* transcription repression when KorA and KorB were added individually or combined (Figure 6E). The instability of the *PkorA* RPo complex, coupled with the observations of KorA and KorB binding to DNA but not to Eσ^70^, suggests the following model. *PkorA* RPo is inherently unstable, resulting in frequent RNAP dissociation that allows KorA and KorB to bind their respective operator sites and, upon sliding, to form a repressome on the promoter that occludes RNAP from the promoter. Our findings suggest that promoters leading to unstable RPo are more susceptible to inhibition by competitive occlusion.

## DISCUSSION

### CTP and KorA regulate a KorB functional switch

KorB is essential for segregation of the RK2 plasmid and for regulating gene expression for all basic RK2 functions, including replication and conjugative transfer^5^. Both functions are necessary to ensure stable vertical inheritance and horizontal transmission of this broad-host-range multi-drug resistance plasmid. Our work has shown that CTP is required for KorB to form an *OB*-dependent sliding clamp on DNA, which enables KorB to travel a long distance to repress distal promoters (Figure 7). Similar to chromosomal ParB^7,50^, we propose that KorB-CTP loads at the *OB* site and then switches to a closed-clamp conformation through NTD engagement, allowing it to escape the high-affinity loading site and slide away while entrapping DNA (Figure 7A). Our co-crystal structures of a closed-clamp KorB revealed a DBD incompatible with *OB*-binding (Figure S6B), lending support to our proposal that a closed-clamp KorB can escape the *OB* site. Once KorB-CTP slides away, another KorB-CTP molecule can bind a now-free *OB* and repeat the process, enabling multiple KorB molecules to decorate the DNA surrounding *OB*. Similar to the ParAB*S* system, it is most likely that a high local concentration of KorB is also required to activate the ATPase activity of a ParA homolog IncC, which then segregates the KorB-DNA complex^3,21,29,31,51–57^.

**Figure 7.**
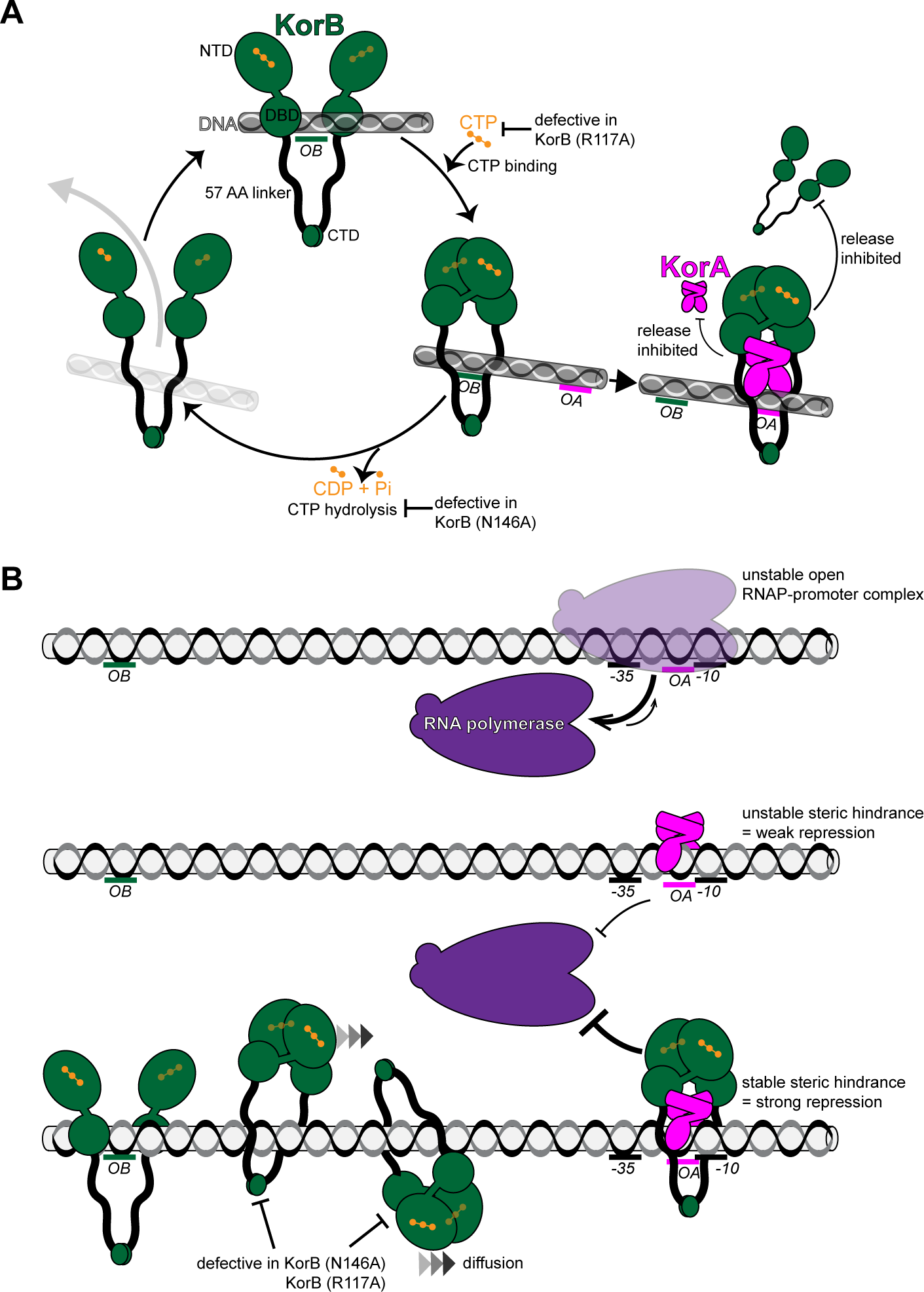
A proposed model for the CTP-dependent clamp-sliding activity of KorB, clamp-locking activity of KorA, and their cooperation to enable tight transcription repression from a distance. **(A)** A model for KorB (dark green) loading, sliding, and release cycle. Loading KorB is likely an open clamp, in which *OB* DNA binds at the DBD. The presence of CTP (orange) and *OB* DNA likely triggers KorB clamp closing. In this state, KorB can slide away from the *OB* site by diffusion while entrapping DNA. CTP hydrolysis and/or the release of hydrolytic products (CDP and inorganic phosphate Pi) likely reopen the clamp to release DNA. Substitutions that affect key steps in the CTP binding/hydrolysis cycle are also indicated on the schematic diagram. KorA (magenta) bound at the *OA* site can form a complex with and promote or trap KorB in a closed clamp state. In this state, KorB is most likely still entraps DNA. The formation of the tripartite KorAB-DNA reduces the release of KorB from DNA as well as the release of KorA from *OA* DNA. **(B)** A model for KorA-KorB cooperation to heighten long-range transcription repression. On the RK2 plasmid, the *OB* site can position kb away from the target core promoter elements (-10 -35) while *OA* is almost invariably near to these core promoter elements. Owing to the presence of an unfavorable discriminator sequence, the RNAP holoenzyme-promoter DNA complex is inherently unstable. In the absence of KorB-CTP, KorA binds *OA* with a low retention time, thus only providing an unstable steric hindrance to occlude RNA polymerase (magenta) from the core promoter elements, resulting in weak transcriptional repression. In the presence of CTP, KorB-CTP loads at a distal *OB* site, closes the clamp and slides by diffusion to reach the distal *OA* site. *OA*-bound KorA captures and locks KorB in a clamp-closed conformation. In this state, the KorAB-DNA co-complex presents (i) a larger steric hindrance and (ii) a more stable steric hindrance because the release of KorB from DNA as well as the release of KorA from *OA* DNA are now reduced. As a result, the KorAB-DNA co-complex can exploit the unstable RNAP-promoter complex and occludes RNA polymerase more effectively, hence stronger transcriptional repression than each protein alone can provide.

While a sliding clamp is an elegant evolutionary solution to ensure faithful plasmid segregation, it is seemingly at odds with transcription repression. Such a clamp could, for example, slide past the core promoter region, providing access to RNAP. Accordingly, our *in vivo* transcriptional reporter assay showed that KorB alone, even at high expression levels, only repressed the proximal and distal promoters weakly (Figure 3). It has also been shown that transcribing RNA polymerases can steadily traverse the *B. subtilis* ParB-DNA partition complex *in vitro*^30^ and ParB sliding to neighboring DNA does not affect gene expression *in vivo*^58^. However, there are exceptions; for examples, P1 ParB autoregulates its expression^59–61^ or *Pseudomonas aeruginosa*, *Vibrio cholerae*, and *Streptococcus pneumoniae* ParBs controlling transcription of neighboring genes, but again the repression is weak^62–64^. Here, our data suggest that KorA has a role in switching KorB from a sliding clamp, ineffective at transcription repression, to a sitting clamp which is more effective at repression. Stationary *OA*-bound KorA captures KorB sliding towards it from a distal *OB* site (Figure 7B). Since *OA* is invariably found next to core promoter elements, a stationary *OA*-bound KorA-KorB co-complex likely provides a larger and more persistent steric hindrance to RNAP than KorA or KorB alone (Figure 7B). Furthermore, our optical tweezers experiments suggest that forming a KorB-KorA complex improves retention of KorA at the *OA* site (Figure 5), providing a more stable hindrance to RNAP (Figure 7A-B). It remains unclear how KorB binding reduces the dissociation of KorA from DNA. Unfortunately, we lack a structure of full-length KorB in complex with DNA-bound KorA since we removed the 52-AA predicted intrinsically disordered linker region and the C-terminal domain of KorB to facilitate crystallization. Given that KorA-DNA docks underneath the DBD of KorB, we speculate that KorB entraps both DNA and DNA-bound KorA within its lumen (Figure 7A). The 52-AA linker that connects the DBD and CTD and constitutes KorB’s lumen is theoretically large enough to encompass a complex of KorA on DNA, potentially trapping or preventing recently dissociated KorA from diffusing too far before it re-binds *OA* DNA. The equivalent lumen in canonical ParB proteins is half the size of that in the KorB clamp (∼20 AA)^11,22,24^, and there is no identified KorA homolog in the chromosomal ParABS system. Future studies are necessary to experimentally test whether the distinct length of the KorB lumen is biologically significant and if this feature co-evolved with the ability to interact with a partner KorA.

While KorB promotes KorA retention on *OA* DNA, KorA potentially also helps KorB to entrap DNA for a longer time (Figure 7A). Our finding that KorA binding causes apo-KorB to close the clamp independently of *OB* DNA and CTP (Figure 4B) suggests that KorA might prolong the closed-clamp state of KorB-CTP, even in the case that bound KorB eventually hydrolyzes CTP. We speculate that a self-reinforcing interplay between KorA and KorB on DNA cooperatively creates a super-repressive complex. Finally, our data also suggest that this co-repression occurs via a competitive occlusion mechanism (Figure 6), with KorAB exploiting an intrinsically unstable open RNAP-promoter complex to exclude RNAP holoenzymes from the target promoters (Figure 7B).

### Regulating gene expression from a distance: DNA looping vs. DNA sliding

By providing evidence that KorB-CTP can slide, while entrapping DNA, from its initial loading site *OB* over a long distance to reach the promoter and repress transcription (Figure 7B), we are now able to reconcile seemingly incongruous observations from previous studies. For example, moving the *OB* site upstream of the promoter more than ∼1.5 kb did not reduce the efficiency of repression by KorB, and neither did moving the *OB* site downstream of the promoter^13^. However, repression and cooperativity with KorA were alleviated significantly when a DNA-bound “roadblock” (*lacO*-bound LacI) was inserted in between *OB* and the promoter^13^. Furthermore, moving the *OB* site to the opposite face of the DNA helix (by inserting/removing 5 bp i.e., half a turn of DNA helix) did not significantly affect KorB repression and cooperativity with KorA, suggesting that DNA looping might not play a major role in KorB-mediated long-rage transcriptional repression. In agreement with this, we found no evidence for KorB-mediated DNA condensation (Figure S2C), suggesting that KorB does not bridge/loop DNA.

Transcriptional regulation by a DNA-sliding clamp, as observed for KorB-CTP here, is reminiscent of the activation of virulence genes in *Shigella flexneri* by a VirB-CTP sliding clamp^65–67^. However, how a sliding VirB mechanistically counteracts the silencing activity of a nucleoid-associated protein, H-NS, to allow transcription is incompletely understood. In another case, activation of T4-phage late genes requires a gp45-gp33-gp55-RNAP complex^68,69^. T4-encoded gp45 protein is a DNA-sliding clamp with a primary function in phage DNA replication, however, gp45 also moonlights as a transcription activator^68^. The ability of KorA to trap sliding clamp KorB is functionally similar to eukaryotic CTCF and cohesin, respectively^70^. Cohesin, a DNA-entrapping protein, translocates on DNA and extrudes genomic DNA into loops. A DNA-binding protein CTCF restrains the translocation and loop extrusion by cohesin at specific DNA sites, thus switching a symmetrical DNA loop extrusion to an asymmetrical one, to generate topologically associating domains (TADs) on the chromosome^70,71^. TADs are now known to play important roles in gene regulation and recombination during development and disease^72^, and the clamp-trapping ability of CTCF is crucial to switch the function of cohesin between sister chromosome cohesion and gene expression regulation and genome folding. Our new insights into KorAB might have an impact beyond the bacterial transcription field, providing a conceptual advance in our understanding of phage, bacterial, and eukaryotic transcriptional regulation, and a new model for studying the analogous mechanisms deployed by CTCF-Cohesin and KorA-KorB.

### Final perspectives

Our work provides further insights into the mechanisms underlying gene expression regulation and stable inheritance of a broad host range multidrug resistance plasmid. Our finding that KorB requires CTP as a co-factor raises the tantalizing possibility that drugging the CTP-binding pocket will interfere with KorB function. Such a therapeutic could re-sensitize bacteria to antibiotics they may otherwise be resistant to, while also preventing the spread of multi-drug resistant plasmids. Finally, beyond the classical role of CTP in DNA segregation, the discovery of broader roles for CTP in controlling Noc membrane-binding activity and in gene expression regulation by KorB and VirB suggest that CTP-dependent molecular switches are likely to be more widespread in biology than previously appreciated^7,25,65–67,73^. We speculate that future discovery and characterization of further CTP-binding proteins will ultimately lead to a full appreciation of CTP switches in biology.

## MATERIALS AND METHODS

### Key resources table

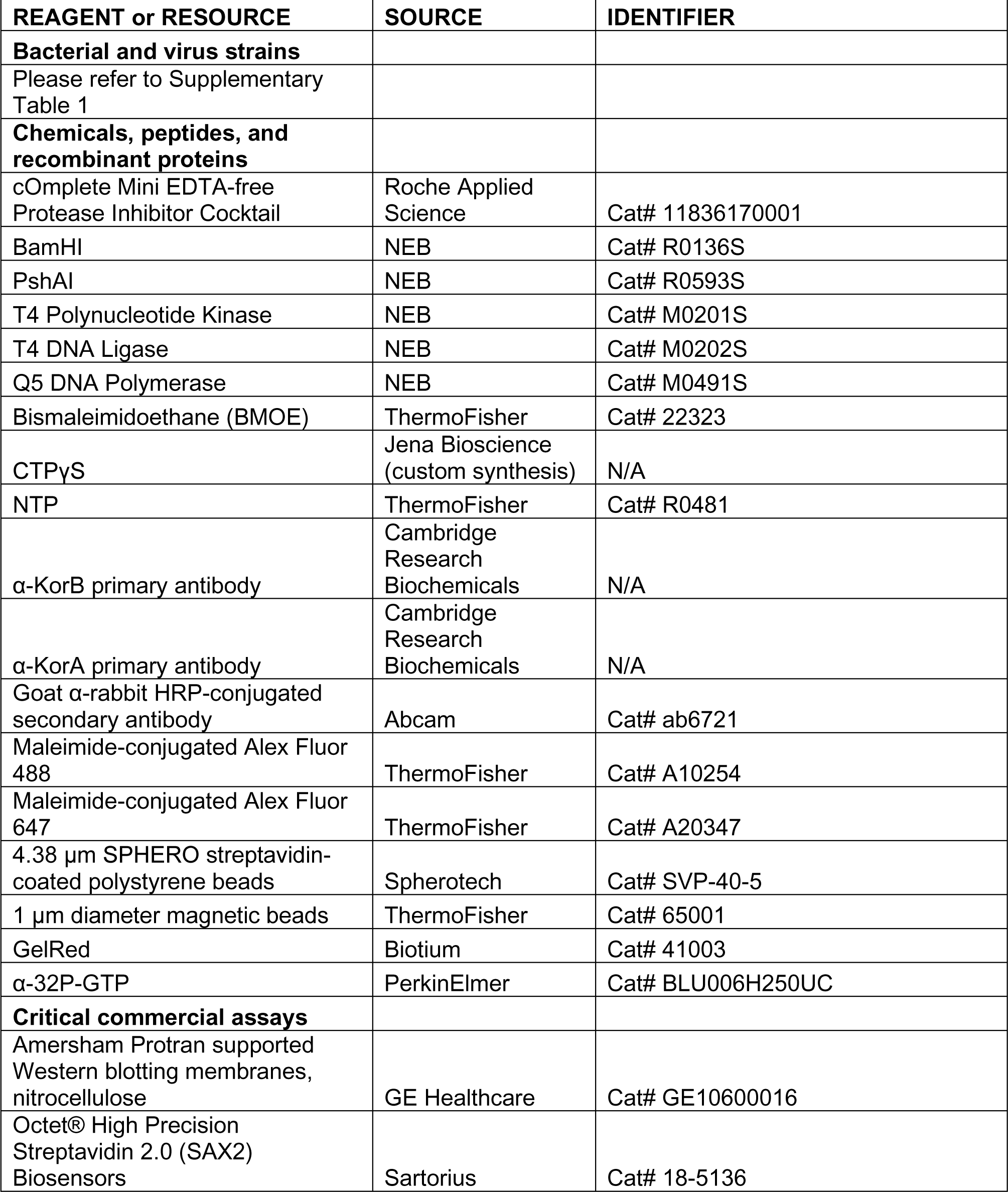
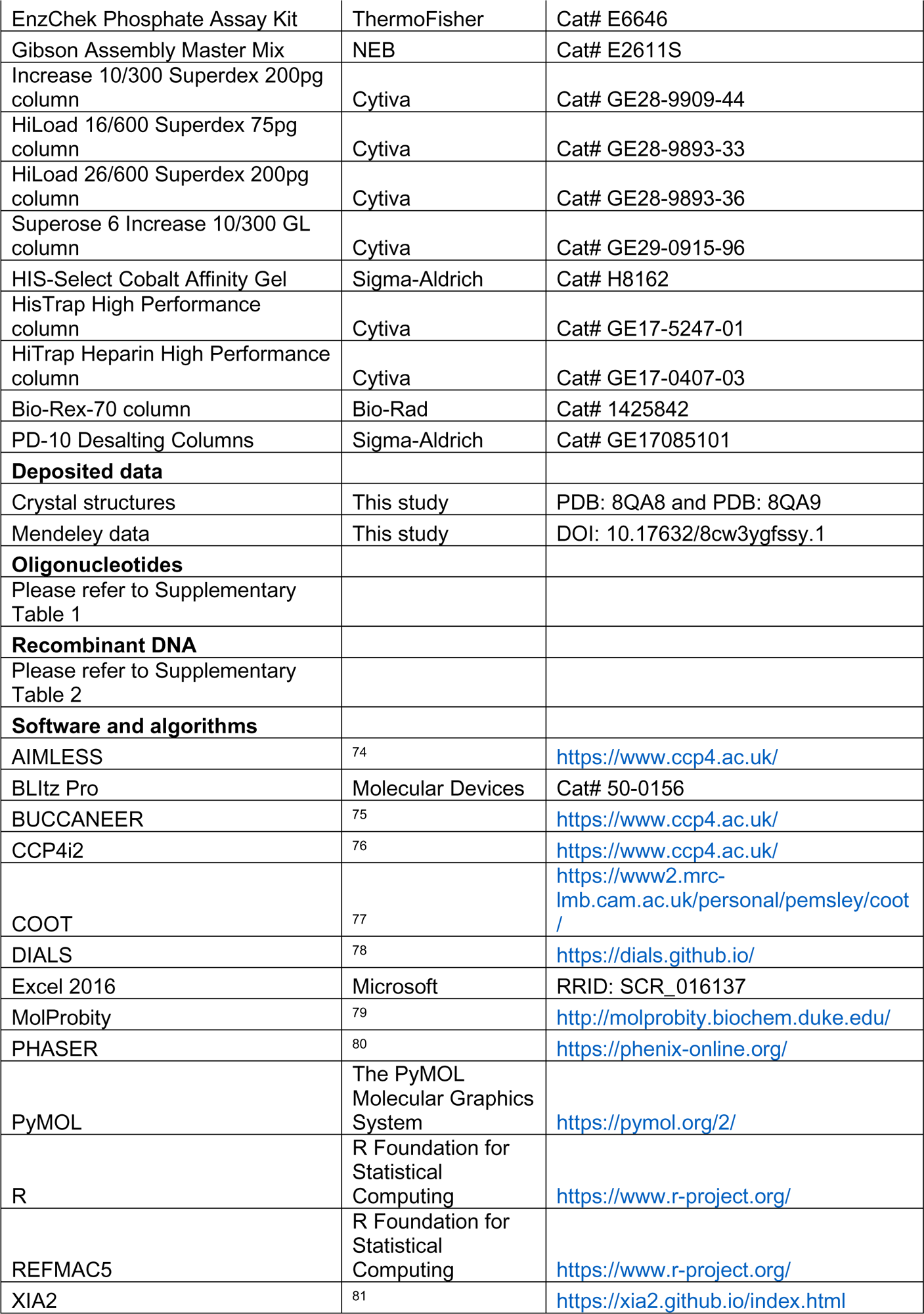
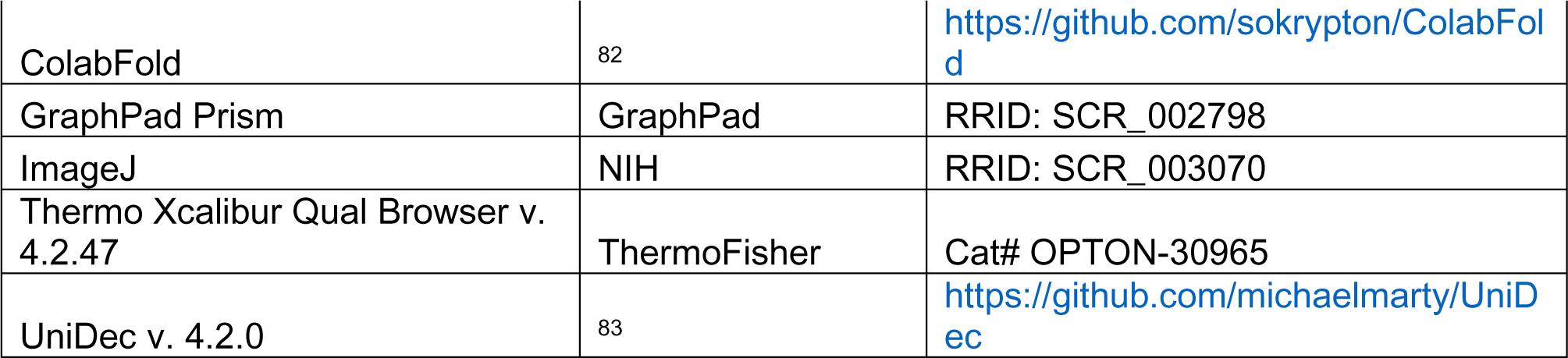

### Materials availability

Plasmids and strains (Supplementary Table 1) generated in this study are available upon request.

### Data and code availability

The crystallographic structures of KorBΔN30ΔCTD and KorBΔN30ΔCTD-KorA-*OA* have been deposited in the PDB with accession codes: 8QA8 and 8QA9, respectively. All images and source data presented in figures are available in Mendeley (https://data.mendeley.com/preview/8cw3ygfssy?a=3d52d331-8e0e-450d-9740-6ddcf18570ef). This paper does not report original code.

## Materials and Methods

### Strains, media, and growth conditions

*E. coli* strains were grown in LB medium. When appropriate the media was supplemented with antibiotics at the following concentrations (liquid/solid [μg/mL]): carbenicillin (50/100), chloramphenicol (20/30), kanamycin (30/50), streptomycin (50/50), tetracycline (12.5/12.5).

### Plasmid and strain construction

A dsDNA fragment containing a desired sequence was chemically synthesized (gBlocks, IDT). The target plasmid was double-digested with restriction enzymes and gel-purified. A 10 μL reaction mixture was created with 5 μL 2X Gibson master mix (NEB) and 5 μL of combined equimolar concentration of purified backbone and gBlock(s). This mixture was incubated at 50°C for 60 min. Gibson assembly was possible owing to shared sequence similarity between the digested backbone and the gBlock fragment(s). The resulting plasmids were verified by Sanger sequencing (Genewiz, UK).

#### Construction of pET21b::korB-(his)6 (WT and mutants)

DNA fragments containing mutated *korB* genes (*korB\**) were chemically synthesized (gBlocks, IDT). The NdeI-HindIII-cut pET21b plasmid backbone and *korB\** gBlocks fragments were assembled using a 2X Gibson master mix (NEB). Gibson assembly was possible owing to a 37-bp sequence shared between the NdeI-HindIII-cut pET21b backbone and the gBlocks fragment. The resulting plasmids were verified by Sanger sequencing (Genewiz, UK).

#### Construction of pET21b::korA-(his)6 (WT and mutants)

DNA fragments containing mutated *korA* genes (*korA\**) were chemically synthesized (gBlocks, IDT). The NdeI-HindIII-cut pET21b plasmid backbone and *korA\** gBlocks fragments were assembled using a 2X Gibson master mix (NEB). Gibson assembly was possible owing to a 37-bp sequence shared between the NdeI-HindIII-cut pET21b backbone and the gBlocks fragment. The resulting plasmids were verified by Sanger sequencing (Genewiz, UK).

#### Construction of pBAD33::korB (WT and mutants)

DNA fragments containing mutated *korB* genes (*korB\**) were chemically synthesized (gBlocks, IDT). The SacI-HindIII-cut pBAD33 plasmid backbone and *korB\** gBlocks fragments were assembled using a 2X Gibson master mix (NEB). Gibson assembly was possible owing to a 38-bp sequence shared between the SacI-HindIII-cut pBAD33 backbone and the gBlocks fragment. The resulting plasmids were verified by Sanger sequencing (Genewiz, UK).

#### Construction of pDM1.2::korA (WT and mutants)

DNA fragments containing mutated *korA* genes (*korA\**) were chemically synthesized (gBlocks, IDT). The EcoRI-SalI-cut pDM2.1 plasmid backbone and *korA\** gBlocks fragments were assembled using a 2X Gibson master mix (NEB). Gibson assembly was possible owing to a 38-bp sequence shared between the EcoRI-SalI-cut pDM2.1 backbone and the gBlocks fragment. The resulting plasmids were verified by Sanger sequencing (Genewiz, UK).

#### Construction of pSC101::PkorA (WT and mutants)

DNA fragments containing mutated *PkorA* promoters (*PkorA\**) were chemically synthesized (gBlocks, IDT). The BamHI-cut pSC101 plasmid backbone and *PkorA\** gBlocks fragments were assembled using a 2X Gibson master mix (NEB). Gibson assembly was possible owing to a 38-bp sequence shared between the BamHI-cut pSC101 backbone and the gBlocks fragment. The resulting plasmids were verified by Sanger sequencing (Genewiz, UK).

#### Construction of pUC19::146-bp-PkorA

A 146-bp DNA fragment containing *PkorA* was chemically synthesized (gBlocks, IDT) and subsequently 5’-phosphorylated using T4 PNK (NEB). Phosphorylated 146-bp *PkorA* DNA was blunt-end ligated with a dephosphorylated SmaI-cut pUC19 using T4 DNA ligase (NEB). The resulting plasmid was verified by Sanger sequencing (Genewiz, UK).

#### Construction of pUC19::146-bp-PkorA:λP_R_

To clone the 146-bp *PkorA:λP_R_* discriminator DNA into pUC19, pUC19::146bp-*PkorA* was used as a template for site-directed mutagenesis with Q5 DNA Polymerase (NEB) using the primers 5’-AACATTTCTCGCACG-3’ and 5’-TAGCTAAACTGGTTGCATGTGCTGGCG-3’ at an annealing temperature of 57°C. PCR product was introduced into *E. coli* DH5α and carbenicillin-resistant colonies were isolated. Subsequently, plasmids were isolated and verified by Sanger sequencing.

#### Construction of E. coli DH5α and BL21 pLysS strains containing the desired combinations of plasmids

Plasmids were introduced/co-introduced into *E. coli* DH5α or *E. coli* BL21 pLysS via heat shock transformation (42°C, 30 s) in the required combinations. The combinations of strains and plasmids can be found in Supplementary Table S1.

### Protein overexpression and purification

#### KorA and KorB (WT and mutants)

Protein preparation for crystallization, ITC, and biochemical experiments, excluding BMOE crosslinking, was performed as follows. C-terminally His-tagged KorA and KorB (WT and mutants) were expressed from the plasmid pET21b in *E. coli* Rosetta (BL21 DE3) pLysS competent cells (Merck, UK). 120 mL of overnight culture was used to inoculate 6 L of LB with selective antibiotics. Cultures were incubated at 37°C with shaking at 220 rpm until OD_600_ ∼0.6. Cultures were cooled for 2 hrs at 4°C before isopropyl-β-D-1-thiogalactopyranoside (IPTG) was added to a final concentration of 0.5 mM. The cultures were incubated overnight at 16°C with shaking at 220 rpm before cells were pelleted by centrifugation. Cell pellets were resuspended in buffer A (100 mM Tris-HCl, 300 mM NaCl, 10 mM imidazole, 5% (v/v) glycerol, pH 8.0) with 5 mg lysozyme (Merck, UK) and a cOmplete EDTA-free protease inhibitor cocktail tablet (Merck, UK) at room temperature for 30 min with gentle rotation. Cells were lysed on ice with 10 cycles of sonication: 15 s on / 15 s off at an amplitude of 20 microns, pelleted at 16,000 rpm for 35 mins at 4°C and the supernatant filtered through a 0.22 µm sterile filter (Sartorius, UK). Clarified lysate was loaded onto a 1 mL HisTrap HP column (Cytiva, UK) pre-equilibrated with buffer A. Protein was eluted from the column using an increasing gradient of imidazole (10-500 mM) in the same buffer. Desired protein fractions were pooled and diluted in buffer B (100 mM Tris-HCl, 20 mM NaCl, 5% v/v glycerol, pH 8.0) until the final concentration of NaCl was 60 mM. Pooled fractions were loaded onto a 1 mL Heparin HP column (Cytiva, UK) pre-equilibrated with buffer B. Protein was eluted from the column using an increasing gradient of NaCl (20-1000 mM) in the same buffer. Desired protein fractions were pooled and loaded onto a preparative-grade HiLoad 16/600 Superdex 75pg gel filtration column (GE Healthcare, UK) pre-equilibrated with elution buffer (10 mM Tris-HCl, 150 mM NaCl, pH 8.0). Desired fractions were identified and analyzed for purity via SDS-PAGE before being pooled. Aliquots were flash-frozen in liquid nitrogen and stored at -80°C. For protein samples to be used for protein-nucleotide binding ITC experiments, Mg^2+^ was introduced via an overnight dialysis step at 4°C in buffer containing 10 mM Tris-HCl, 150 mM NaCl, 5 mM MgCl_2_, pH 8.0 before concentration and quantification as described above.

Protein preparations for BMOE crosslinking were purified using a one-step Ni-affinity column with all buffers adjusted to pH 7.4 for optimal crosslinking. Purified proteins were subsequently desalted using a PD-10 column (Merck, UK) before being concentrated using an Amicon Ultra-4 10 kDa cut-off spin column (Merck, UK). Final protein samples were aliquoted and stored at -80°C in storage buffer (100 mM Tris-HCl, 300 mM NaCl, 5% v/v glycerol, 1 mM TCEP, pH 7.4).

Both biological (new sample preparations from a stock aliquot) and technical (same sample preparation) replicates were performed for assays in this study.

#### E. coli His_10_-PPX-RNAP

Plasmid pVS11 (also called pEcrpoABC(-XH)Z)^84,85^ was used to overexpress each subunit of *E. coli* RNAP (full-length α, β, ω) as well as β’-PPX-His_10_ (PPX; PreScission protease site, LEVLFQGP, GE Healthcare) were co-introduced with a pACYCDuet-1::*E.coli rpoZ* into *E. coli* BL21(DE3) to ensure saturation of all RNAPs with *E. coli* RpoZ. Cells were grown in the presence of 100 µg/mL ampicillin and 34 μg/mL chloramphenicol to an OD_600_ of ∼0.6 in a 37°C shaker. Protein expression was induced with 1 mM IPTG (final concentration) for 4 hrs at 30°C. Cells were harvested by centrifugation and resuspended in 50 mM Tris-HCl pH 8.0, 5% w/v glycerol, 10 mM DTT, 1 mM PMSF, and 1X protease inhibitor cocktail. For 200X protease inhibitor cocktail (40 mL volume), the following are dissolved into 100% ethanol (696 mg PMSF, 1.248 g benzamidine, 20 mg chymostatin, 20 mg leupeptin, 4 mg pepstatin A, and 40 mg aprotinin).

After lysis by French press (Avestin) at 4°C, the lysate was centrifuged twice for 30 mins each. Polyethyleneimine (PEI, 10% w/v pH 8.0, Acros Organics, ThermoFisher Scientific) was slowly added to the supernatant to a final concentration of ∼0.6% PEI with continuous stirring. The mixture was stirred at 4°C for an additional 25 mins, then centrifuged for 1.5 hrs at 4°C. The pellets were washed three times with 50 mM Tris-HCl pH 8.0, 500 mM NaCl, 10 mM DTT, 5% w/v glycerol, 1 mM PMSF, 1X protease inhibitor cocktail. For each wash, the pellets were homogenized and then centrifuged again. RNAP was eluted by washing the pellets three times with 50 mM Tris-HCl pH 8.0, 1 M NaCl, 10 mM DTT, 5% w/v glycerol, 1X protease inhibitor cocktail, 1 mM PMSF. The PEI elutions were combined and precipitated overnight with ammonium sulfate at a final concentration of 35% w/v. The mixture was centrifuged, and the pellets were resuspended in 20 mM Tris-HCl pH 8.0, 1 M NaCl, 5% w/v glycerol, 1 mM β-mercaptoethanol (BME). The mixture was loaded onto two 5 mL HiTrap IMAC HP columns (Cytiva) for a total column volume of 10 mL. RNAP(β’-PPX-His_10_) was eluted at 250 mM imidazole in column buffer (20 mM Tris-HCl pH 8.0, 1 M NaCl, 5% w/v glycerol, 1 mM BME). The eluted RNAP fractions were combined and dialyzed into 10 mM Tris-HCl pH 8.0, 0.1 mM EDTA pH 8.0, 100 mM NaCl, 5% w/v glycerol, 5 mM DTT. The sample was then loaded onto a 40 mL Bio-Rex-70 column (Bio-Rad), washed with 10 mM Tris-HCl pH 8.0, 0.1 mM EDTA, 5% w/v glycerol, 5 mM DTT in isocratic steps of increasing concentration of NaCl (eluted at 0.5 M NaCl). The eluted fractions were combined, concentrated by centrifugal filtration, then loaded onto a 320 mL HiLoad 26/600 Superdex 200 column (Cytiva) pre-equilibrated in gel filtration buffer (10 mM Tris-HCl pH 8.0, 0.1 mM EDTA pH 8.0, 0.5 M NaCl, 5% w/v glycerol, 5 mM DTT). The eluted RNAP was concentrated to ∼8-10 mg/mL by centrifugal concentration and supplemented with glycerol to 20% w/v, flash frozen in liquid nitrogen, and stored at −80°C.

#### E. coli His_10_-SUMO-σ^70^

Plasmid pSAD1403^86^ was introduced into *E. coli* BL21(DE3). The cells were grown in the presence of 50 μg/mL kanamycin to an OD_600_ of ∼0.6 at 37°C. Protein expression was induced with 1 mM IPTG for 1-1.5 hrs at 30°C. Cells were harvested by centrifugation and resuspended in 20 mM Tris-HCl pH 8.0, 500 mM NaCl, 0.1 mM EDTA pH 8.0, 5 mM imidazole, 5% w/v glycerol, 0.5 mM BME, 1 mM PMSF, 1X protease inhibitor cocktail. After lysis by French press (Avestin) at 4°C, cell debris was removed by centrifugation twice. The lysate was loaded onto two 5 mL HiTrap IMAC HP columns (Cytiva) for a total column volume of 10 mL. His_10_-SUMO-σ^70^ was eluted at 250 mM imidazole in 20 mM Tris-HCl pH 8.0, 500 mM NaCl, 0.1 mM EDTA pH 8.0, 5% w/v glycerol, 0.5 mM BME. Peak fractions were combined, cleaved with Ulp1 protease, and dialyzed against 20 mM Tris-HCl pH 8.0, 500 mM NaCl, 0.1 mM EDTA pH 8.0, 5% w/v glycerol, 0.5 mM BME, resulting in a final concentration of 25 mM imidazole. The cleaved sample was loaded onto one 5 mL HiTrap IMAC HP to remove His_10_-SUMO-tag along with any remaining uncleaved σ^70^. Untagged σ^70^ fractions are pooled and diluted with 10 mM Tris-HCl pH 8.0, 0.1 mM EDTA pH 8.0, 5% w/v glycerol, 1 mM DTT until the conductivity corresponds to NaCl concentration slightly below 200 mM. The diluted sample was injected onto three 5 mL HiTrap Heparin HP columns (total column volume of 15 mL; Cytiva) which were pre-equilibrated at the same diluent buffer but with 200 mM NaCl, with a gradient to 1 M NaCl, with the first major peak as the target peak. The target peak sample was pooled and concentrated by centrifugal filtration before being loaded onto a HiLoad 16/60 Superdex 200 (Cytiva) which was pre-equilibrated in 20 mM Tris-HCl pH 8.0, 500 mM NaCl, 5% w/v glycerol, 1 mM DTT. Peak fractions of σ^70^ were pooled, supplemented with glycerol to a final concentration of 20% w/v, flash-frozen in liquid nitrogen, and stored at -80°C.

### Protein crystallization

Crystallization screens for both KorBΔN30ΔCTD-CTPγS and KorBΔN30ΔCTD-KorA-*OA* complexes were performed in sitting-drop vapor diffusion format in MRC2 96-well crystallization plates. Drops consisted of 0.3 μL precipitant solution and 0.3 μL protein complex with incubation at 293 K.

#### KorBΔN30ΔCTD-CTPγS

Purified His-tagged KorBΔN30ΔCTD was premixed at 20 mg/mL with 1 mM MgCl_2_ and 1 mM CTPγS in buffer containing 10 mM Tris-HCl, 150 mM NaCl, pH 8.0. The KorBΔN30ΔCTD-CTPγS crystals grew in a solution containing 160 mM LiOAc and 2.0 M ammonium sulfate. Suitable crystals were cryoprotected with 20% (v/v) ethylene glycol and mounted in Litholoops (Molecular Dimensions, UK). Crystals were flash cooled by plunging into liquid nitrogen.

#### KorBΔN30ΔCTD-KorA-OA

Purified His-tagged KorBΔN30ΔCTD was combined with purified His-tagged KorA in equimolar concentrations before being purified by gel filtration as described above. The protein complex was premixed at 20 mg/mL with a 14-bp dsDNA (*OA*: TGTTTAGCTAAACA) at a molar ratio 1:1.2 (protein complex:DNA) in buffer containing 10 mM Tris-HCl, 150 mM NaCl, pH 8.0. Crystals grew in a solution containing 1.95 M ammonium sulfate and 0.1 M NaOAc, pH 4.6. Suitable crystals were cryoprotected with 25% (v/v) glycerol and mounted in Litholoops (Molecular Dimensions, UK). Crystals were flash cooled by plunging into liquid nitrogen.

### Structure determination and refinement

X-ray data were recorded either on beamline I04-1 or I03 at the Diamond Light Source (Oxfordshire, UK) using either an Eiger2 XE 9M or an Eiger2 XE 16M hybrid photon counting detector (Dectris), respectively, with crystals maintained at 100 K by a Cryojet cryocooler (Oxford Instruments). Diffraction data were integrated and scaled using DIALS^81^ via the XIA2 expert system^81^ then merged using AIMLESS^74^. The majority of the downstream analysis was performed through the CCP4i2 graphical user interface^76^. Data collection statistics are summarized in Table 1.

**Table 1.**
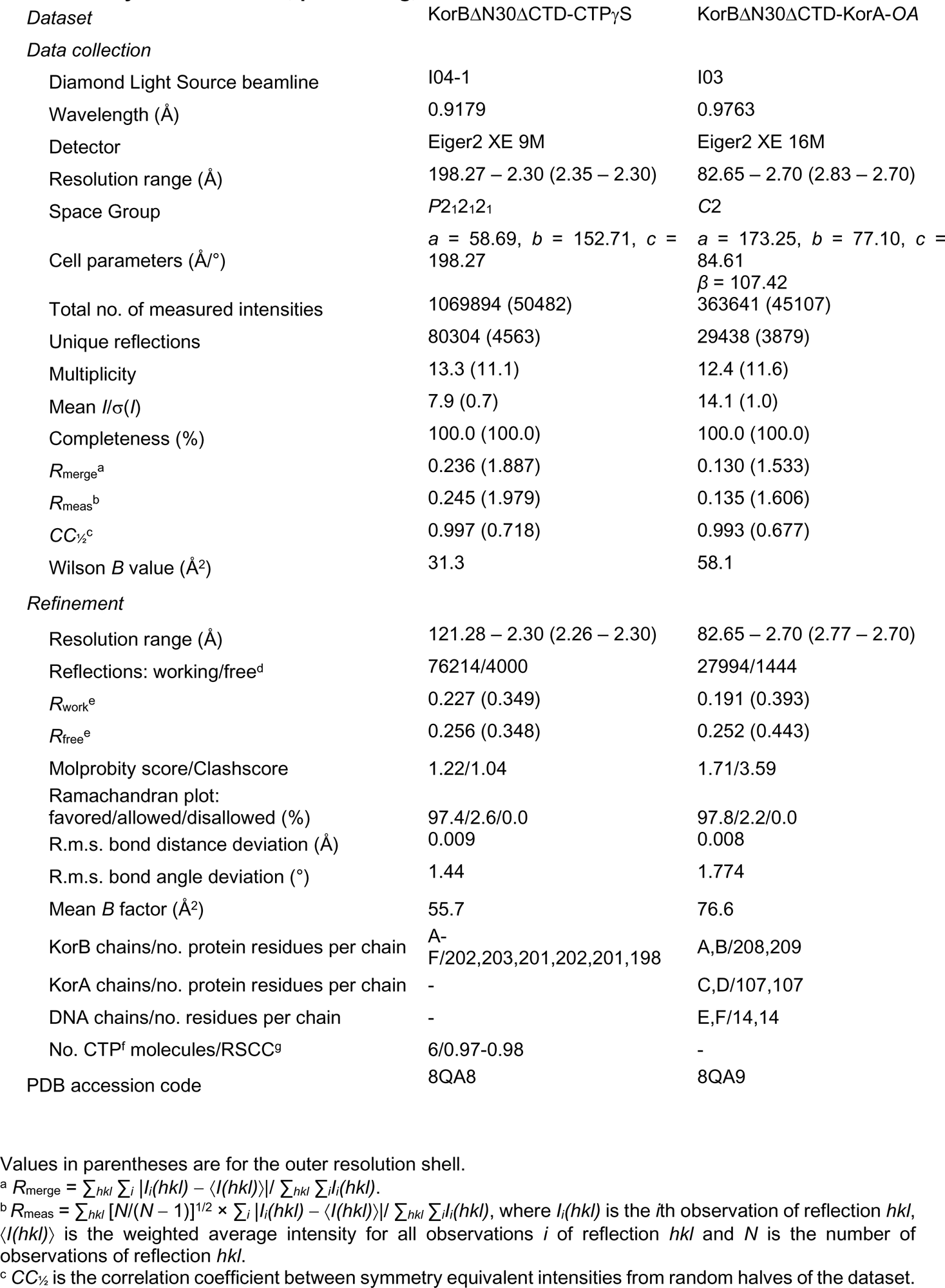
X-ray data collection, processing, and refinement statistics.

| Dataset | KorBΔN30ΔCTD-CTPγS | KorBΔN30ΔCTD-KorA-OA |
| --- | --- | --- |
| *Data collection* |  |  |
| Diamond Light Source beamline | I04-1 | I03 |
| Wavelength (Å) | 0.9179 | 0.9763 |
| Detector | Eiger2 XE 9M | Eiger2 XE 16M |
| Resolution range (Å) | 198.27 – 2.30 (2.35 – 2.30) | 82.65 – 2.70 (2.83 – 2.70) |
| Space Group | *P*2<sub>1</sub>2<sub>1</sub>2<sub>1</sub> | *C*2 |
| Cell parameters (Å/°) | *a* = 58.69, *b* = 152.71, *c* = 198.27 | *a* = 173.25, *b* = 77.10, *c* = 84.61<br>*β* = 107.42 |
| Total no. of measured intensities | 1069894 (50482) | 363641 (45107) |
| Unique reflections | 80304 (4563) | 29438 (3879) |
| Multiplicity | 13.3 (11.1) | 12.4 (11.6) |
| Mean *I*/σ(*I*) | 7.9 (0.7) | 14.1 (1.0) |
| Completeness (%) | 100.0 (100.0) | 100.0 (100.0) |
| *R*<sub>merge</sub><sup>a</sup> | 0.236 (1.887) | 0.130 (1.533) |
| *R*<sub>meas</sub><sup>b</sup> | 0.245 (1.979) | 0.135 (1.606) |
| *CC*<sub>1/2</sub><sup>c</sup> | 0.997 (0.718) | 0.993 (0.677) |
| Wilson *B* value (Å<sup>2</sup>) | 31.3 | 58.1 |
| *Refinement* |  |  |
| Resolution range (Å) | 121.28 – 2.30 (2.26 – 2.30) | 82.65 – 2.70 (2.77 – 2.70) |
| Reflections: working/free<sup>d</sup> | 76214/4000 | 27994/1444 |
| *R*<sub>work</sub><sup>e</sup> | 0.227 (0.349) | 0.191 (0.393) |
| *R*<sub>free</sub><sup>e</sup> | 0.256 (0.348) | 0.252 (0.443) |
| Molprobity score/Clashscore | 1.22/1.04 | 1.71/3.59 |
| Ramachandran plot:<br>favored/allowed/disallowed (%) | 97.4/2.6/0.0 | 97.8/2.2/0.0 |
| R.m.s. bond distance deviation (Å) | 0.009 | 0.008 |
| R.m.s. bond angle deviation (°) | 1.44 | 1.774 |
| Mean *B* factor (Å<sup>2</sup>) | 55.7 | 76.6 |
| KorB chains/no. protein residues per chain | A-F/202,203,201,202,201,198 | A,B/208,209 |
| KorA chains/no. protein residues per chain | - | C,D/107,107 |
| DNA chains/no. residues per chain | - | E,F/14,14 |
| No. CTP<sup>f</sup> molecules/RSCC<sup>g</sup> | 6/0.97-0.98 | - |
| PDB accession code | 8QA8 | 8QA9 |

Values in parentheses are for the outer resolution shell.
<sup>a</sup> $R_{\text{merge}} = \sum_{hkl} \sum_i |I_i(hkl) - \langle I(hkl) \rangle| / \sum_{hkl} \sum_i I_i(hkl)$ .
<sup>b</sup> $R_{\text{meas}} = \sum_{hkl} [N/(N-1)]^{1/2} \times \sum_i |I_i(hkl) - \langle I(hkl) \rangle| / \sum_{hkl} \sum_i I_i(hkl)$ , where $I_i(hkl)$ is the *i*th observation of reflection *hkl*, $\langle I(hkl) \rangle$ is the weighted average intensity for all observations *i* of reflection *hkl* and *N* is the number of observations of reflection *hkl*.
<sup>c</sup> *CC*<sub>1/2</sub> is the correlation coefficient between symmetry equivalent intensities from random halves of the dataset.

X-ray data for KorBΔN30ΔCTD-CTPγS were collected from a single crystal at a wavelength of 0.9179 Å and processed to 2.3 Å-resolution in space group *P*2_1_2_1_2_1_, with approximate cell parameters of *a* = 58.7, *b* = 152.8, *c* = 198.3 Å. Analysis of the likely composition of the asymmetric unit suggested that it could contain between four and eight copies of the KorB subunit with an estimated solvent content in the range 43-72%.

Structural predictions for KorB were generated using AlphaFold2 (AF2)^87^, as implemented through ColabFold^82^. There was good sequence coverage, and the predicted local distance difference test (pLDDT) scores were generally good (e.g. average of 85 from the rank 1 prediction). For a single subunit simulation, the predicted aligned error (PAE) scores indicated a two-domain structure with very low confidence in the relative placement of the two domains, whilst for a dimer simulation, the PAE scores suggested high confidence in the relative placement of all four domains. Consistent with this, all five independently generated models were closely superposable.

The KorBΔN30ΔCTD-CTPγS complex structure was solved via molecular replacement using PHASER^80^. A dimer template was prepared from the rank 1 AF2 model using the “Process Predicted Models” CCP4i2 task, which removed low-confidence regions (based on pLDDT) and converted the pLDDT scores in the *B*-factor field of the PDB coordinate files to pseudo *B* factors. Initial attempts used an isomorphous dataset at 3.25 Å-resolution. After much trial and error, searching with separate templates corresponding to the two KorB domains showed the most promise. PHASER produced a partial solution where three pairs of domains were juxtaposed such that they could be connected into single subunits using COOT^77^. One of the latter was then used as the template for a subsequent run, where PHASER placed five of these in the ASU, which were arranged as two dimers and a single subunit. Inspection of this solution in COOT revealed residual electron density adjacent to the latter which could be filled by a sixth subunit by extrapolation from one of the dimers. This final composition of six protomers per ASU gave an estimated solvent content of 57%. When the 2.3 Å-resolution dataset became available, the preliminary model was refined directly against this in REFMAC5^88^. At this stage, residual electron density at the interface between the N-terminal domains of each dimer indicated the presence of two symmetry-related ligands. These were built as CTP molecules, as it was not possible to define the locations of the sulfur atoms of CTPγS. The model was completed through several iterations of model building in COOT and restrained refinement in REFMAC5. Pairwise superpositions of the three dimers gave overall RMS deviations of 0.63 – 1.18 Å, indicating that they were closely similar. Comparison against the AF2 dimer model showed that this experimental structure had a domain-swapped arrangement, whilst the predicted structure did not. Refinement and validation statistics are summarized in Table 1.

X-ray data for KorBΔN30ΔCTD-KorA-*OA* were collected from a single crystal at a wavelength of 0.9763 Å (2 x 360° passes) and processed to 2.7 Å-resolution in space group *C*2, with approximate cell parameters of *a* = 173.2, *b* = 77.1, *c* = 84.6 Å, *β* = 107.4°. Analysis of the likely composition of the asymmetric unit suggested that it contained a single complex comprised of two copies each of the KorA and KorB subunits and a single DNA duplex, giving an estimated solvent content of 61%.

The structure was solved via molecular replacement using PHASER. Separate templates were prepared for the two KorB domains from the A chain of the above KorBΔN30ΔCTD-CTPγS complex; for the KorA subunit by taking a single chain from the previously solved KorA-DNA complex (PDB code 2W7N)^89^; and for the DNA by generating an ideal B-form DNA duplex in COOT from the palindromic sequence TGTTTAGCTAAACA. PHASER was able to locate the four domains expected for a KorB dimer and the DNA duplex, but not the two KorA subunits. However, after refinement in REFMAC5, a clear difference density was visible for the missing KorA subunits. These could be manually placed from a superposition of the KorA-DNA complex. Several sulfates were built into the density, presumably derived from the precipitant. Two of these occupied positions equivalent to the β-phosphates of the CTP ligands in the previous structure. The model was completed through several iterations of model building in COOT and restrained refinement in REFMAC5. In contrast to the KorB dimer from the CTP complex, this dimer does not have a domain-swapped architecture. Moreover, a superposition of this KorB dimer onto the AF2 dimer model revealed that they were almost identical at the protein backbone level, giving an overall RMS deviation of only 1.01 Å. Refinement and validation statistics are summarized in Table 1.

### Measurement of protein-DNA interaction by BLI assay

BLI experiments were conducted using a BLItz system equipped with High Precision Streptavidin 2.0 (SAX2) Biosensors (ForteBio, UK) at 22°C. The SAX2 biosensor was first hydrated in a low-salt buffer (100 mM Tris-HCl, 150 mM NaCl, 1 mM MgCl_2_, and 0.005% v/v Tween 20, pH 8.0) for at least 10 min before each experiment. Biotinylated 175-bp dsDNA was immobilized onto the surface of the SAX2 biosensor through one complete BLI cycle. Briefly, the tip of the biosensor was dipped into a binding buffer for 30 s to establish the baseline, then into 20 nM biotinylated dsDNA for 120 s, and finally into low-salt buffer for 120 s to allow for dissociation. After each cycle, the SAX2 biosensor was recycled using 120 s of washing in high-salt buffer (100 mM Tris-HCl, 2 M NaCl, 1 mM EDTA, 0.005% v/v Tween 20, pH 8.0) before being returned into low-salt buffer.

Where it was required to cut the closed-loop constructs the DNA-coated tips were dipped into 300 µL of restriction solution (267 µL of water, 30 µL of 10X CutSmart buffer (NEB), 3 µL of BamHI-HF restriction enzyme [20,000 units/mL]) for 4 hrs at 37°C. As a result, a free end was created on the immobilized DNA^90^.

All experiments were performed in triplicate. The results were analyzed in Excel and plotted using GraphPad Prism 9. Where required representative sensorgrams are presented.

### DNA preparation for in vitro NTPase, crosslinking, and ITC experiments

Palindromic single-stranded DNA oligonucleotides (*OB*: GGGAT<u>ATTTTAGCGGCTAAAAG</u>GA, *OA*: <u>TGTTTAGCTAAACA</u>) (100 µM in 1 mM Tris-HCl pH 8.0, 5 mM NaCl buffer) were heated at 98°C for 5 min before being left to cool down to room temperature overnight to form 50 µM double-stranded DNA. The core sequence of *OB* or *OA* is underlined.

### Measurement of CTPase activity by EnzChek phosphate release assay

CTP hydrolysis was monitored using an EnzCheck Phosphate Assay Kit (Thermo Fisher Scientific, UK). Samples (100 µL) containing a reaction buffer supplemented with an increasing concentration of CTP (1, 5, 10, 50, 100, 500, and 1000 µM), 0.5 µM of 24-bp *OB* DNA, and 1 µM KorB (WT or mutants) were assayed in a CLARIOstar Plus plate reader (BMG Labtech, UK) at 25°C for 5 hrs with readings every 2 min with continuous orbital shaking at 300 rpm between reads. The reaction buffer (1 mL) typically contained 740 μL Ultrapure water, 50 μL 20X reaction buffer (100 mM Tris-HCl, 2 M NaCl, and 20 mM MgCl_2_, pH 8.0), 200 μL MESG substrate solution, and 10 μL purine nucleoside phosphorylase enzyme (one unit). Reactions with buffer only or buffer + CTP + 24-bp *OB* DNA only were also included as controls. The inorganic phosphate standard curve was also constructed according to the instruction guidelines. The results were analyzed using Excel and plotted in GraphPad Prism 9. The CTPase rates were calculated using a linear regression fitting in GraphPad Prism 9. Error bars represent standard deviations from triplicate experiments.

### *In vitro* crosslinking assay using a sulfhydryl-to-sulfhydryl crosslinker BMOE

A 50 µL mixture of 8 µM KorB WT or mutants ± 1 mM NTP ± 0.5 µM 24-bp dsDNA containing *OB* or scrambled *OB* was assembled in a reaction buffer (10 mM Tris-HCl pH 7.4, 100 mM NaCl, and 1 mM MgCl_2_) and incubated for 5 min at room temperature. BMOE was added to a final concentration of 1 mM, and the reaction was quickly mixed by three pulses of vortexing. The reaction was then immediately quenched through the addition of SDS-PAGE sample buffer containing 23 mM β-mercaptoethanol. Samples were heated to 50°C for 5 min before being loaded on 12% Novex WedgeWell Tris-Glycine gels (Thermo Fisher Scientific, UK). Protein bands were stained with an InstantBlue Coomassie protein stain (Abcam, UK) and band intensity was quantified using ImageJ. The results were analyzed in Excel and plotted using GraphPad Prism 9.

For the experiments containing KorA in addition to KorB, an equimolar amount was used (8 µM of WT or mutants). The reaction was otherwise assembled identically and loaded on 4-20% Novex WedgeWell Tris-Glycine gels (Thermo Fisher Scientific, UK) for sufficient separation of KorA in the samples. Band intensity was quantified using ImageJ and the results were analyzed in Excel and plotted using GraphPad Prism 9.

### Isothermal titration calorimetry (ITC)

All ITC experiments were recorded using a MicroCal PEAQ ITC instrument (Malvern Panalytical, UK). Experiments were performed at 25°C. For protein-nucleotide binding experiments, all components were in 10 mM Tris-HCl, 150 mM NaCl, 5 mM MgCl_2_, pH 8.0 buffer. For protein-protein binding experiments, all components were in 10 mM Tris-HCl, 150 mM NaCl, pH 8.0 buffer. For each ITC run the calorimetric cell was filled with 20 μM dimer concentration of KorB (WT or mutant) and a single injection of 0.4 μL of 500 μM small molecule nucleotides or 200 μM protein partner was performed first, followed by 19 injections of 2 μL each. Injections were carried out at 150 s intervals with a stirring speed of 750 rpm. The raw titration data were integrated and fitted to a one-site binding model using the built-in software of the MicroCal PEAQ ITC instrument. Each experiment was run in duplicate. Controls of ligand into buffer and buffer into protein were performed with no signal observed. Where required representative data are presented.

### *xylE* reporter gene assays

Reporter gene assays were carried out using a modified version of Zukowski *et al* (1983)^91^. In short, *E. coli* DH5α cells containing relevant expression plasmids were grown to a logarithmic phase from a 1:100 dilution of overnight culture. Induction of KorB WT/mutant expression was achieved with 0.2% (*PkorA,* Figure 5E) and 0.02% (*PtrbB*, Figure 3) arabinose. In the three plasmid experiments, induction of KorA WT/Y84A required no additional IPTG. 10 or 50 mL of culture was pelleted and resuspended in 500 μL resuspension buffer (0.1 M sodium phosphate buffer pH 7.4, 10% v/v acetone). From this point onwards samples were kept on ice. Cells were disrupted using sonication at 10 microns for 10 sec and subsequently pelleted. The supernatant was transferred to a fresh microcentrifuge tube and assayed for catechol 2,3-oxygenase activity. Samples were diluted 1:10 in reaction buffer (0.1 M sodium phosphate buffer pH 7.4, 200 μM catechol), and incubated at room temperature for one minute before the absorbance at 374 nm was determined using a BioMate 3 spectrophotometer (Thermo Fisher Scientific, UK). Protein concentration, determined using Bradford assay, was used to normalize the samples. The results were analyzed in Excel and plotted using GraphPad Prism 9.

### Immunoblot analysis

For western blot analysis samples 200 ng total protein lysate was resuspended in 1X SDS-PAGE sample buffer and heated to 95°C for 10 min before loading. Denatured samples were run on 12% Novex WedgeWell gels (Thermo Fisher Scientific, UK) at 150 V for 55 mins. Resolved proteins were transferred to PVDF membranes using the Trans-Blot Turbo Transfer System (BioRad, UK) and incubated with a 1:5000 dilution of α-KorB primary antibody (Cambridge Research Biochemicals, UK) or with 1:300 dilution of α-KorA. Membranes were washed and subsequently probed with a 1:10000 dilution of mouse α-rabbit HRP-conjugated secondary antibody (Abcam, UK). Blots were imaged after incubation with SuperSignal West PICO PLUS Chemiluminescent Substrate (Thermo Fisher Scientific, UK) using an Amersham Imager 600 (GE HealthCare, UK). Loading controls of denatured 200 ng total protein lysate were run on 12% Novex WedgeWell gels (Thermo Fisher Scientific, UK) at 150 V for 55 mins and stained with InstantBlue Coomassie protein stain (Abcam, UK).

### Protein labeling with Alexa Fluor for Confocal-Optical Tweezers (C-trap) experiments

C-terminally His-tagged versions of KorB (A6C) and KorA (WT/Y84A variant) with an extra cysteine residue at the C-terminus were coupled to maleimide-conjugated Alexa Fluor (AF) 488 and 647, respectively. A6C was selected because it resides in a surface-exposed intrinsically disordered region at the N-terminal region of KorB. His-tagged KorB (A6C) and His-tagged KorA-extra C were purified as described for the wild-type proteins. 250 µL of 50 µM KorB (A6C) or KorA-extra C were incubated with 0.3 mM TCEP for 30 min at room temperature in a buffer containing 10 mM Tris-HCl, 300 mM NaCl, pH 7.4. Subsequently, 6 µL of 30 mM AF488 or AF647 (dissolved in DMSO) was added, and the reaction was incubated with rotation at 4°C overnight. The conjugate solution was then loaded onto a Superdex increase 200pg 10/300 gel filtration column (pre-equilibrated with 10 mM Tris-HCl, 300 mM NaCl, pH 8.0) to separate labeled KorB/A from unincorporated fluorophore. AF-labeled KorB/A was pooled and concentrated before storage as described for wild-type KorB and KorA.

### Design and construction of a DNA plasmid with 16x *OB* sites and 1x *OA* site for magnetic tweezers (MT) experiments

A DNA plasmid containing 16x *OB* sites (ATTTTAGCGGCTAAAAG) and 1x *OA* site (AATGTTTAGCTAAACCTT) was produced by modification of a pUC19 plasmid containing a single copy of each site separated by 1016 bp (pUC19_v1), following several cloning steps and methods described elsewhere^27,92^.

First, the original pUC19 plasmid (4886 bp) with one of each site was enlarged by introducing a piece of DNA obtained from a lab plasmid. This resulted in a larger plasmid (7699 bp) which, after digestion with appropriate restriction enzymes, produces the central part of an MT DNA construct with centered *OB* and *OA* sites.

To increase the number of *OB* sites two long oligonucleotides (Supplementary Table 1) containing 2x *OB* sites separated by 40 bp with a PshAI restriction site in the middle of this region were annealed by heating at 95°C for 5 min and cooling down to 20°C at a rate of -1 °C min^-1^ in hybridization buffer (10 mM Tris-HCl pH 8.0, 1 mM EDTA, 200 mM NaCl, and 5 mM MgCl_2_) followed by a phosphorylation step of the 5′-terminal ends by the T4 PNK (NEB). This dsDNA duplex was ligated into the previous plasmid of 7699 bp that already contained 1x *OA* and 1x *OB* sites digested with PshAI restriction enzyme (NEB) and dephosphorylated. These oligonucleotides were designed to lose the original PshAI site at both ends after ligation, so that once ligated into a cloning plasmid they could not be cleaved again by PshAI. The single *bona fide* PshAI site located in the middle of the duplex allows for repetition of the ligation process to be repeated as many times as desired in the cloning plasmid to add new pairs of *OB* sites. Plasmids containing 1x *OA* site and up to 8x *OB* have been obtained following this procedure. N.B. in one of the rounds of cloning and by chance, half of a previous duplex was lost during the ligation process, therefore the final plasmid contains 8x*OB* instead of 9x*OB* as expected.

A plasmid with 1x *O_A_* site and 16x *OB* was produced by PCR amplifying an 8x *OB* cassette with Phusion High-Fidelity DNA Polymerase (Thermo Scientific) (see Supplementary Table S1 for primer sequences). The PCR fragment was then digested with SpeI and XhoI (both from NEB) and ligated into the plasmid already containing 1x *OA* site and 8x *OB* copies previously digested with XbaI (NEB) and XhoI and dephosphorylated. This resulted in a plasmid with 1x *OA* site and 16x *OB* sites (8705 bp). Plasmids were introduced into *E. coli* DH5α competent cells and potentially positive colonies were then selected by colony PCR. Plasmids were purified from the cultures using a QIAprep Spin Miniprep Kit (QIAGEN), analyzed by restriction enzyme digestion, and finally verified by Sanger sequencing. This plasmid was used to produce a magnetic tweezers dsDNA construct (see below).

### Construction of a large plasmid with 8x *OB* sites and 1x *OA* site for confocal optical tweezers (C-trap) experiments

The large DNA plasmid containing 8x *OB* sites and 1x *OA* site was produced as follows. First, a large DNA plasmid with 1x *OA* site was fabricated by ligating a DNA fragment containing a single copy of the *OA* site into a previously prepared large plasmid in our laboratory that did not contain either of these sites. The fragment containing the *OA* site was obtained by PCR amplification using Phusion High-Fidelity DNA Polymerase with the pUC19 plasmid containing a single copy of the *OB* and *OA* sites as a template (see Supplementary Table S1 for primer sequences). The PCR fragment was digested with KpnI (NEB) and ligated into the previously large plasmid prepared in our laboratory that did not contain either of these sites, digested with KpnI and dephosphorylated. This resulted in a large plasmid with a single *OA* site. A DNA fragment with 8x *OB* sites was then inserted over this new large plasmid. The 8x *OB* fragment was obtained by PCR amplification (see Supplementary Table S1 for primer sequences) using Phusion High-Fidelity DNA Polymerase with the MT plasmid that contained 1x *OA* and 8x *OB* sites as a template. The PCR fragment was digested with SpeI and XbaI, and ligated into the large plasmid, which already contained 1x *OA* site, digested with XbaI and dephosphorylated, resulting in a large plasmid with 1x *OA* site and 8x *OB* sites (22394 bp). The plasmids were cloned and analyzed as described for MT plasmids. This plasmid was used to prepare a C-trap dsDNA construct (see below).

### Magnetic tweezers dsDNA construct with 16x *OB* sites and 1x *OA*

The dsDNA construct for magnetic tweezers experiments consisted of a central dsDNA fragment of 8693 bp containing 16x *OB* and 1x *OA* sites, obtained by digestion with NotI and ApaI (NEB) of the final MT plasmid described above, flanked by two highly labeled DNA fragments, one with digoxigenins and the other with biotins, of 997 bp and 140 bp, respectively, used as immobilization handles. The biotinylated handle was shorter to minimize the attachment of two beads per DNA tether. Handles for MT constructs were prepared by PCR (see Supplementary Table S1 for primers) with 200 µM final concentration of each dNTP (dGTP, dCTP, dATP), 140 µM dTTP and 66 µM Bio-16-dUTP or Dig-11 dUTP (Roche) using plasmid pSP73-JY0^93^ as template, followed by digestion with the restriction enzyme ApaI or PspOMI (NEB), respectively. Labeled handles were ligated to the central part overnight using T4 DNA Ligase (NEB). The sample was then ready for use in MT experiments without further purification. The DNAs were never exposed to intercalating dyes or UV radiation during their production and were stored at 4°C. The sequence of the central part of the MT construct is included in Supplementary Table S2.

### C-trap dsDNA construct with 8x *OB* sites and 1x *OA*

The C-trap dsDNA construct consisted of a large central fragment of 22394 bp containing 8x *OB* and 1x *OA* sites, was produced by digestion of the large C-trap plasmid (see above) with NotI. Without further purification, the fragment was ligated to highly biotinylated handles of ∼1 kb ending in PspOMI. Handles for C-trap constructs were prepared by PCR (see Supplementary Table S1 for primers) as described for biotin-labeled MT handles. These handles were highly biotinylated to facilitate the capture of DNA molecules in the C-trap experiments. As both sides of the DNA fragment end in NotI, it is possible to generate tandem (double-length) tethers flanked by the labeled handles. The sample was ready for use in —trap experiments without further purification. The DNAs were not exposed to intercalating dyes or UV radiation during their production and were stored at 4°C. The sequence of the central part of the C-trap construct is included in Supplementary Table S2.

### Confocal optical tweezers experiments

Confocal-optical tweezers experiments were carried out using a dual optical tweezers setup combined with confocal microscopy and microfluidics (C-Trap; Lumicks)^94,95^. A computer-controlled stage allowed rapid displacement of the optical traps within a five-channel fluid cell, allowing the transfer of the tethered DNA between different channels separated by laminar flow. Channel 1 contained 4.38 µm streptavidin-coated polystyrene beads (Spherotech). Channel 2 contained the DNA substrate labelled with multiple biotins at both ends. Both DNA and beads were diluted in 20 mM HEPES pH 7.8, 100 mM KCl and 5 mM MgCl_2_. A single DNA tether was assembled by first capturing two beads in channel 1, one in each optical trap, and fishing for a DNA molecule in channel 2. The tether was then transferred to channel 3 filled with reaction buffer (10 mM Tris pH 8, 100 mM NaCl and 5 mM MgCl_2_, 1 mM DTT) to verify the length of the correct length of the DNA by force-extension curves. The DNA was then incubated for one minute in channel 4 filled with KorB and/or KorA proteins in reaction buffer and supplemented with 2 mM CTP as indicated. To reduce the fluorescence background in single KorB diffusion measurements, imaging was occasionally performed in channel 3 after protein incubation in channel 4.

The system is equipped with three laser lines for confocal microscopy (488, 532 and 635 nm). In this study the 488 nm laser was used to excite AF488-KorB and the 635 nm laser to excite AF647-KorA, with emission filters of 500-525 nm and 650-750 nm, respectively. Protein-containing channels were passivated with BSA (0.1% w/v in PBS) for 30 min before the experiment. Kymographs were generated by single line scans between the two beads using a pixel size of 100 nm and a pixel time of 0.1 ms, resulting in a typical time per line of 22.4 ms. The confocal laser intensity at the sample was 2.2 µW for the 488 laser and 1.92 µW for the 635 laser. Experiments were performed in constant-force mode at 15 pN.

### Magnetic tweezers experiments

Magnetic tweezers experiments were performed using a homemade setup that was previously described^96,97^. Briefly, optical images of micron-sized superparamagnetic beads tethered to a glass surface by DNA substrates were acquired using a 100x oil immersion objective and a CCD camera operating at 120 Hz. Real-time image analysis allows the spatial coordinates of the beads to be determined with nm accuracy in the x, y and z directions. We controlled the stretching force of the DNA by using a step motor coupled to a pair of magnets located above the sample. The applied force is quantified from the Brownian motion of the bead and the extension of the tether, obtained by direct comparison of images taken at different focal planes^98,99^.

Magnetic tweezer experiments were performed as follows. The DNA sample containing 16xOB sites was diluted in 10 mM Tris-HCl pH 8.5, 1 mM EDTA and mixed with 1 µm diameter magnetic beads (Dynabeads, MyOne Streptavidin, Invitrogen, Carlsbad, CA) for 10 minutes. Magnetic beads were previously washed three times with PBS and resuspended in PBS/BSA at a 1:10 dilution. The DNA-bead ratio was adjusted to obtain as many single-tethered beads as possible. After incubation, we introduced the DNA-bead sample in a double PARAFILM (Sigma)-layer flow cell and allowed them to sink for 10 min to promote the binding of the DIG-labeled end of the DNA to the anti-DIG glass-coated surface. Then, a force of 5 pN was applied to remove non-attached molecules from the surface. The chamber was washed with ∼500 µl of PBS before experiments. Torsionally constrained molecules and beads containing more than a single DNA molecule were identified from their distinct rotation-extension curves and discarded for further analysis. Force-extension curves were generated by measuring the extension of the tethers at decreasing forces from 5.5 pN to 0.002 pN. The curves were first measured on naked DNA molecules and then the experiment was repeated using different concentrations of *Bacillus subtilis* ParB and KorB in a reaction buffer (10 mM Tris pH 8, 100 mM NaCl and 5 mM MgCl_2_, 1 mM DTT and 0.1 mg/mL BSA) supplemented with 2 mM CTP. Data were analyzed and plotted using Origin software.

### Construction of *PkorA* and *PkorA*:λ*P_R_* linear scaffolds

The plasmid pUC19::146-bp-*PkorA* was used as a template to amplify a 146-bp linear *PkorA* DNA scaffold using primers 5’-AGACGAAAGCCCGGTTTCCGGG-3’ and 5’-CTCCGCGCCTTGGTTGAACATAG-3’ in a PCR reaction with *Taq* DNA polymerase (Promega) at an annealing temperature of 65°C (Supplementary Table S2). The correct band was gel extracted from a 2% w/v agarose gel and eluted into TE buffer. The plasmid (pUC19-146-bp-*PkorA*:λ*P_R_*) was similarly used as a template to amplify a linear 146-bp *PkorA*:λ*P_R_* DNA scaffold (Supplementary Table S2).

### Construction of a *PkorA* linear scaffold for native mass spectrometry

The *PkorA* sequence was shortened to its minimal elements as a 100-bp DNA scaffold and synthesized as separate PAGE-purified top and bottom strand oligos (IDT) (Supplementary Table S1). The two strands were resuspended separately to 1 mM solutions in 10 mM Tris-HCl pH 8.0, 50 mM NaCl, 1 mM EDTA, pH 8.0. The strands were mixed in a 1:1 molar ratio for a 500 μM dsDNA (final concentration) and were heated to 95°C before being cooled down to 25°C in a 1°C-stepwise decrease using a Thermocycler PCR machine (Eppendorf). The resulting dsDNA was assayed by 2% w/v agarose gel electrophoresis for purity and was quantified using a Qubit (Invitrogen) dsDNA broad-range quantification kit.

### Construction of a promoter bubble DNA for half-life abortive initiation assays

An ideal promoter bubble DNA (generated with a non-complementary intervening sequence) was used as a competitor DNA for the *in vitro* half-life abortive initiation assays (Supplementary Table S2)^100^. The top and bottom strands were synthesized and annealed as described for wild-type *PkorA* 100-bp DNA scaffold used in native mass spectrometry.

### Electrophoretic mobility shift assays (EMSA)

To check for *E. coli* Eσ^70^ binding to the *PkorA* DNA scaffolds, 50 nM of *PkorA* DNA scaffold was mixed with 50 nM *E. coli* Eσ^70^ and incubated at 37°C for 5 min. In all biochemical experiments except native mass spectrometry, Eσ^70^ was assembled by incubating *E. coli* His_10_-PPX-RNAP with a fivefold molar excess of σ^70^ at 37°C for 15 min (excess σ^70^ was not purified away). The Eσ^70^-DNA sample was then loaded onto a 4.5% native Tris-Borate-EDTA (90mM TBE pH 8.3) polyacrylamide gel and run at a constant current of 15 mA for 1.5 hrs at 5-10°C in a cold room. The gel was stained for dsDNA using GelRed (Biotium).

### *In vitro* abortive initiation assay for transcription initiation repression

To assay for transcription initiation repression, we monitored levels of abortive initiation RNA products in an NTP-restricted *in vitro* promoter-based transcription reaction. The 146-bp WT *PkorA* and *PkorA*:λ*P_R_* discriminator linear DNA scaffolds were used to assay KorA and KorB-mediated repression on WT and mutant DNA-derived transcription, respectively. *E. coli* Eσ^70^ was assembled as described for EMSAs. The transcription buffer consists of 50 mM Tris-HCl pH 8.0, 10 mM MgCl_2_, 150 mM KCl, 0.1 mg/mL BSA, 1 mM DTT. Assembly of protein-DNA mixes prior to the addition of NTP mix involved a sequential incubation of DNA (10 nM), *E. coli* Eσ^70^ (50 nM) and KorAB factors (250 nM; and saturating CTP where relevant) at 37°C in 5 min incubations. NTP mix (50 μM GTP, 250 μM ApU dinucleotide, 0.05 μCi/(μL reaction volume) of α-^32^P-GTP (PerkinElmer)) was added once all factors were added and incubated for 10 min at 37°C to synthesize one abortive RNA band (5’-ApUpG*-3’; +1 to +3 RNA). Reactions were quenched using a 2X STOP buffer (45 mM TBE pH 8.3, 8 M urea, 30 mM EDTA pH 8.0, 0.05% w/v bromophenol blue, 0.05% w/v xylene cyanol). Reaction samples were analyzed on a 23% denaturing PAGE (19:1 acrylamide:bis-acrylamide, 90 mM TBE pH 8.3, 6 M urea) for 1.5-2 hrs at a constant voltage of 800 V, and the gel was exposed on a storage phosphor screen for 1-2 hrs and imaged using a Typhoon Phosphorimager (Cytiva). Band intensities were quantified on ImageJ^101^, with measured values subtracted of background and normalized to an *E.coli* Eσ^70^-DNA only control (no repression) for values to be averaged amongst replicates (*n* of at least 3 in all repression conditions). Repression as percentage values was calculated as (1-normalized intensity) × 100%, graphed and statistically analyzed in GraphPad Prism using unpaired Welch’s t-tests.

### *In vitro* abortive initiation assay for half-life estimation

To estimate half-lives of *E. coli* Eσ^70^-open promoter complexes on both the WT and mutant *PkorA* DNA scaffolds, we adapted the *in vitro* abortive initiation assay. Eσ^70^ was incubated in transcription buffer for 5 mins at 37°C before adding promoter DNA scaffold and incubated for another 15 mins at 37°C. This formed a master mix to take out aliquots at desired timepoints (*t* = 0, 1, 2, 5, 10, 30, 45, 60 min), to which NTP mix was added to start transcription of abortive RNAs. Timepoints began when competitor DNA (100 nM final concentration) was added to the master mix (*t* = 0 s was transcribed as a separate reaction) and was incubated at 37°C throughout. In this way, abortive RNA production is a proxy for the proportion of open promoter complexes remaining as time goes on, as Eσ^70^ that dissociates from the promoter binds to the ideal bubble competitor DNA and does not easily dissociate (re-binding to the *PkorA* scaffold is thereby negligible). Quantification of abortive RNA bands was the same as performed for the repression assays. Plots of normalized band intensities against time resulted in a double exponential curve, with the slow decaying component (i.e. basal noise-level signal) dominating at timepoints with low fractions of competitor-resistant open promoter complexes. Due to the rapid decline in WT *PkorA* open complexes, a correction was applied by subtracting values that occur at *t* ≥ 30 min to remove the slow decaying component and derive the true half-life from the fast decay as a single exponential decay curve. This correction was not applied to *PkorA*:λ*P_R_* discriminator open complex values as the timepoints measured did not decline to noise-level signal.

The plots were presented on a semi-log scale, with the fraction of competitor-resistant open promoter complexes on a log_10_ scale and time on a linear scale, with single exponential decay trendlines fitted (WT *PkorA* with R^2^ = 0.7637; *PkorA*:λ*P_R_* discriminator with R^2^ = 0.9934). Analysis and plotting were performed using Microsoft Excel and GraphPad Prism.

### Preparation of *E. coli* Eσ^70^ holoenzyme for native mass spectrometry

Eσ^70^ was formed by mixing purified His_10_-PPX-RNAP and 2.5-fold molar excess of σ^70^ and incubated for 15 mins at 37°C. Eσ^70^ was buffer exchanged into 20 mM Tris-HCl pH 8.0, 500 mM NaCl, 5 mM DTT (to remove most glycerol from protein storage buffer) by centrifugal filtration and purified on a Superose 6 Increase 10/300 GL column (Cytiva) pre-equilibrated in 50 mM Tris-HCl pH 8.0, 150 mM KCl, 10 mM MgCl_2_, 1 mM DTT. The eluted Eσ^70^ was concentrated to ∼10-12 mg/mL (∼21-26 μM) by centrifugal filtration. Purified Eσ^70^ was supplemented with glycerol to a final concentration of 20% w/v, flash-frozen in liquid nitrogen, and stored at -80°C.

### Native mass spectrometry (nMS) analysis

Frozen samples were thawed and reconstituted in various combinations. Sample concentrations for conditions without Eσ^70^ used 2 μM WT *PkorA* DNA and 5 μM KorA/B factors. For conditions with Eσ^70^, the sample contained 5 μM Eσ^70^, 5.5 μM WT *PkorA* DNA and 2.5-fold molar excess (12.5 μM) of KorA/B factors. For KorB samples with CTP, 0.5 mM CTP was used to ensure saturation of KorB.

For nMS analysis, samples were buffer-exchanged into either 150 mM, 300 mM or 500 mM ammonium acetate pH 7.5, 0.01% Tween-20 using Zeba microspin desalting columns with a 7-kDa or 40-kDa molecular weight cutoff (Thermo Scientific). A 2-3 µL aliquot of the sample was then loaded into a gold-coated quartz capillary tip that was prepared in-house and then electrosprayed into an Exactive Plus with extended mass range (EMR) instrument (Thermo Fisher Scientific) with a static direct infusion nanospray source^102^. The typical MS parameters for all samples included: spray voltage, 1.20-1.22 kV; capillary temperature, 125-150 °C; in-source dissociation, 0-10 V; S-lens RF level, 200; resolving power, 8,750 at *m/z* of 200; AGC target, 1 x 10^6^; maximum injection time, 200 ms; number of microscans, 5; total number of scans, at least 100. Additional specific nMS parameters were injection flatapole, 8 V; interflatapole, 4 V; bent flatapole, 4 V; high energy collision dissociation (HCD), 150-200 V; ultrahigh vacuum pressure, 5.2-5.8 × 10^−10^ mbar. Mass calibration in positive EMR mode was performed using cesium iodide. For data processing, the collected nMS spectra were visualized using Thermo Xcalibur Qual Browser (v. 4.2.47). Spectral deconvolution was performed using UniDec v. 4.2.0^83,103^ with the following general parameters: for background subtraction (if applied), subtract curve 10; smooth charge state distribution, enabled; peak shape function, Gaussian; Degree of Softmax distribution (beta parameter): 10-20.

The expected masses for the component proteins of the Eσ^70^ include α: 36,511.7 Da, β: 150,632.2 Da, β’: 158,008.1 Da (includes one Mg^2+^ and two Zn^2+^ ions), ω (lost N-terminal methionine): 10,105.4 Da and σ^70^: 70,263.3 Da. The expected masses for the Kor proteins and the DNA scaffold used are KorA monomer: 12,825.6 Da, KorB (lost N-terminal methionine): 40,399.6 Da and *PkorA* dsDNA: 61,667.2 Da. The observed mass deviations (calculated as the % difference between the measured and expected masses relative to the expected mass) ranged from 0.001 – 0.3% with a typical mass deviation of 0.12%.

## Supporting information

PDB and validation reports

## ACKNOWLEDGEMENTS

We thank Miloš Tišma and Cees Dekker for preliminary experiments, Stephan Gruber and members of our laboratories for helpful comments, and Mark Dillingham for providing purified *B. subtilis* ParB. This work is supported by the Royal Society University Fellowship Renewal URF\R\201020 and the Lister Institute fellowship (T.B.K.L), by the Wellcome Trust Investigator grant 221776/Z/2/Z (T.B.K.L and T.C.M), by the BBSRC funded Institute Strategic Program Harnessing Biosynthesis for Sustainable Food and Health (HBio) (BB/X01097X/1), and by grants from the NIH to S.A.D. (R35 GM118130), E.A.C. (R01 GM14450), and B.T.C (P41 GM109824 and P41 GM103314). Work in the F.M.H laboratory was supported by grants PID2020-112998GB-I00, funded by Ministerio de Ciencia e Innovación (MICINN)/Agencia Estatal de Investigación (AEI/10.13039/501100011033)_FEDER, EU, and co-funded by the European Regional Development Fund (ERDF), and grant EUREXCEL Ref. 951214, funded by CSIC. We thank Diamond Light Source for access to beamlines I04-1 and I03 under proposal MX25108.

---

**Figure S1.**
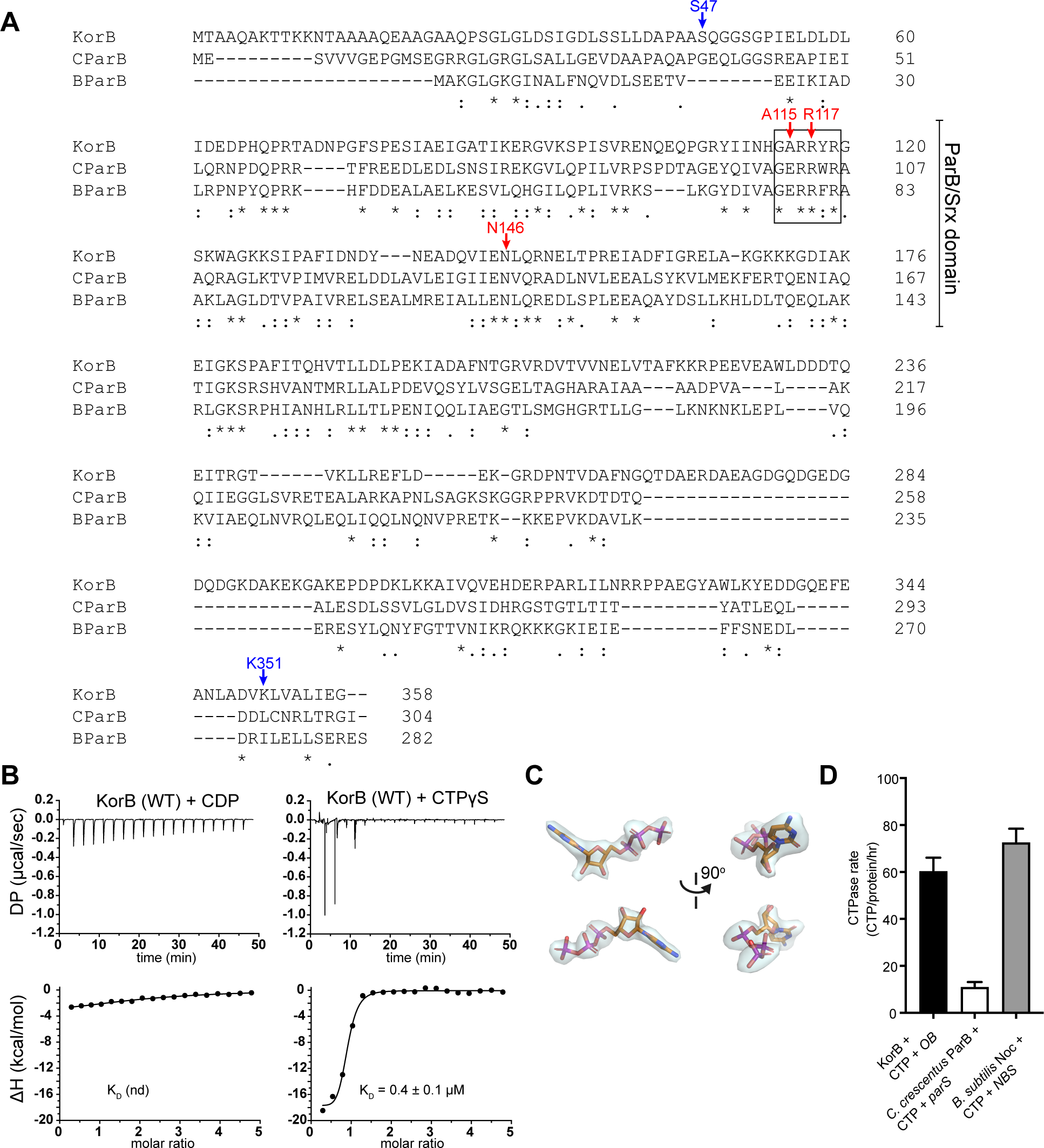
KorB is a CTPase enzyme. **(A)** A sequence alignment between KorB, *Caulobacter crescentus* ParB (CParB), and *Bacillus subtilis* ParB (BParB). Residues R117, N146, and A115 (red) of KorB are indicated on its amino acid sequence. The position of the P-motif 2 (GARRYR for KorB and GERRxR for canonical ParB), and the N-terminal ParB/Srx-like domain are also indicated on the sequence alignment. Residues S47 or K351C (blue) were substituted by cysteine to generate KorB (S47C) and KorB (K351C) variants which was subsequently used in BMOE crosslinking assays (Figure 2A and S2A). **(B)** Analysis of the interaction of KorB with CDP and CTPɣS by ITC. Each experiment was duplicated. Regression curves were fitted, and binding affinities (K_D_) were shown. **(C)** An omit difference map for CTPγS was calculated after removing the ligands from the final structure and re-refining to convergence at 2.3 Å resolution. Shown are orthogonal views of the two ligands from the AB dimer only, together with their associated omit density displayed as a semi-transparent cyan surface contoured at 2.0 σ. It was not possible to unambiguously assign the positions of the ligand sulfur atoms, thus they were modelled as CTP molecules. **(D)** CTP hydrolysis rates of KorB, *C. crescentus* ParB, and *B. subtilis* Noc were measured by continuous detection of released inorganic phosphates (see Methods). CTPase rates were measured at 1 mM concentrations of CTP and in the presence of 0.5 µM of cognate DNA duplexes. Experiments were triplicated, and the SDs of the CTPase rates were presented.

**Figure S2.**
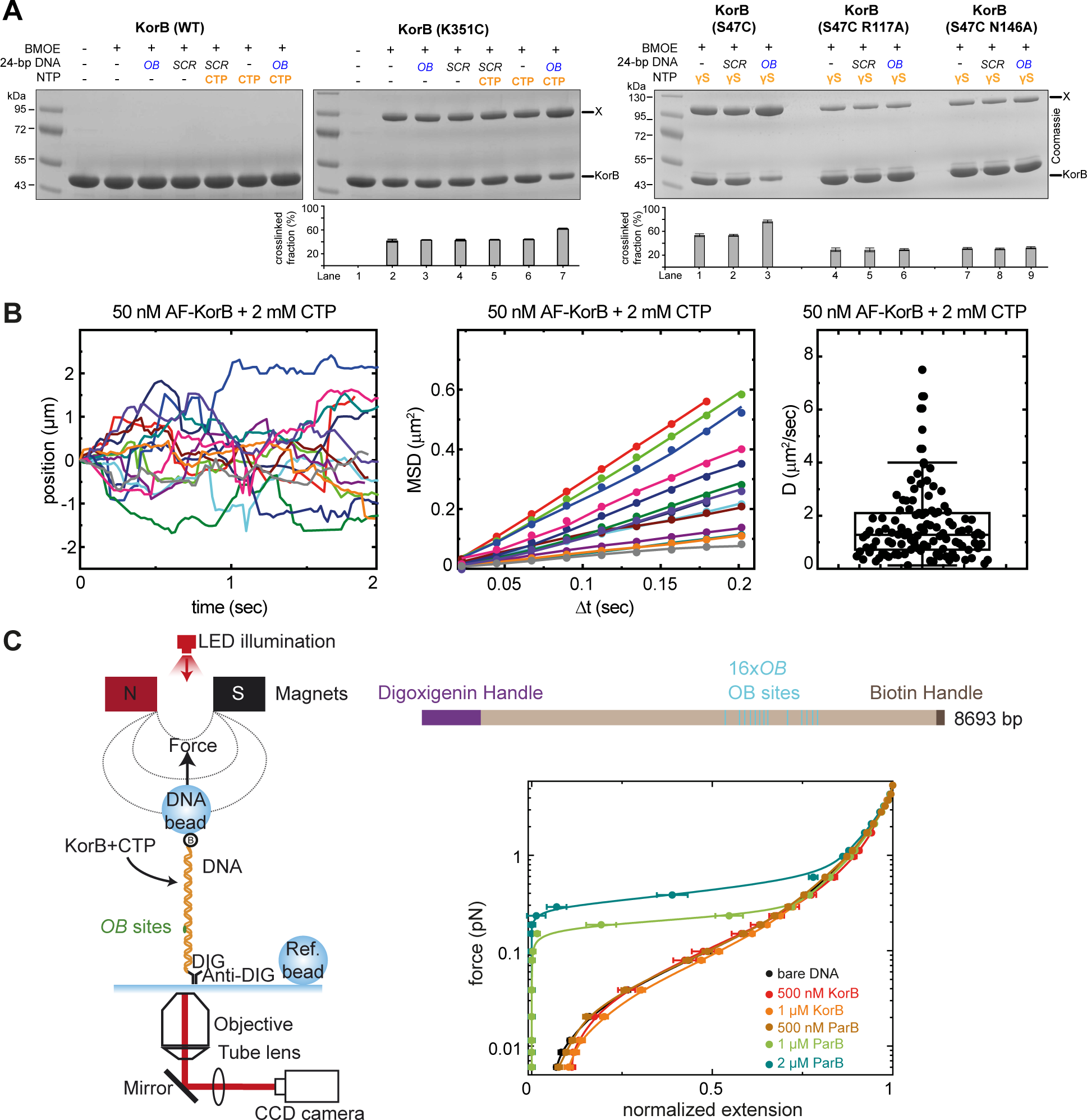
CTP enables KorB diffusion on *OB*-containing DNA, but KorB-CTP does not condense *OB* DNA *in vitro*. **(A)** (left panel) KorB is cysteine-less and did not crosslink in the presence of BMOE. (middle panel) The CTD of KorB is likely a constant dimerization domain. SDS-PAGE analysis of BMOE crosslinking products of 8 µM KorB (K351C) (and variants) ± 0.5 µM 24 bp *OB*/scrambled *OB* DNA ± 1 mM CTP. (right panel) a non-hydrolyzable analog CTPɣS readily promotes crosslinking of KorB S47C regardless of DNA. X indicates a crosslinked form of KorB. Quantification of the crosslinked fraction is shown below each representative image. Error bars represent SEM from three replicates. **(B)** Determination of the diffusion constant of KorB. (left panel) Representative KorB trajectories measured on the DNA (n = 111). (middle panel) Mean squared displacement (MSD) of KorB trajectories for different time intervals (Δt). (right panel) The diffusion constant of KorB (1.61 ± 0.12 µm^2^/s, mean ± SEM) was calculated as half of the slope of the linear fit of MSD versus Δt. **(C)** KorB did not condense a DNA containing 16x *OB* sites in the presence of CTP. (left panel) Cartoon of the basic magnetic tweezers (MT) components and the layout of the experiment, and a schematic representation of a 16x *OB* DNA. The position of the *OB* sites is represented to scale. (right panel) Average force-extension curves of bare 16x *OB* DNA molecules (n = 56) and in the presence of different concentrations of KorB + 2 mM CTP (500 nM, n = 11, and 1 µM, n= 13) or *B. subtilis* ParB + 2 mM CTP (500 nM, n = 11, 1 µM, n= 21 and 2 µM, n =17).

**Figure S3.**
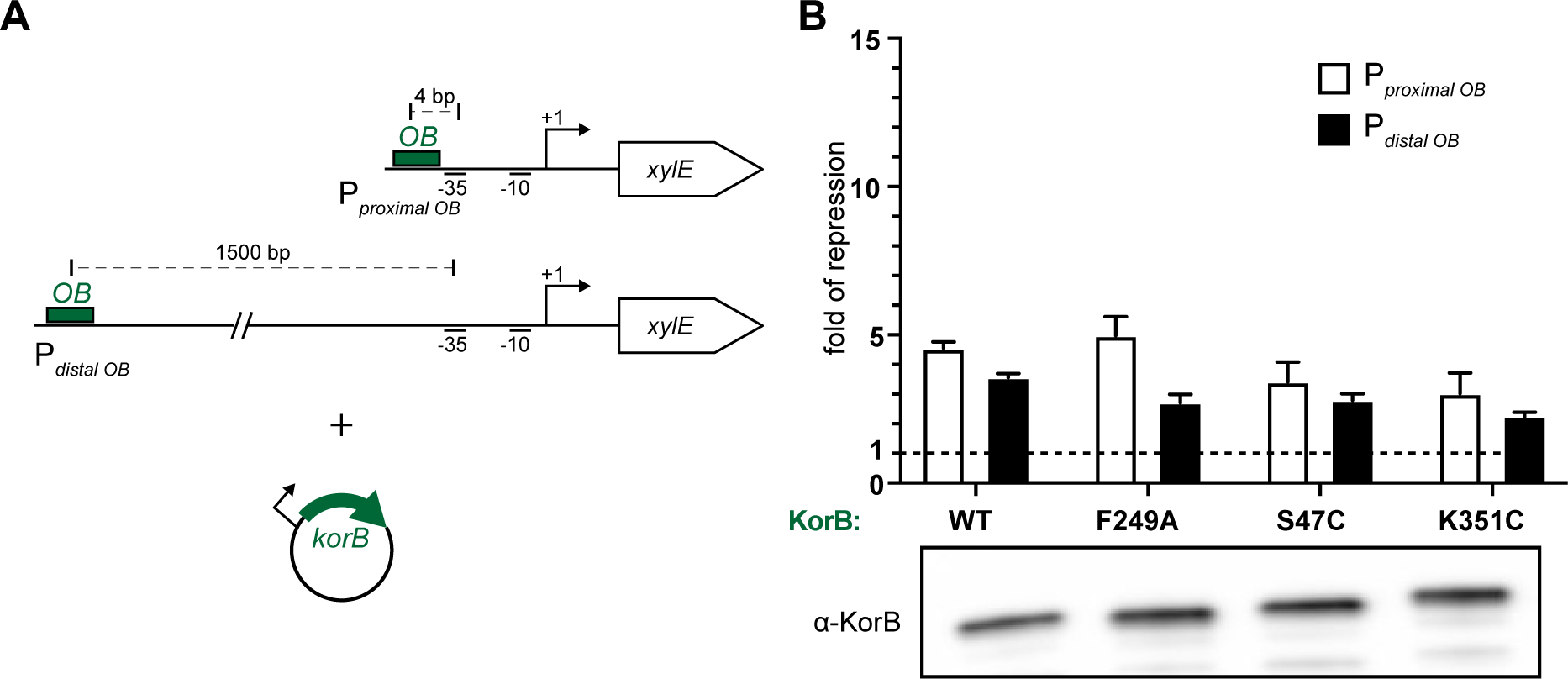
Substitutions S47C, K351C or F249A on KorB did not impact KorB ability to repress *OB*-proximal or distal promoters. **(A)** Schematic diagrams of promoter-*xylE* reporter constructs. For the *OB*-proximal promoter, *OB* is positioned 4 bp upstream of the -35 core promoter element. For the *OB*-distal promoter, *OB* is 1500 bp upstream of the -35 element. *E. coli* cells were co-transformed with the reporter plasmid and an inducible plasmid expressing KorB (WT or variants) or empty plasmid as a negative control. **(B)** Substitutions S47C, K351C or F249A on KorB did not drastically affect transcriptional repression at an *OB*-proximal or *OB*-distal promoters. Values shown are fold of repression, a ratio of XylE activities from cells co-harboring a reporter plasmid and KorB-expressing plasmid to that of cells co-harboring a reporter plasmid and an empty plasmid (KorB-minus control). Experiments were triplicated, and the SDs were presented. An α-KorB immunoblot from lysates of cells used in the experiments is also shown below.

**Figure S4.**
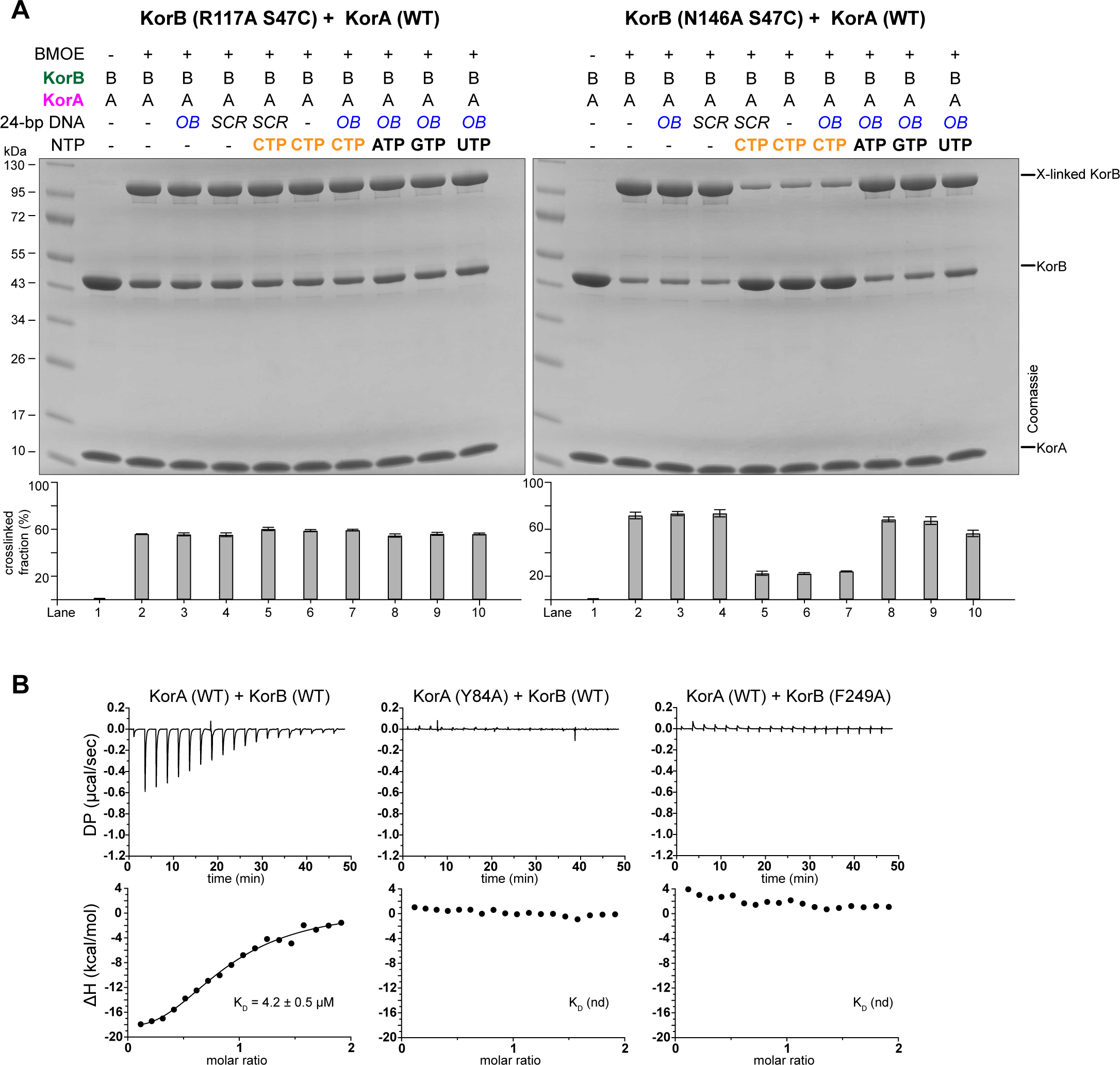
KorA interacts with KorB to promote the N-engagement of KorB. **(A)** KorA can promote/lock clamp-defective KorB (R117A) and (N146A) variants to N-engage. SDS-PAGE analysis of BMOE crosslinking products of 8 µM KorB (S47C R117A) and KorB (S47C N146A) ± KorA± 0.5 µM 24 bp *OB*/scrambled *OB* DNA ± 1 mM CTP. X indicates a crosslinked form of KorB (S47C R117A) or (S47C N146A). Quantification of the crosslinked fraction is shown below each representative image. Error bars represent SEM from three replicates. **(B)** Substitutions Y84A on KorA or F249A on KorB eliminated KorA-KorB interaction. Analysis of the interaction of KorB (WT or variant) with KorA (WT or variant) by ITC. Each experiment was duplicated. Regression curves were fitted, and binding affinities (K_D_) were shown.

**Figure S5.**
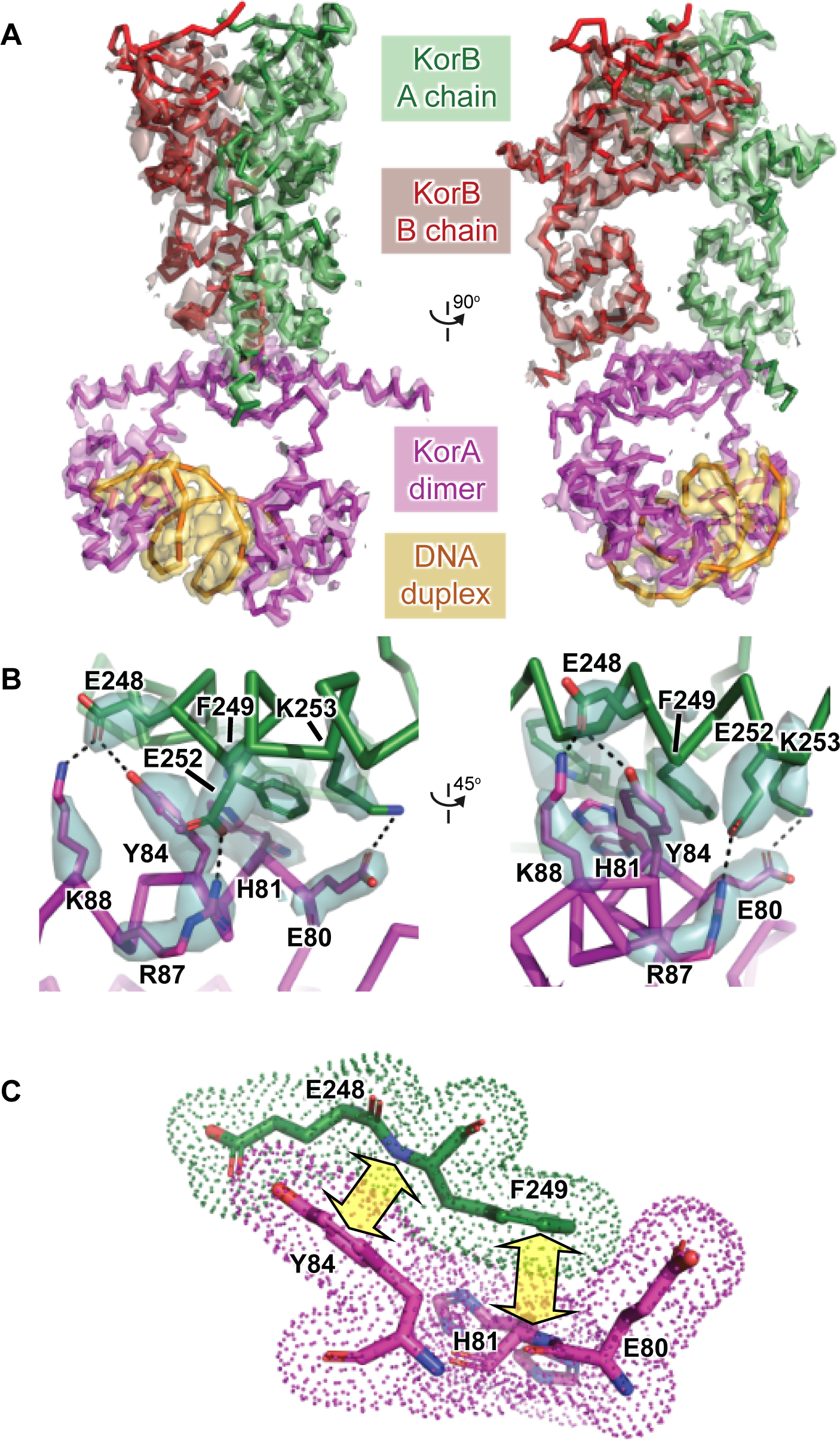
Structure of the KorA-KorB-DNA ternary complex. The crystal structure of the KorA-KorB-DNA complex was determined at 2.7 Å resolution. **(A)** A series of omit difference maps were calculated by separately removing parts of the final structure and re-refining to convergence at 2.7 Å resolution. Maps for KorB chain A (green), KorB chain B (red), the KorA dimer (magenta) and the DNA duplex (orange) are displayed as semi-transparent surfaces on a color-coded backbone trace of the structure, contoured at 2.0 σ and shown as orthogonal views. **(B)** Close-up of a KorB-KorA interface with only the side-chains of key residues displayed. Also shown is omit difference density (semi-transparent cyan surface, contoured at 2.0 σ) calculated for the model after removal of these side-chains and re-refining. **(C)** Further detail on the KorB-KorA interface with color-coded van der Waals dots illustrating intimate contact. In addition to the hydrogen bonds highlighted in panel B, the aromatic ring of Tyr84 in KorA makes pi-pi interactions with the Glu248-Phe249 peptide bond of KorB, which is reciprocated by the aromatic ring of Phe249 in KorB making pi-pi interactions with the Glu80-His81 peptide bond of KorB. These interactions are indicated by the pale yellow double-headed arrows.

**Figure S6.**
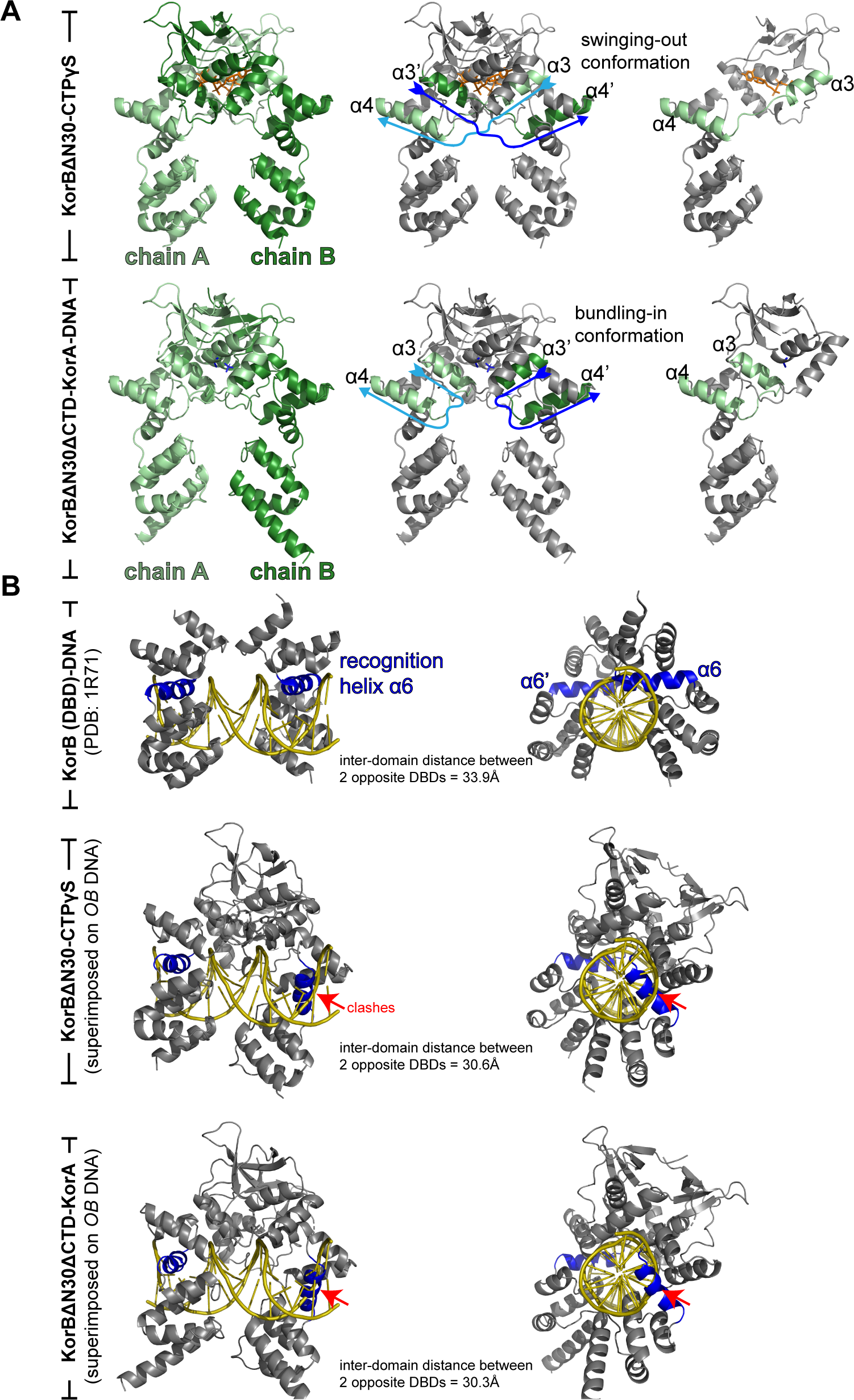
KorB adopts a closed-clamp conformation in the co-crystal structures of KorBΔN30ΔCTD-CTPγS complex and KorBΔN30ΔCTD-KorA-DNA complex, and the closed-clamp conformation is incompatible with site-specific DNA-binding at the *OB* site. **(A)** Structures of closed-clamp KorBΔN30ΔCTD dimers with the pair of helices α3 and α4 colored differently (light green and dark green) to highlight the “swinging-out” or the “bundling-in” conformations. Light blue and dark blue arrows indicate the direction of the α3-α4 pair or the α3’-α4’ pair from the opposite subunit. **(B)** The structure of a nucleotide-bound or KorA-bound KorBΔN30ΔCTD is incompatible with specific *OB* binding at the DNA-binding domain (DBD). Superimposing the KorBΔN30ΔCTD-CTPɣS and KorBΔN30ΔCTD-KorA-DNA structures onto *OB* DNA (from the previously published KorB(DBD only)-*OB* co-crystal structure, PDB: 1R71) shows DNA-recognition helices (α6 and α6′, dark blue) positioning closer and away from the two consecutive major grooves of *OB* DNA (yellow), and helices α6 and α6′–α9′ at the DBD (red arrows) clashing with *OB* DNA.

**Figure S7.**
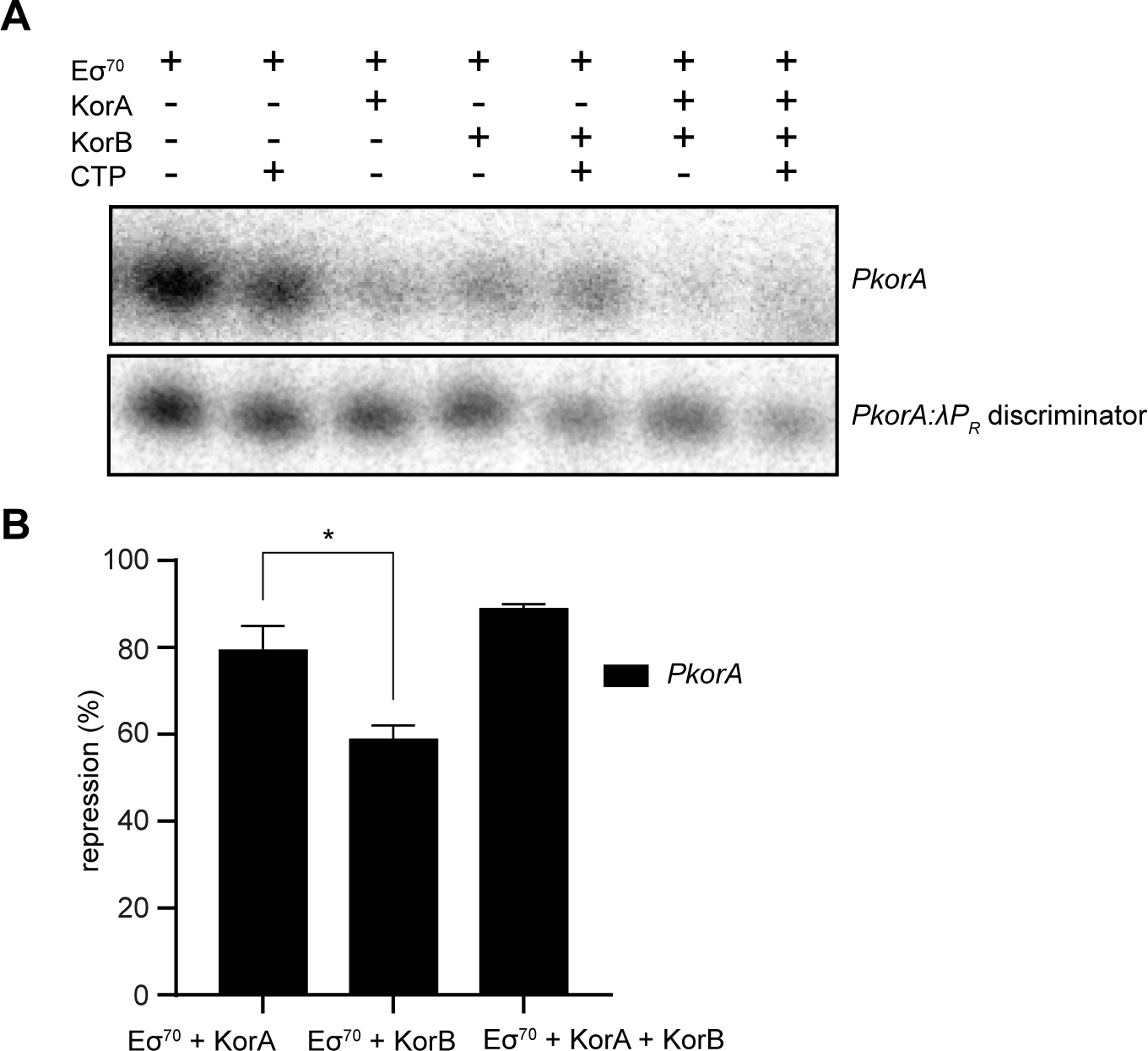
KorA and KorB are transcriptional repressors that exclude *Eco* RNAP from promoters. **(A)** Representative urea-PAGE gel closeup on the abortive RNA product (5’-ApUpG*-3’; *radiolabelled on guanosine alpha-phosphate) transcribed in *in vitro* abortive initiation assays on WT *PkorA* and the *PkorA*:λ*P_R_* discriminator mutant linear DNA scaffolds with different mixes of Eσ^70^ and/or fivefold excess KorA, fivefold excess KorB and/or saturating CTP. **(B)** WT *PkorA in vitro* transcription repression of *E. coli* RNAP:σ^70^ holoenzyme (*Eco* Eσ^70^) in the presence of fivefold excess KorA and/or KorB. All values are normalized to holoenzyme only control as mean values ± SEM. P-value was calculated by an unpaired Welch’s t-test; * p ≤ 0.05.

