## Supplementary tables for "Molecular switching of a DNA-sliding clamp to a repressor mediates long-range gene silencing"

### Supplementary Table S1. Strains, plasmids, and DNA oligonucleotides used in this work

| Strains | Strains/description | Source |
|---|---|---|
| DH5α | *E. coli* host for DNA cloning and propagation of plasmid | Lab collection |
| *E. coli* Rosetta (BL21 DE3) | *E. coli* host for protein overexpression from an IPTG-inducible T7 promoter F- *ompT hsdS$_B$*($r_B$- $m_B$-) *gal dcm* (DE3) pRARE (chloramphenicol$^R$) | Merck |
| | BL21 Rosetta (DE3) + various pET21b-based protein overexpression vectors (see the plasmid list for the complete collection of protein overexpression plasmids) | This study |
| | DH5α + various combinations of inducible expression vectors for *xylE* experiments | This study |

| Plasmids | Description | Replicon | Source |
|---|---|---|---|
| TCMV0001 | pET21b overexpression of C-terminally His$_6$-tagged RK2 KorB, carbenicillin$^R$ | pMB1 | This study |
| TCMV0003 | pET21b overexpression of C-terminally His$_6$-tagged RK2 KorB truncated to residues 30-252, carbenicillin$^R$ | pMB1 | This study |
| TCMV0007 | pET21b overexpression of C-terminally His$_6$-tagged RK2 KorB S47C, carbenicillin$^R$ | pMB1 | This study |
| TCMV0008 | pET21b overexpression of C-terminally His$_6$-tagged RK2 KorB K351C, carbenicillin$^R$ | pMB1 | This study |
| TCMV0009 | pET21b overexpression of C-terminally His$_6$-tagged RK2 KorB S47C K351C, carbenicillin$^R$ | pMB1 | This study |
| TCMV0018 | pET21b overexpression of C-terminally His$_6$-tagged RK2 KorB R117A, carbenicillin$^R$ | pMB1 | This study |
| TCMV0024 | pET21b overexpression of C-terminally His$_6$-tagged RK2 KorB N146A, carbenicillin$^R$ | pMB1 | This study |
| TCMV0028 | pET21b overexpression of C-terminally His$_6$-tagged RK2 KorB S47C R117A, carbenicillin$^R$ | pMB1 | This study |
| TCMV0032 | pET21b overexpression of C-terminally His$_6$-tagged RK2 KorB S47C N146A, carbenicillin$^R$ | pMB1 | This study |
| RK2 | RK2 plasmid, kanamycin$^R$, tetracycline$^R$, penicillin$^R$ | IncP$\alpha$ | Gift from Rahmi Lale |
| TCMV0044 | pET21b overexpression of C-terminally His$_6$-tagged RK2 KorA, carbenicillin$^R$ | pMB1 | This study |
| TCMV0063 | pET21b overexpression of C-terminally His$_6$-tagged RK2 KorA Y84A, carbenicillin$^R$ | pMB1 | This study |
| TCMV0070 | pET21b overexpression of C-terminally His$_6$-tagged RK2 KorB S47C F249A, carbenicillin$^R$ | pMB1 | This study |
| pPT01 | Empty *xylE* promoter fusion vector, kanamycin$^R$ | pSC101 | Gift from Christopher Thomas |
| pLB40 | pPT01 *trbBp-xylE* promoter fusion vector - *OB* cloned in proximal position, kanamycin$^R$ | pSC101 | Gift from Christopher Thomas |
| pLB105 | pPT01 *trbBp-xylE* promoter fusion vector - *OB* cloned in distal position, kanamycin$^R$ | pSC101 | Gift from Christopher Thomas |
| pDM1.2 | Empty *lacIp-tacp* expression vector, streptomycin$^R$ | IncQ | Gift from Christopher Thomas |
| TCMV0083 | pDM1.2 *lacIp-tacp* inducible RK2 KorA Y84A expression vector, streptomycin$^R$ | IncQ | This study |
| TCMV0084 | pDM1.2 *lacIp-tacp* inducible RK2 KorA expression vector, streptomycin$^R$ | IncQ | This study |
| pBAD33 | Empty arabinose inducible promoter vector, chloramphenicol$^R$ | p15A | [1] |

| | | | |
|---|---|---|---|
| TCMV0086 | pBAD33 arabinose inducible promoter vector expressing RK2 KorB, chloramphenicol$^R$ | p15A | This study |
| TCMV0087 | pPT01 *korAp-xylE* promoter fusion vector - *OB* cloned in proximal position, kanamycin$^R$ | pSC101 | This study |
| TCMV0091 | pBAD33 arabinose inducible promoter vector expressing RK2 KorB ΔCTD, chloramphenicol$^R$ | p15A | This study |
| TCMV0093 | pBAD33 arabinose inducible promoter vector expressing RK2 KorB S47C, chloramphenicol$^R$ | p15A | This study |
| TCMV0094 | pBAD33 arabinose inducible promoter vector expressing RK2 KorB K351C, chloramphenicol$^R$ | p15A | This study |
| TCMV0095 | pBAD33 arabinose inducible promoter vector expressing RK2 KorB F249A, chloramphenicol$^R$ | p15A | This study |
| TCMV0105 | pBAD33 arabinose inducible promoter vector expressing RK2 KorB R117A, chloramphenicol$^R$ | p15A | This study |
| TCMV0109 | pBAD33 arabinose inducible promoter vector expressing RK2 KorB N146A, chloramphenicol$^R$ | p15A | This study |
| TCMV0117 | pET21b overexpression of C-terminally $His_6$-tagged RK2 KorA with C-terminal cysteine added, carbenicillin$^R$ | pMB1 | This study |
| TCMV0118 | pET21b overexpression of C-terminally $His_6$-tagged RK2 KorA A6C, carbenicillin$^R$ | pMB1 | This study |
| TCMV0119 | pET21b overexpression of C-terminally $His_6$-tagged RK2 KorB F249A, carbenicillin$^R$ | pMB1 | This study |
| TCMV0120 | pPT01 *korAp-xylE* promoter fusion vector - *OB* cloned in distal position, kanamycin$^R$ | pSC101 | This study |
| TCMV0121 | pPT01 *korAp-xylE* promoter fusion vector - scrambled *OB* cloned in distal position, kanamycin$^R$ | pSC101 | This study |
| pUC19 | High-copy-number plasmid for general cloning purposes, carbenicillin$^R$ | pMB1 | Lab collection |
| pUC19_v1 | pUC19 containing 1x *OB* and 1x *OA* separated by 1016 bp | pMB1 | This study |
| pSP73-JY0 | template for PCR to generate handles for magnetic tweezers experiments | pMB1 | [2] |
| pVS11 | pEcrpoABC(-XH)Z), overexpressing each subunit of *E. coli* RNAP (full-length α, β, ω) as well as β'-PPX-His10 | pBR322/f1 | [1,2] |
| pACYCDuet-1::*E. coli rpoZ* | Overexpression of *E. coli* RpoZ, chloramphenicol$^R$ | p15A | [1,2] |
| pSAD1403 | Overexpression of *E. coli* His10-SUMO-σ70, kanamycin$^R$ | pBR322/f1 | [3] |

| Oligos | Sequence | Description | Source |
|---|---|---|---|
| TCMP0019 | gggatattttagcggctaaaagga | *OB*-containing sequence for crosslinking and EnzCheck assays | This study |
| TCMP0020 | tccttttagccgctaaaatatccc | *OB*-containing sequence for crosslinking and EnzCheck assays | This study |
| TCMP0021 | gggatataactgttatgagcagga | *SCR*-containing sequence for crosslinking and EnzCheck assays | This study |
| TCMP0022 | tcctgctcataacagttatatccc | *SCR*-containing sequence for crosslinking and EnzCheck assays | This study |
| TCMP0054 | tgtttagctaaaca | *OA*-containing sequence for crystallisation of KorBΔN30ΔCTD-KorA-*OA* | This study |
| Biotin-M13-F | Biotin-cgccagggttttcccagtcacgac | For amplification and biotinylation of 175bp dsDNA for BLI | Lab collection |
| Biotin-M13-R | Biotin-atggtcatagctgtttcct | For amplification and biotinylation of 175bp dsDNA for BLI | Lab collection |

| Fragment | Oligonucleotide | Sequence |
|---|---|---|

| | | |
|---|---|---|
| Annealed oligonucleotides with 2 *OB* sites | 279.up 2OB PshAI-SnaBI | CTGACTACGTATTGGTCAGGATTTTAGCGGCTAAA AGGGTATGAGAGCTTGGGACGCTCGTCGCGGTTG GGGACTCTATTTTAGCGGCTAAAAGTGCCGTATTT GCAGTACCAG |
| | 280.down 2OB PshAI-SnaBI | CTGGTACTGCAAATACGGCACTTTTAGCCGCTAAA ATAGAGTCCCCAACCGCGACGAGCGTCCCAAGCT CTCATACCCTTTTAGCCGCTAAAATCCTGACCAATA CGTAGTCAG |
| MT 8x *OB* PCR fragment | 276.F 132 SpeI | GCGTAAGTACTAGTCTCGAAGACTAATCCGGCGG |
| | 281.R colPCR y seq 134 XhoI | GCGTAAGTCTCGAGCGCCAGAATGTGTCAGAGAC |
| C-trap 1x *OA* PCR fragment | 283.M13 forw | GTAAAACGACGGCCAGT |
| | 284.R KpnI 133 cloning | GCGTAAGTGGTACCCGCTGGAGAAGCCATGCG |
| C-trap 8x *OB* PCR fragment | 276.F 132 SpeI | GCGTAAGTACTAGTCTCGAAGACTAATCCGGCGG |
| | 285.M13 rev | CAGGAAACAGCTATGAC |
| MT DIG handle | 57.FMH_F2_BamHI-ApaI | GCGTAAGTGGATCCGGGCCCCGACTCACTATAGG GAGAC<br>CGGC |
| | JOE_R1 | AGTAAGCGCCGTCAGACCAG |
| MT BIO handle | 57.FMH_F2_BamHI-ApaI | GCGTAAGTGGATCCGGGCCCCGACTCACTATAGG GAGAC<br>CGGC |
| | 209.BsrGI 71short handle | CGATAACCAACTGGCGATG |
| C-trap BIO handle | 235.F-JY0 PspOMI no XhoI | GCGTAAGCGGGCCCGCCATTTAAGGCGTTATCCC C |
| | JOE_R1 | AGTAAGCGCCGTCAGACCAG |

**Supplementary Table S2. dsDNA constructs used in this work.**

*OA* site is highlighted in magenta and *OB* sites in direct or inverted orientation are highlighted in green.

| Fragment | Sequence |
|---|---|
| *PkorA*<br>(146 bp) | AGACGAAAGCCCGGTTTCCGGGCTTTTGTTTTGTTACGCCAAGGACGAGTTTTAGCGGC TAAAGGTGTTGACGTGCGAGAAATGTTTAGCTAAACTTCTCTCATGTGCTGGCGGCTGTC ACCGCTATGTTCAACCAAGGCGCGGAG |
| *PkorA:λP$_R$*<br>(146 bp) | AGACGAAAGCCCGGTTTCCGGGCTTTTGTTTTGTTACGCCAAGGACGAGTTTTAGCGGC TAAAGGTGTTGACGTGCGAGAAATGTTTAGCTAAACTGGTTGCATGTGCTGGCGGCTGT CACCGCTATGTTCAACCAAGGCGCGGAG |
| *PkorA*<br>(100 bp for nMS) | GTTACGCCAAGGACGAGTTTTAGCGGCTAAAGGTGTTGACGTGCGAGAAATGTTTAGCT AAACTTCTCTCATGTGCTGGCGGCTGTCACCGCTATGTTCA |
| promoter bubble DNA competitor | **NT-DNA strand:**<br>CGAAATTTTTTTTTGAAAAGTACTTGACAAAAGTGTTAAATTGTGCTATACTGGGATGGTT ACGTACACAGAATTCGG<br>**T-DNA strand (different to create a non-complementary bubble region):** |

| | |
|---|---|
| | GCTTTAAAAAAAAACTTTTCATGAACTGTTTTCACAATTTAACACGAAATGTGGGATGGAA<br>TGCATGTGTCTTAAGCC |
| MT central fragment<br>(8693 bp) | GGCCGCCCGCCATCGGGATTTTCACCACATCAATGGGGTAACGGTTTTTCCCAGCCACA<br>CGCTGCATGACATGCCACCGGCCATTTTTCAGTTGCTGAATAAACGCGCCGGGAATACG<br>ACGGTTACCCACCACAAGCACGCTGCCGCCACCTTTCAGGGATGAACGCTGCCCCTTTT<br>TACGACGCCTGCGGCGCGAAAGGACAACCCGCGCATTACCCAGCTTGATTACGGGCAA<br>ATCCCCCCGGTTAACTTTGATTCTGGCCTGCGGATTTTTGACCGTGGCCCTTTTCAGCCT<br>GGCCCTTTCCTTTACCAGTTTCCGGCGTACCTTTGTCTCACGGGCAACCTGTGACGCCG<br>ACTGCGATATCGCGGATGAAGCAACGCGGTTAATGGCCATTGCGGCGGCACCAGGCAC<br>CGCCGTTTTGCTGATACGGCTGAGGTTTTCAACGGCCTGCTCAAGACCTTTTATGGCCA<br>TACATCCCCCTTTCAGCGGCGACGGTTAACGGCAGGCGGTACGCCCCGTCCAAGCCAG<br>AGATGACAACTTCCGCCATCATCCGGCGAAACCCGATCTACCCAGAAATTTTCCTCACC<br>GATGGTCAGCGTGTCTCCACGCCGCAGCTGCCGCACCTCATCAGTCCGGACAAACAGG<br>GACGGGCTGGAGCCTTCAACGCGCACGCCCTGTCCGGCATAGCTGATATTTTCAGGGT<br>CATCAAAAACACCACGTATCACCGCACCTGACTGCTCACCGGATGTAATGGTGGCTGAC<br>GTTCCCATGTACCCGCGTATCGTTTCATCGGCGCGGGCAATGGCAGCATCGAACAGGTT<br>ATCGAAATCAGCCACAGCGCCTCCCGTTATTGCATTCTGGCCAGGCCGCGCTCTGTCAT<br>TTCGGCTGCCACACCGGCAGAGACACGAAACGCCGTTCCCGGCAGCACAAATGCCACA<br>GGTTCATCCCGCGTGGCGTGAAGTGCATCAGTATGCAGCTTCACCAGTGCCACGACCG<br>TGACCAGTTCAGACGTATCCAGAATCACGGTATCCGGCTGCGCTGATCCCACCTCATTT<br>TCATGTCCGGTCAGCACATTTTCCCGGCTGAGAGGGGTGTCCTGACCGGCAGTTTCATC<br>CGTGTCATCAAGCTCCTCTTTCAGCTCTGCCACACGGAGCGCCAGTTCTTCTTTCGTCC<br>CCGTCAGGCTGACATCACGGTTCAGTTGTTCACCCAGCGAGCGGAGACGGGCAATCAG<br>TTCATCTTTCGTCATGGACTCCTCCACAGAGAAACAATGGCCCCGAAGGGCCATGATTA<br>CGCCAGTTGTACGGACACGAACTCATCAGGGTCAGCCAGCAGCATCAGCGGTGCTGAC<br>TGAATCATGGTGAACTCACGCGCCGGATCGCCGGTGGTCACCCAGTTTTTCGGGTAACG<br>GGCAGAGGCGTTAATGCCTTCGCGCTGTGCGTCCGCATCCTGAATGCAGCCATAGGTG<br>CGCAGACCGCGTGCCTGAGTGTTCCCCAGCACCATCGTGTTGTCCGGCAGGAAGTTCT<br>TTTTGACGCCGTTTTCCACGTACTGTCCGGAATACACGACGATGGCCACATCGCCATAC<br>ATCCCCTTATAGGACACCGCTTTGCCCAGGTCTTTCACCGCTGTCTCCAGCTCGGAATTA<br>GAGCCACGACGGGTATCCAGCTTCTCCTTGACGGCTTTGAAGGAACGGAACAGCGCCC<br>AGCCTTTCGGATCGAACACGATGATATTCACCACACCGCTGGCGTTCAGCGCGTAGGCT<br>TCGATATCGTCGGTCGGGTCATACGTGGACTTGTCACGCTTGCTCCACTCCGTGCCGCC<br>GGACTGCGTGATGTTATTCTCCTCACTGCGGCCCATATCCACCTCAACCGGATCGAAGG<br>CTTCACCGGTCATGGTGTATTTGCCCTTAAGCACGGCAGAAACTGCCTGCATCTCTTCG<br>ACCTGAGCAATGGCCAGCTCTTCGTCACGCATGTTCTGCATGATGATGCGACGGCGGC<br>GGTAAGCCGGGTCCGCCAGATTCTGCGGATCTTCATCCGGCAGGCGACGCAGGGTCAT<br>CTGCGGATTCACTTCATGCTTCGGCTTGACATATCCCGGCGTAAATTCAGAGGTGGAGC<br>CGCCACGGGAACGGATAACCTCACCGGAAACAATCGGCGAAACGTACAGCGCCATGTT<br>TACCAGTCCCGGAATTTGTGAGAGATAGACTTTCTCCGTGGTGAAGGGATAGCTCTCAC<br>GGAAAAAGAGACGCAGAAACAGCGGATCAAACTTAAATTTCTGCTCATTTGCCGCCAGC<br>AGTTGGGCGGTTGTGTACATCGACATAAAAAAATCCCGTAAAAAAAGCCGCACAGGCGG<br>CCTTTAGTGATGAAGGGTAAAGTTAAACGATGCTGATTGCCGTTCCGGCAAACGCGGTC<br>CGTTTTTTCGTCTCGTCGCTGGCAGCCTCCGGCCAGAGCACATCCTCATAACGGAACGT<br>GCCGGACTTGTAGAACGTCAGCGTGGTGCTGGTCTGGTCAGCAGCAACCGCAAGAATG<br>CCAACGGCAGCACCGTCGGTGGTGCCATCCCACGCAACCAGCTTACGGCTGGAGGTGT<br>CCAGCATCAGCGGGGTCATTGCAGGCGCTTTCGCACTCAATCCGCCGGGCGCGGTTGC<br>GGTATGAGCCGGGTCACTGTTGCCCTGCGGCTGGTAATGGGTAAAGGTTTCTTTGCTCG<br>TCATAAACATCCCTTACACTGGTGTGTTCAGCAAATCGTTAACGGCATCAGATGCCGGGT<br>TACCTGCAGCCAGCGGTGCCGGTGCCCCCTGCATCAGACGATCCAGCGCAGTGTCACT<br>GCGCGCCTGTGCACTCTGTGGTGCTGCGGCCAGAATGCGGCGGGCCGTTTTCACGGTC<br>ATACCGGGGGTTTCTGCCAGCACGCGTGCCTGTTCTTCGCGTCCGTGAGCCTCCTCACA<br>GTTGAGAATTCGTCGTAGAATTCAACGTGGAATTCCTATCGGAATTCTCGGATGAATTCG<br>GATCCTTATGATTCTCGTTCAGGGTACCCTGATCCTGAAAAAATCGTATGAAGTGATGCA<br>TGGGGAATTGGGGAATGTGGATTCGATCCAGAGCTGGTCGAATGCGTAATGTTGACGTG<br>CGAGAAATGTTTAGCTAAACCTTTCGGTATATCGTTCGACGCCTACTGCGTAGCCCTGGT<br>CCATAGCGAGTTAACAAACTTACCGGCCGTTGTCCACTCAATGGTATCTCTTGAACAAGA<br>AGAGTCCCTGCGCTTCTCAGTAAGGGTGGTAACCGTTTGTTAAACACCCAACAACTCGA<br>AGTAGTGTGCGCTTTGTCTAAGTGCAGGTGTCCTCAGTATGTTTGTCCATACTGTTGTCT<br>AGTCTACGTTCGGCTACATCTACCTATTTGAGGGCCAGCGGTACACAGAGGTGTATTAT |

| | |
|---|---|
| | CCAACGGACGTAACGTAAGGGCTTCGATGTCGAGAGTAGATAACACACTATGAAGAGCT<br>GCTACCATGTCCCCTGGCGCATAAGGGTCCCCGCATGGCTTCTCCAGCGAAGTGGTCA<br>TTAGACCTTCCCGTGCTGGATCAGATCACCTAGCTGTGCACGTAGTTCGGTTGGCCTTG<br>GCCATAACTGTTATCGCGATCAGGAATTAGCTTACGTGCTACCACAGCACAATAGTGACA<br>TCGTAGAAGCTCAGTTAGTGCCGGTTTTTCGGAAGGAGGACTTCTATACCTCCACAAGA<br>ATCGCTCCGTTGGAACTATTTAGTTGCTAGTTACACTTTTACTCACAACCCGACGAACTC<br>TTAACAAAGAGTGCTCGGGATAACTGCGATAGTCGATGCACCGCCGGCCGCGACTTCAT<br>TGATCAGATGCCTTAAATAACGAATAAACGATAACATGAGTGCGTCGAACAATAACAGGC<br>AACATAATGCGGTTCCGAGCCGGCTCGCTTCCGCGGGTCCGGAAGTGTGCCCTGGGTT<br>GAACTATCCGTGCTCCCCAGTGGTAACCGTCACGAGAGGCCTGGTCCTGAGGATCATA<br>CTTTCTCGATTATTCTTGCACTCGTTCTCGAAGACTAATCCGGCGGAAGATTTAAATTGTC<br>ACCTTTTGCAGACCGCAAGAGTTGTAGGATAACGTCCTGTATCACCCCAAGCGCCTTCT<br>AAGATTGATGGCGCATGCGGTTGGGGACTCTATTTTAGCGGCTAAAAGTGCCGTATTTG<br>CAGTACCAGCGTACGGCCCACAGAATGATGTCACGCTGAAAATGCCGGCCTTTGAATGG<br>GTTCATGTGCAGCTCCATAAGCAAAAGGGGATGATAAGTTTATCACCACCGACTATTTGC<br>AACAGTGCCGTTGATCGTGCTATGATCGACTCTGGTACTGCAAATACGGCACTTTTAGCC<br>GCTAAAATAGAGTCCCCAACCGCGACGACTGGTACTGCAAATACGGCACTTTTAGCCGC<br>TAAAATAGAGTCCCCAACCGCGACGACTGGTACTGCAAATACGGCACTTTTAGCCGCTA<br>AAATAGAGTCCCCAACCGCGACGAGCGTCCCAAGCTCTCATACCCTTTTAGCCGCTAAA<br>ATCCTGACCAATACGTAGTCAGGCGTCCCAAGCTCTCATACCCTTTTAGCCGCTAAAATC<br>CTGACCAATACGTAGTCAGGCGTCCCAAGCTCTCATACCCTTTTAGCCGCTAAAATCCTG<br>ACCAATACGTAGTCAGGCGTCCCAAGCTCTCATACCCTTTTAGCCGCTAAAATCCTGACC<br>AATACGTAGTCAGTCGTCATGCCCTCGTGGTCAGGTCTGGACGACGAGCCGTTCGATCC<br>TGCCACGTCGCCCGTTACACCGGACCTTGGAGTTGTCTCTGACACATTCTGGCGCCTGC<br>CAAATGTAAAGCGCAGCGCCCATCCATTTGCCTTTGCGGCAGCGGGGCCACAGGCAGA<br>GCAGATCATCTCTGATCCATTGCCCCTGCCACCTCACTCGCCTGCAAGCCCGGTCGCCC<br>GTGTCCATGAACTCGATGGGCAGGTACTTCTCCTCGGCGTGGGACACGATGCCAACAC<br>GACGCTGCATCTTGCCGAGTTGATGGCAAAGGTTCCCTATGGGGTGCCGAGACACTGC<br>ACCATTCTTCAGGATGGCAAGTTGGTACGCGTCGATTATCTAGTCGCGAAGACTAATCC<br>GGCGGAAGATTTAAATTGTCACCTTTTGCAGACCGCAAGAGTTGTAGGATAACGTCCTG<br>TATCACCCCAAGCGCCTTCTAAGATTGATGGCGCATGCGGTTGGGGACTCTATTTTAGC<br>GGCTAAAAGTGCCGTATTTGCAGTACCAGCGTACGGCCCACAGAATGATGTCACGCTGA<br>AAATGCCGGCCTTTGAATGGGTTCATGTGCAGCTCCATAAGCAAAAGGGGATGATAAGT<br>TTATCACCACCGACTATTTGCAACAGTGCCGTTGATCGTGCTATGATCGACTCTGGTACT<br>GCAAATACGGCACTTTTAGCCGCTAAAATAGAGTCCCCAACCGCGACGACTGGTACTGC<br>AAATACGGCACTTTTAGCCGCTAAAATAGAGTCCCCAACCGCGACGACTGGTACTGCAA<br>ATACGGCACTTTTAGCCGCTAAAATAGAGTCCCCAACCGCGACGAGCGTCCCAAGCTCT<br>CATACCCTTTTAGCCGCTAAAATCCTGACCAATACGTAGTCAGGCGTCCCAAGCTCTCAT<br>ACCCTTTTAGCCGCTAAAATCCTGACCAATACGTAGTCAGGCGTCCCAAGCTCTCATACC<br>CTTTTAGCCGCTAAAATCCTGACCAATACGTAGTCAGGCGTCCCAAGCTCTCATACCCTT<br>TTAGCCGCTAAAATCCTGACCAATACGTAGTCAGTCGTCATGCCCTCGTGGTCAGGTCT<br>GGACGACGAGCCGTTCGATCCTGCCACGTCGCCCGTTACACCGGACCTTGGAGTTGTC<br>TCTGACACATTCTGGCGCTCGAGAATGACCACTGCTGTGAGCGCTTTGCCTTGGCGGAC<br>AGGTGGCTCAAGGAGAAGAGCCTTCAGAAGGAAGGTCCAGTCGGTCATGCCTTTGCTC<br>GGTTGATCCGCTCCCGCGACATTGTGGCGACAGCCCTGGGTCAACTGGGCCGAGATCC<br>GTTGATCTTCCTGCATCCGCCAGAGGCGGGATGCGAAGAATGCGATGCCGCTCGCCAG<br>TCGATTGGCTGAGCTCATAAGTGAGGAAGGCCGGGCGGGAAACTGCCCGGCCTGAACA<br>TACCTGAATGGTTATCCCCGCTGACGCGGGGAACATAAGTTCGGGGATCCTCTAGAGTC<br>GACCTGCAGGCATGCAAGCTTGGCGTAATCATGGTCATAGCTGTTTCCTGTGTGAAATT<br>GTTATCCGCTCACAATTCCACACAACATACGAGCCGGAAGCATAAAGTGTAAAGCCTGG<br>GGTGCCTAATGAGTGAGCTAACTCACATTAATTGCGTTGCGCTCACTGCCCGCTTTCCA<br>GTCGGGAAACCTGTCGTGCCAGCTGCATTAATGAATCGGCCAACGCGCGGGGAGAGGC<br>GGTTTGCGTATTGGGCGCTCTTCCGCTTCCTCGCTCACTGACTCGCTGCGCTCGGTCGT<br>TCGGCTGCGGCGAGCGGTATCAGCTCACTCAAAGGCGGTAATACGGTTATCCACAGAAT<br>CAGGGGATAACGCAGGAAAGAACATGTGAGCAAAAGGCCAGCAAAAGGCCAGGAACCG<br>TAAAAAGGCCGCGTTGCTGGCGTTTTTCCATAGGCTCCGCCCCCCTGACGAGCATCACA<br>AAAATCGACGCTCAAGTCAGAGGTGGCGAAACCCGACAGGACTATAAAGATACCAGGC<br>GTTTCCCCCTGGAAGCTCCCTCGTGCGCTCTCCTGTTCCGACCCTGCCGCTTACCGGAT<br>ACCTGTCCGCCTTTCTCCCTTCGGGAAGCGTGGCGCTTTCTCATAGCTCACGCTGTAGG<br>TATCTCAGTTCGGTGTAGGTCGTTCGCTCCAAGCTGGGCTGTGTGCACGAACCCCCCGT |

| | |
|---|---|
| | TCAGCCCGACCGCTGCGCCTTATCCGGTAACTATCGTCTTGAGTCCAACCCGGTAAGAC<br>ACGACTTATCGCCACTGGCAGCAGCCACTGGTAACAGGATTAGCAGAGCGAGGTATGTA<br>GGCGGTGCTACAGAGTTCTTGAAGTGGTGGCCTAACTACGGCTACACTAGAAGGACAGT<br>ATTTGGTATCTGCGCTCTGCTGAAGCCAGTTACCTTCGGAAAAAGAGTTGGTAGCTCTTG<br>ATCCGGCAAACAAACCACCGCTGGTAGCGGTGGTTTTTTTGTTTGCAAGCAGCAGATTA<br>CGCGCAGAAAAAAAGGATCTCAAGAAGATCCTTTGATCTTTTCTACGGGGTCTGACGCT<br>CAGTGGAACGAAAACTCACGTTAAGGGATTTTGGTCATGAGATTATCAAAAAGGATCTTC<br>ACCTAGATCCTTTTAAATTAAAAATGAAGTTTTAAATCAATCTAAAGTATATATGAGTAAAC<br>TTGGTCTGACAGTTACCAATGCTTAATCAGTGAGGCACCTATCTCAGCGATCTGTCTATT<br>TCGTTCATCCATAGTTGCCTGACTCCCCGTCGTGTAGATAACTACGATACGGGAGGGCT<br>TACCATCTGGCCCCAGTGCTGCAATGATACCGCGAGACCCACGCTCACCGGCTCCAGA<br>TTTATCAGCAATAAACCAGCCAGCCGGAAGGGCCGAGCGCAGAAGTGGTCCTGCAACT<br>TTATCCGCCTCCATCCAGTCTATTAATTGTTGCCGGGAAGCTAGAGTAAGTAGTTCGCCA<br>GTTAATAGTTTGCGCAACGTTGTTGCCATTGCTACAGGCATCGTGGTGTCACGCTCGTC<br>GTTTGGTATGGCTTCATTCAGCTCCGGTTCCCAACGATCAAGGCGAGTTACATGATCCC<br>CCATGTTGTGCAAAAAAGCGGTTAGCTCCTTCGGTCCTCCGATCGTTGTCAGAAGTAAG<br>TTGGCCGCAGTGTTATCACTCATGGTTATGGCAGCACTGCATAATTCTCTTACTGTCATG<br>CCATCCGTAAGATGCTTTTCTGTGACTGGTGAGTACTCAACCAAGTCATTCTGAGAATAG<br>TGTATGCGGCGACCGAGTTGCTCTTGCCCGGCGTCAATACGGGATAATACCGCGCCAC<br>ATAGCAGAACTTTAAAAGTGCTCATCATTGGAAAACGTTCTTCGGGGCGAAAACTCTCAA<br>GGATCTTACCGCTGTTGAGATCCAGTTCGATGTAACCCACTCGTGCACCCAACTGATCTT<br>CAGCATCTTTTACTTTCACCAGCGTTTCTGGGTGAGCAAAAACAGGAAGGCAAAATGCC<br>GCAAAAAAGGGAATAAGGGCGACACGGAAATGTTGAATACTCATACTCTTCCTTTTTCAA<br>TATTATTGAAGCATTTATCAGGGTTATTGTCTCATGAGCGGATACATATTTGAATGTATTT<br>AGAAAAATAAACAAATAGGGGTTCCGCGCACATTTCCCCGAAAAGTGCCACCTGACGTC<br>TAAGAAACCATTATTATCATGACATTAACCTATAAAAATAGGCGTATCACGAGGCCCTTTC<br>GTCTCGCGCGTTTCGGTGATGACGGTGAAAACCTCTGACACATGCAGCTCCCGGAGAC<br>GGTCACAGCTTGTCTGTAAGCGGATGCCGGGAGCAGACAAGCCCGTCAGGGCGCGTCA<br>GCGGGTGTTGGCGGGTGTCGGGGCTGGCTTAACTATGCGGCATCAGAGCAGATTGTAC<br>TGAGAGTGCACCATATGGGGCC |
| C-trap central fragment<br>(22394 bp) | GGCCGCGGTGTGCTCCTTATTTATACATAACGAAAAACGCCTCGAGTGAAGCGTTATTG<br>GTATGCGGTAAAACCGCACTCAGGCGGCCTTGATAGTCATATCATCTGAATCAAATATTC<br>CTGATGTATCGATATCGGTAATTCTTATTCCTTCGCTACCATCCATTGGAGGCCATCCTT<br>CCTGACCATTTCCATCATTCCAGTCGAACTCACACACAACACCATATGCATTTAAGTCGC<br>TTGAAATTGCTATAAGCAGAGCATGTTGCGCCAGCATGATTAATACAGCATTTAATACAG<br>AGCCGTGTTTATTGAGTCGGTATTCAGAGTCTGACCAGAAATTATTAATCTGGTGAAGTT<br>TTTCCTCTGTCATTACGTCATGGTCGATTTCAATTTCTATTGATGCTTTCCAGTCGTAATC<br>AATGATGTATTTTTTGATGTTTGACATCTGTTCATATCCTCACAGATAAAAAATCGCCCTC<br>ACACTGGAGGGCAAAGAAGATTTCCAATAATCAGAACAAGTCGGCTCCTGTTTAGTTAC<br>GAGCGACATTGCTCCGTGTATTCACTCGTTGGAATGAATACACAGTGCAGTGTTTATTCT<br>GTTATTTATGCCAAAAATAAAGGCCACTATCAGGCAGCTTTGTTGTTCTGTTTACCAAGTT<br>CTCTGGCAATCATTGCCGTCGTTCGTATTGCCCATTTATCGACATATTTCCCATCTTCCAT<br>TACAGGAAACATTTCTTCAGGCTTAACCATGCATTCCGATTGCAGCTTGCATCCATTGCA<br>TCGCTTGAATTGTCCACACCATTGATTTTTATCAATAGTCGTAGTCATACGGATAGTCCTG<br>GTATTGTTCCATCACATCCTGAGGATGCTCTTCGAACTCTTCAAATTCTTCTTCCATATAT<br>CACCTTAAATAGTGGATTGCGGTAGTAAAGATTGTGCCTGTCTTTTAACCACATCAGGCT<br>CGGTGGTTCTCGTGTACCCCTACAGCGAGAAATCGGATAAACTATTACAACCCCTACAG<br>TTTGATGAGTATAGAAATGGATCCACTCGTTATTCTCGGACGAGTGTTCAGTAATGAACC<br>TCTGGAGAGAACCATGTATATGATCGTTATCTGGGTTGGACTTCTGCTTTTAAGCCCAGA<br>TAACTGGCCTGAATATGTTAATGAGAGAATCGGTATTCCTCATGTGTGGCATGTTTTCGT<br>CTTTGCTCTTGCATTTTCGCTAGCAATTAATGTGCATCGATTATCAGCTATTGCCAGCGC<br>CAGATATAAGCGATTTAAGCTAAGAAAACGCATTAAGATGCAAAACGATAAAGTGCGATC<br>AGTAATTCAAAACCTTACAGAAGAGCAATCTATGGTTTTGTGCGCAGCCCTTAATGAAGG<br>CAGGAAGTATGTGGTTACATCAAAACAATTCCCATACATTAGTGAGTTGATTGAGCTTGG<br>TGTGTTGAACAAAACTTTTTCCCGATGGAATGGAAAGCATATATTATTCCCTATTGAGGAT<br>ATTTACTGGACTGAATTAGTTGCCAGCTATGATCCATATAATATTGAGATAAAGCCAAGG<br>CCAATATCTAAGTAACTAGATAAGAGGAATCGATTTTCCCTTAATTTTCTGGCGTCCACTG<br>CATGTTATGCCGCGTTCGCCAGGCTTGCTGTACCATGTGCGCTGATTCTTGCGCTCAAT<br>ACGTTGCAGGTTGCTTTCAATCTGTTTGTGGTATTCAGCCAGCACTGTAAGGTCTATCGG<br>ATTTAGTGCGCTTTCTACTCGTGATTTCGGTTTGCGATTCAGCGAGAGAATAGGGCGGTT |

| | |
|---|---|
| | AACTGGTTTTGCGCTTACCCCAACCAACAGGGGATTTGCTGCTTTCCATTGAGCCTGTTT<br>CTCTGCGCGACGTTCGCGGCGGCGTGTTTGTGCATCCATCTGGATTCTCCTGTCAGTTA<br>GCTTTGGTGGTGTGTGGCAGTTGTAGTCCTGAACGAAAACCCCCCGCGATTGGCACATT<br>GGCAGCTAATCCGGAATCGCACTTACGGCCAATGCTTCGTTTCGTATCACACACCCCAA<br>AGCCTTCTGCTTTGAATGCTGCCCTTCTTCAGGGCTTAATTTTTAAGAGCGTCACCTTCA<br>TGGTGGTCAGTGCGTCCTGCTGATGTGCTCAGTATCACCGCCAGTGGTATTTATGTCAA<br>CACCGCCAGAGATAATTTATCACCGCAGATGGTTATCTGTATGTTTTTTATATGAATTTAT<br>TTTTTGCAGGGGGGCATTGTTTGGTAGGTGAGAGATCTGAATTGCTATGTTTAGTGAGTT<br>GTATCTATTTATTTTTCAATAAATACAATTGGTTATGTGTTTTGGGGGCGATCGTGAGGCA<br>AAGAAAACCCGGCGCTGAGGCCGGGTTATTCTTGTTCTCTGGTCAAATTATATAGTTGGA<br>AAACAAGGATGCATATATGAATGAACGATGCAGAGGCAATGCCGATGGCGATAGTGGGT<br>ATCATGTAGCCGCTTATGCTGGAAAGAAGCAATAACCCGCAGAAAAACAAAGCTCCAAG<br>CTCAACAAAACTAAGGGCATAGACAATAACTACCGATGTCATATACCCATACTCTCTAAT<br>CTTGGCCAGTCGGCGCGTTCTGCTTCCGATTAGAAACGTCAAGGCAGCAATCAGGATTG<br>CAATCATGGTTCCTGCATATGATGACAATGTCGCCCCAAGACCATCTCTATGAGCTGAAA<br>AAGAAACACCAGGAATGTAGTGGCGGAAAAGGAGATAGCAAATGCTTACGATAACGTAA<br>GGAATTATTACTATGTAAACACCAGGCATGATTCTGTTCCGCATAATTACTCCTGATAATT<br>AATCCTTAACTTTGCCCACCTGCCTTTTAAAACATTCCAGTATATCACTTTTCATTCTTGC<br>GTAGCAATATGCCATCTCTTCAGCTATCTCAGCATTGGTGACCTTGTTCAGAGGCGCTGA<br>GAGATGGCCTTTTTCTGATAGATAATGTTCTGTTAAAATATCTCCGGCCTCATCTTTTGCC<br>CGCAGGCTAATGTCTGAAAATTGAGGTGACGGGTTAAAAATAATATCCTTGGCAACCTTT<br>TTTATATCCCTTTTAAATTTTGGCTTAATGACTATATCCAATGAGTCAAAAAGCTCCCCTTC<br>AATATCTGTTGCCCCTAAGACCTTTAATATATCGCCAAATACAGGTAGCTTGGCTTCTAC<br>CTTCACCGTTGTTCGGCCGATGAAATGCATATGCATAACATCGTCTTTGGTGGTTCCCCT<br>CATCAGTGGCTCTATCTGAACGCGCTCTCCACTGCTTAATGACATTCCTTTCCCGATTAA<br>AAAATCTGTCAGATCGGATGTGGTCGGCCCGAAAACAGTTCTGGCAAAACCAATGGTGT<br>CGCCTTCAACAAACAAAAAAGATGGGAATCCCAATGATTCGTCATCTGCGAGGCTGTTCT<br>TAATATCTTCAACTGAAGCTTTAGAGCGATTTATCTTCTGAACCAGACTCTTGTCATTTGT<br>TTTGGTAAAGAGAAAAGTTTTTCCATCGATTTTATGAATATACAAATAATTGGAGCCAACC<br>TGCAGGTGATGATTATCAGCCAGCAGAGAATTAAGGAAAACAGACAGGTTTATTGAGCG<br>CTTATCTTTCCCTTTATTTTTGCTGCGGTAAGTCGCATAAAAACCATTCTTCATAATTCAAT<br>CCATTTACTATGTTATGTTCTGAGGGGAGTGAAAATTCCCCTAATTCGATGAAGATTCTTG<br>CTCAATTGTTATCAGCTATGCGCCGACCAGAACACCTTGCCGATCAGCCAAACGTCTCTT<br>CAGGCCACTGACTAGCGATAACTTTCCCCACAACGGAACAACTCTCATTGCATGGGATC<br>ATTGGGTACTGTGGGTTTAGTGGTTGTAAAAACACCTGACCGCTATCCCTGATCAGTTTC<br>TTGAAGGTAAACTCATCACCCCCAAGTCTGGCTATGCAGAAATCACCTGGCTCAACAGC<br>CTGCTCAGGGTCAACGAGAATTAACATTCCGTCAGGAAAGCTTGGCTTGGAGCCTGTTG<br>GTGCGGTCATGGAATTACCTTCAACCTCAAGCCAGAATGCAGAATCACTGGCTTTTTTGG<br>TTGTGCTTACCCATCTCTCCGCATCACCTTTGGTAAAGGTTCTAAGCTTAGGTGAGAACA<br>TCCCTGCCTGAACATGAGAAAAAACAGGGTACTCATACTCACTTCTAAGTGACGGCTGC<br>ATACTAACCGCTTCATACATCTCGTAGATTTCTCTGGCGATTGAAGGGCTAAATTCTTCAA<br>CGCTAACTTTGAGAATTTTTGTAAGCAATGCGGCGTTATAAGCATTTAATGCATTGATGC<br>CATTAAATAAAGCACCAACGCCTGACTGCCCCATCCCCATCTTGTCTGCGACAGATTCCT<br>GGGATAAGCCAAGTTCATTTTTCTTTTTTTCATAAATTGCTTTAAGGCGACGTGCGTCCTC<br>AAGCTGCTCTTGTGTTAATGGTTTCTTTTTTGTGCTCATACGTTAAATCTATCACCGCAAG<br>GGATAAATATCTAACACCGTGCGTGTTGACTATTTTACCTCTGGCGGTGATAATGGTTGC<br>ATGTACTAAGGAGGTTGTATGGAACAACGCATAACCCTGAAAGATTATGCAATGCGCTTT<br>GGGCAAACCAAGACAGCTAAAGATCTCGGCGTATATCAAAGCGCGATCAACAAGGCCAT<br>TCATGCAGGCCGAAAGATTTTTTTAACTATAAACGCTGATGGAAGCGTTTATGCGGAAGA<br>GGTAAAGCCCTTCCCGAGTAACAAAAAAACAACAGCATAAATAACCCCGCTCTTACACAT<br>TCCAGCCCTGAAAAAGGGCATCAAATTAAACCACACCTATGGTGTATGCATTTATTTGCA<br>TACATTCAATCAATTGTTATCTAAGGAAATACTTACATATGGTTCGTGCAAACAAACGCAA<br>CGAGGCTCTACGAATCGAGAGTGCGTTGCTTAACAAAATCGCAATGCTTGGAACTGAGA<br>AGACAGCGGAAGCTGTGGGCGTTGATAAGTCGCGTCGAGCCTCAGCTCATGTCATCCT<br>CAGCACACTTGACCCTCAGCTCAGCTAGCCTCAGCCTACAATCACCTCAGCGAATTCGG<br>TGACCCTTACGCGAATCCGCTTTCAGACGTTGACTGGTCGCGTCTGGCAAAAGTTAAAG<br>ACCTGACGCCCGGCGAACTGACCGCTGAGTCCTATGACGACAGCTATCTCGATGATGAA<br>GATGCAGACTGGACTGCGACCGGGCAGGGGCAGAAATCTGCCGGAGATACCAGCTTCA<br>CGCTGGCGTGGATGCCCGGAGAGCAGGGGCAGCAGGCGCTGCTGGCGTGGTTTAATG<br>AAGGCGATACCCGTGCCTATAAAATCCGCTTCCCGAACGGCACGGTCGATGTGTTCCGT |

| | |
|---|---|
| | GGCTGGGTCAGCAGTATCGGTAAGGCGGTGACGGCGAAGGAAGTGATCACCCGCACG<br>GTGAAAGTCACCAATGTGGGACGTCCGTCGATGGCAGAAGATCGCAGCACGGTAACAG<br>CGGCAACCGGCATGACCGTGACGCCTGCCAGCACCTCGGTGGTGAAAGGGCAGAGCA<br>CCACGCTGACCGTGGCCTTCCAGCCGGAGGGCGTAACCGACAAGAGCTTTCGTGCGGT<br>GTCTGCGGATAAAACAAAAGCCACCGTGTCGGTCAGTGGTATGACCATCACCGTGAACG<br>GCGTTGCTGCAGGCAAGGTCAACATTCCGGTTGTATCCGGTAATGGTGAGTTTGCTGCG<br>GTTGCAGAAATTACCGTCACCGCCAGTTAATCCGGAGAGTCAGCGATGTTCCTGAAAAC<br>CGAATCATTTGAACATAACGGTGTGACCGTCACGCTTTCTGAACTGTCAGCCCTGCAGC<br>GCATTGAGCATCTCGCCCTGATGAAACGGCAGGCAGAACAGGCGGAGTCAGACAGCAA<br>CCGGAAGTTTACTGTGGAAGACGCCATCAGAACCGGCGCGTTTCTGGTGGCGATGTCC<br>CTGTGGCATAACCATCCGCAGAAGACGCAGATGCCGTCCATGAATGAAGCCGTTAAACA<br>GATTGAGCAGGAAGTGCTTACCACCTGGCCCACGGAGGCAATTTCTCATGCTGAAAACG<br>TGGTGTACCGGCTGTCTGGTATGTATGAGTTTGTGGTGAATAATGCCCCTGAACAGACA<br>GAGGACGCCGGGCCCGCAGAGCCTGTTTCTGCGGGAAAGTGTTCGACGGTGAGCTGA<br>GTTTTGCCCTGAAACTGGCGCGTGAGATGGGGCGACCCGACTGGCGTGCCATGCTTGC<br>CGGGATGTCATCCACGGAGTATGCCGACTGGCACCGCTTTTACAGTACCCATTATTTTCA<br>TGATGTTCTGCTGGATATGCACTTTTCCGGGCTGACGTACACCGTGCTCAGCCTGTTTTT<br>CAGCGATCCGGATATGCATCCGCTGGATTTCAGTCTGCTGAACCGGCGCGAGGCTGAC<br>GAAGAGCCTGAAGATGATGTGCTGATGCAGAAAGCGGCAGGGCTTGCCGGAGGTGTCC<br>GCTTTGGCCCGGACGGGAATGAAGTTATCCCCGCTTCCCCGGATGTGGCGGACATGAC<br>GGAGGATGACGTAATGCTGATGACAGTATCAGAAGGGATCGCAGGAGGAGTCCGGTAT<br>GGCTGAACCGGTAGGCGATCTGGTCGTTGATTTGAGTCTGGATGCGGCCAGATTTGAC<br>GAGCAGATGGCCAGAGTCAGGCGTCATTTTTCTGGTACGGAAAGTGATGCGAAAAAAAC<br>AGCGGCAGTCGTTGAACAGTCGCTGAGCCGACAGGCGCTGGCTGCACAGAAAGCGGG<br>GATTTCCGTCGGGCAGTATAAAGCCGCCATGCGTATGCTGCCTGCACAGTTCACCGACG<br>TGGCCACGCAGCTTGCAGGCGGGCAAAGTCCGTGGCTGATCCTGCTGCAACAGGGGG<br>GGCAGGTGAAGGACTCCTTCGGCGGGATGATCCCCATGTTCAGGGGGCTTGCCGGTGC<br>GATCACCCTGCCGATGGTGGGGGCCACCTCGCTGGCGGTGGCGACCGGTGCGCTGGC<br>GTATGCCTGGTATCAGGGCAACTCAACCCTGTCCGATTTCAACAAAACGCTGGTCCTTTC<br>CGGCAATCAGGCGGGACTGACGGCAGATCGTATGCTGGTCCTGTCCAGAGCCGGGCA<br>GGCGGCAGGGCTGACGTTTAACCAGACCAGCGAGTCACTCAGCGCACTGGTTAAGGCG<br>GGGGTAAGCGGTGAGGCTCAGATTGCGTCCATCAGCCAGAGTGTGGCGCGTTTCTCCT<br>CTGCATCCGGCGTGGAGGTGGACAAGGTCGCTGAAGCCTCTAGAGGATCCCCGAACTT<br>ATGTTCCCCGCGTCAGCGGGGATAACCATTCAGGTATGTTCAGGCCGGGCAGTTTCCC<br>GCCCGGCCTTCCTCACTTATGAGCTCAGCCAATCGACTGGCGAGCGGCATCGCATTCTT<br>CGCATCCCGCCTCTGGCGGATGCAGGAAGATCAACGGATCTCGGCCCAGTTGACCCAG<br>GGCTGTCGCCACAATGTCGCGGGAGCGGATCAACCGAGCAAAGGCATGACCGACTGGA<br>CCTTCCTTCTGAAGGCTCTTCTCCTTGAGCCACCTGTCCGCCAAGGCAAAGCGCTCACA<br>GCAGTGGTCATTCTCGAGATAATCGACGCGTACCAACTTGCCATCCTGAAGAATGGTGC<br>AGTGTCTCGGCACCCCATAGGGAACCTTTGCCATCAACTCGGCAAGATGCAGCGTCGT<br>GTTGGCATCGTGTCCCACGCCGAGGAGAAGTACCTGCCCATCGAGTTCATGGACACGG<br>GCGACCGGGCTTGCAGGCGAGTGAGGTGGCAGGGGCAATGGATCAGAGATGATCTGC<br>TCTGCCTGTGGCCCCGCTGCCGCAAAGGCAAATGGATGGGCGCTGCGCTTTACATTTG<br>GCAGGCGCCAGAATGTGTCAGAGACAACTCCAAGGTCCGGTGTAACGGGCGACGTGGC<br>AGGATCGAACGGCTCGTCGTCCAGACCTGACCACGAGGGCATGACGACTGACTACGTA<br>TTGGTCAGGATTTTAGCGGCTAAAAGGGTATGAGAGCTTGGGACGCCTGACTACGTATT<br>GGTCAGGATTTTAGCGGCTAAAAGGGTATGAGAGCTTGGGACGCCTGACTACGTATTGG<br>TCAGGATTTTAGCGGCTAAAAGGGTATGAGAGCTTGGGACGCCTGACTACGTATTGGTC<br>AGGATTTTAGCGGCTAAAAGGGTATGAGAGCTTGGGACGCTCGTCGCGGTTGGGGACT<br>CTATTTTAGCGGCTAAAAGTGCCGTATTTGCAGTACCAGTCGTCGCGGTTGGGGACTCT<br>ATTTTAGCGGCTAAAAGTGCCGTATTTGCAGTACCAGTCGTCGCGGTTGGGGACTCTAT<br>TTTAGCGGCTAAAAGTGCCGTATTTGCAGTACCAGAGTCGATCATAGCACGATCAACGG<br>CACTGTTGCAAATAGTCGGTGGTGATAAACTTATCATCCCCTTTTGCTTATGGAGCTGCA<br>CATGAACCCATTCAAAGGCCGGCATTTTCAGCGTGACATCATTCTGTGGGCCGTACGCT<br>GGTACTGCAAATACGGCACTTTTAGCCGCTAAAATAGAGTCCCCAACCGCATGCGCCAT<br>CAATCTTAGAAGGCGCTTGGGGTGATACAGGACGTTATCCTACAACTCTTGCGGTCTGC<br>AAAAGGTGACAATTTAAATCTTCCGCCGGATTAGTCTTCGAGACTAGAGAATGCTACGTA<br>CCTGATGAGCTCCAGCTTTTGTTCCCTTTAGTGAGGGTTAATTGCGCGCTTGGCGTAATC<br>ATGGTCATAGCTGTTTCCTGTGTGAAATTGTTATCCGCTCACAATTCCACACAACATACG<br>AGCCGGAAGCATAAAGTGTAAAGCCTGGGGTGCCTAATGAGTGAGCTAACTCACATTAA |

| | |
|---|---|
| | TTGCGTTGCGCTCACTGCCCGCTTTCCAGTCGGGAAACCTGTCGTGCCAGCTGCATTAA<br>TGAATCGGCCAACGCGCGGGGAGAGGCGGTTTGCGTATTGGGCGCTCTTCCGCTTCCT<br>CGCTCACTGACTCGCTGCGCTCGGTCGTTCGGCTGCGGCGAGCGGTATCAGCTCACTC<br>AAAGGCGGTAATACGGTTATCCACAGAATCAGGGGATAACGCAGGAAAGAACATGTGAG<br>CAAAAGGCCAGCAAAAGGCCAGGAACCGTAAAAAGGCCGCGTTGCTGGCGTTTTTCCAT<br>AGGCTCCGCCCCCCTGACGAGCATCACAAAAATCGACGCTCAAGTCAGAGGTGGCGAA<br>ACCCGACAGGACTATAAAGATACCAGGCGTTTCCCCCTGGAAGCTCCCTCGTGCGCTCT<br>CCTGTTCCGACCCTGCCGCTTACCGGATACCTGTCCGCCTTTCTCCCTTCGGGAAGCGT<br>GGCGCTTTCTCATAGCTCACGCTGTAGGTATCTCAGTTCGGTGTAGGTCGTTCGCTCCA<br>AGCTGGGCTGTGTGCACGAACCCCCCGTTCAGCCCGACCGCTGCGCCTTATCCGGTAA<br>CTATCGTCTTGAGTCCAACCCGGTAAGACACGACTTATCGCCACTGGCAGCAGCCACTG<br>GTAACAGGATTAGCAGAGCGAGGTATGTAGGCGGTGCTACAGAGTTCTTGAAGTGGTG<br>GCCTAACTACGGCTACACTAGAAGGACAGTATTTGGTATCTGCGCTCTGCTGAAGCCAG<br>TTACCTTCGGAAAAAGAGTTGGTAGCTCTTGATCCGGCAAACAAACCACCGCTGGTAGC<br>GGTGGTTTTTTTGTTTGCAAGCAGCAGATTACGCGCAGAAAAAAAGGATCTCAAGAAGAT<br>CCTTTGATCTTTTCTACGGGGTCTGACGCTCAGTGGAACGAAAACTCACGTTAAGGGATT<br>TTGGTCATGAGATTATCAAAAAGGATCTTCACCTAGATCCTTTTAAATTAAAAATGAAGTT<br>TTAAATCAATCTAAAGTATATATGAGTAAACTTGGTCTGACAGTTACCAATGCTTAATCAG<br>TGAGGCACCTATCTCAGCGATCTGTCTATTTCGTTCATCCATAGTTGCCTGACTCCCCGT<br>CGTGTAGATAACTACGATACGGGAGGGCTTACCATCTGGCCCCAGTGCTGCAATGATAC<br>CGCGAGACCCACGCTCACCGGCTCCAGATTTATCAGCAATAAACCAGCCAGCCGGAAG<br>GGCCGAGCGCAGAAGTGGTCCTGCAACTTTATCCGCCTCCATCCAGTCTATTAATTGTT<br>GCCGGGAAGCTAGAGTAAGTAGTTCGCCAGTTAATAGTTTGCGCAACGTTGTTGCCATT<br>GCTACAGGCATCGTGGTGTCACGCTCGTCGTTTGGTATGGCTTCATTCAGCTCCGGTTC<br>CCAACGATCAAGGCGAGTTACATGATCCCCCATGTTGTGCAAAAAAGCGGTTAGCTCCT<br>TCGGTCCTCCGATCGTTGTCAGAAGTAAGTTGGCCGCAGTGTTATCACTCATGGTTATG<br>GCAGCACTGCATAATTCTCTTACTGTCATGCCATCCGTAAGATGCTTTTCTGTGACTGGT<br>GAGTACTCAACCAAGTCATTCTGAGAATAGTGTATGCGGCGACCGAGTTGCTCTTGCCC<br>GGCGTCAATACGGGATAATACCGCGCCACATAGCAGAACTTTAAAAGTGCTCATCATTG<br>GAAAACGTTCTTCGGGGCGAAAACTCTCAAGGATCTTACCGCTGTTGAGATCCAGTTCG<br>ATGTAACCCACTCGTGCACCCAACTGATCTTCAGCATCTTTTACTTTCACCAGCGTTTCT<br>GGGTGAGCAAAAACAGGAAGGCAAAATGCCGCAAAAAAGGGAATAAGGGCGACACGGA<br>AATGTTGAATACTCATACTCTTCCTTTTTCAATATTATTGAAGCATTTATCAGGGTTATTGT<br>CTCATGAGCGGATACATATTTGAATGTATTTAGAAAAATAAACAAATAGGGGTTCCGCGC<br>ACATTTCCCCGAAAAGTGCCACCTAAATTGTAAGCGTTAATATTTTGTTAAAATTCGCGTT<br>AAATTTTTGTTAAATCAGCTCATTTTTTAACCAATAGGCCGAAATCGGCAAAATCCCTTAT<br>AAATCAAAAGAATAGACCGAGATAGGGTTGAGTGTTGTTCCAGTTTGGAACAAGAGTCC<br>ACTATTAAAGAACGTGGACTCCAACGTCAAAGGGCGAAAAACCGTCTATCAGGGCGATG<br>GCCCACTACGTGAACCATCACCCTAATCAAGTTTTTTGGGGTCGAGGTGCCGTAAAGCA<br>CTAAATCGGAACCCTAAAGGGAGCCCCCGATTTAGAGCTTGACGGGGAAAGCCGGCGA<br>ACGTGGCGAGAAAGGAAGGGAAGAAAGCGAAAGGAGCGGGCGCTAGGGCGCTGGCAA<br>GTGTAGCGGTCACGCTGCGCGTAACCACCACACCCGCCGCGCTTAATGCGCCGCTACA<br>GGGCGCGTCCCATTCGCCATTCAGGCTGCGCAACTGTTGGGAAGGGCGATCGGTGCG<br>GGCCTCTTCGCTATTACGCCAGCTGGCGAAAGGGGGATGTGCTGCAAGGCGATTAAGT<br>TGGGTAACGCCAGGGTTTTCCCAGTCACGACGTTGTAAAACGACGGCCAGTGAGCGCG<br>CGTAATACGACTCACTATAGGGCGAATTGGGTACCCGCTGGAGAAGCCATGCGGGGAC<br>CCTTATGCGCCAGGGGACATGGTAGCAGCTCTTCATAGTGTGTTATCTACTCTCGACATC<br>GAAGCCCTTACGTTACGTCCGTTGGATAATACACCTCTGTGTACCGCTGGCCCTCAAATA<br>GGTAGATGTAGCCGAACGTAGACTAGACAACAGTATGGACAAACATACTGAGGACACCT<br>GCACTTAGACAAAGCGCACACTACTTCGAGTTGTTGGGTGTTTAACAAACGGTTACCAC<br>CCTTACTGAGAAGCGCAGGGACTCTTCTTGTTCAAGAGATACCATTGAGTGGACAACGG<br>CCGGTAAGTTTGTTAACTCGCTATGGACCAGGGCTACGCAGTAGGCGTCGAACGATATA<br>CCGAAAGGTTTAGCTAAACATTTCTCGCACGTCAACATTACGCATTCGACCAGCTCTGGA<br>TCGAATCCACATTCCCCAATTCCCCATGCATCACTTCATACGATTTTTTCAGGATCAGGG<br>TACCAGTTCAGGAAGCGGTGATGCTGATAGAAGCCGGACTGAGTACCTACGAGAAAGA<br>GTGCGCAAAACGCGGTGACGACTATCAGGAAATTTTTGCCCAGCAGGTCCGTGAAACGA<br>TGGAGCGCCGTGCAGCCGGTCTTAAACCGCCCGCCTGGGCGGCTGCAGCATTTGAATC<br>CGGGCTGCGACAATCAACAGAGGAGGAGAAGAGTGACAGCAGAGCTGCGTAATCTCCC<br>GCATATTGCCAGCATGGCCTTTAATGAGCCGCTGATGCTTGAACCCGCCTATGCGCGGG<br>TTTTCTTTTGTGCGCTTGCAGGCCAGCTTGGGATCAGCAGCCTGGCGGATGCGGTGTCC |

| | |
|---|---|
| | GGCGACAGCCTGACTGCCCAGGAGGCACTCGCGACGCTGGCATTATCCGGTGATGATG |
| | ACGGACCACGACAGGCCCGCAGTTATCAGGTCATGAACGGCATCGCCGTGCTGCCGGT |
| | GTCCGGCACGCTGGTCAGCCGGACGCGGGCGCTGCAGCCGTACTCGGGGATGACCGG |
| | TTACAACGGCATTATCGCCCGTCTGCAACAGGCTGCCAGCGATCCGATGGTGGACGGC |
| | ATTCTGCTCGATATGGACACGCCCGGCGGGATGGTGGCGGGGGCATTTGACTGCGCTG |
| | ACATCATCGCCCGTGTGCGTGACATAAAACCGGTATGGGCGCTTGCCAACGACATGAAC |
| | TGCAGTGCAGGTCAGTTGCTTGCCAGTGCCGCCTCCCGGCGTCTGGTCACGCAGACCG |
| | CCCGGACAGGCTCCATCGGCGTCATGATGGCTCACAGTAATTACGGTGCTGCGCTGGA |
| | GAAACAGGGTGTGGAAATCACGCTGATTTACAGCGGCAGCCATAAGGTGGATGGCAAC |
| | CCCTACAGCCATCTTCCGGATGACGTCCGGGAGACACTGCAGTCCCGGATGGACGCAA |
| | CCCGCCAGATGTTTGCGCAGAAGGTGTCGGCATATACCGGCCTGTCCGTGCAGGTTGT |
| | GCTGGATACCGAGGCTGCAGTGTACAGCGGTCAGGAGGCCATTGATGCCGGACTGGCT |
| | GATGAACTTGTTAACAGCACCGATGCGATCACCGTCATGCGTGATGCACTGGATGCACG |
| | TAAATCCCGTCTCTCAGGAGGGCGAATGACCAAAGAGACTCAATCAACAACTGTTTCAG |
| | CCACTGCTTCGCAGGCTGACGTTACTGACGTGGTGCCAGCGACGGAGGGCGAGAACG |
| | CCAGCGCGGCGCAGCCGGACGTGAACGCGCAGATCACCGCAGCGGTTGCGGCAGAAA |
| | ACAGCCGCATTATGGGGATCCTCAACTGTGAGGAGGCTCACGGACGCGAAGAACAGGC |
| | ACGCGTGCTGGCAGAAACCCCCGGTATGACCGTGAAAACGGCCCGCCGCATTCTGGCC |
| | GCAGCACCACAGAGTGCACAGGCGCGCAGTGACACTGCGCTGGATCGTCTGATGCAGG |
| | GGGCACCGGCACCGCTGGCTGCAGGTAACCCGGCATCTGATGCCGTTAACGATTTGCT |
| | GAACACACCAGTGTAAGGGATGTTTATGACGAGCAAAGAAACCTTTACCCATTACCAGC |
| | CGCAGGGCAACAGTGACCCGGCTCATACCGCAACCGCGCCCGGCGGATTGAGTGCGA |
| | AAGCGCCTGCAATGACCCCGCTGATGCTGGACACCTCCAGCCGTAAGCTGGTTGCGTG |
| | GGATGGCACCACCGACGGTGCTGCCGTTGGCATTCTTGCGGTTGCTGCTGACCAGACC |
| | AGCACCACGCTGACGTTCTACAAGTCCGGCACGTTCCGTTATGAGGATGTGCTCTGGCC |
| | GGAGGCTGCCAGCGACGAGACGAAAAAACGGACCGCGTTTGCCGGAACGGCAATCAG |
| | CATCGTTTAACTTTACCCTTCATCACTAAAGGCCGCCTGTGCGGCTTTTTTTACGGGATT |
| | TTTTTATGTCGATGTACACAACCGCCCAACTGCTGGCGGCAAATGAGCAGAAATTTAAGT |
| | TTGATCCGCTGTTTCTGCGTCTCTTTTTCCGTGAGAGCTATCCCTTCACCACGGAGAAAG |
| | TCTATCTCTCACAAATTCCGGGACTGGTAAACATGGCGCTGTACGTTTCGCCGATTGTTT |
| | CCGGTGAGGTTATCCGTTCCCGTGGCGGCTCCACCTCTGAATTTACGCCGGGATATGTC |
| | AAGCCGAAGCATGAAGTGAATCCGCAGATGACCCTGCGTCGCCTGCCGGATGAAGATC |
| | CGCAGAATCTGGCGGACCCGGCTTACCGCCGCCGTCGCATCATCATGCAGAACATGCG |
| | TGACGAAGAGCTGGCCATTGCTCAGGTCGAAGAGATGCAGGCAGTTTCTGCCGTGCTTA |
| | AGGGCAAATACACCATGACCGGTGAAGCCTTCGATCCGGTTGAGGTGGATATGGGCCG |
| | CAGTGAGGAGAATAACATCACGCAGTCCGGCGGCACGGAGTGGAGCAAGCGTGACAAG |
| | TCCACGTATGACCCGACCGACGATATCGAAGCCTACGCGCTGAACGCCAGCGGTGTGG |
| | TGAATATCATCGTGTTCGATCCGAAAGGCTGGGCGCTGTTCCGTTCCTTCAAAGCCGTC |
| | AAGGAGAAGCTGGATACCCGTCGTGGCTCTAATTCCGAGCTGGAGACAGCGGTGAAAG |
| | ACCTGGGCAAAGCGGTGTCCTATAAGGGGATGTATGGCGATGTGGCCATCGTCGTGTAT |
| | TCCGGACAGTACGTGGAAAACGGCGTCAAAAAGAACTTCCTGCCGGACAACACGATGG |
| | TGCTGGGGAACACTCAGGCACGCGGTCTGCGCACCTATGGCTGCATTCAGGATGCGGA |
| | CGCACAGCGCGAAGGCATTAACGCCTCTGCCCGTTACCCGAAAAACTGGGTGACCACC |
| | GGCGATCCGGCGCGTGAGTTCACCATGATTCAGTCAGCACCGCTGATGCTGCTGGCTG |
| | ACCCTGATGAGTTCGTGTCCGTACAACTGGCGTAATCATGGCCCTTCGGGGCCATTGTT |
| | TCTCTGTGGAGGAGTCCATGACGAAAGATGAACTGATTGCCCGTCTCCGCTCGCTGGGT |
| | GAACAACTGAACCGTGATGTCAGCCTGACGGGGACGAAAGAAGAACTGGCGCTCCGTG |
| | TGGCAGAGCTGAAAGAGGAGCTTGATGACACGGATGAAACTGCCGGTCAGGACACCCC |
| | TCTCAGCCGGGAAAATGTGCTGACCGGACATGAAAATGAGGTGGGATCAGCGCAGCCG |
| | GATACCGTGATTCTGGATACGTCTGAACTGGTCACGGTCGTGGCACTGGTGAAGCTGCA |
| | TACTGATGCACTTCACGCCACGCGGGATGAACCTGTGGCATTTGTGCTGCCGGGAACG |
| | GCGTTTCGTGTCTCTGCCGGTGTGGCAGCCGAAATGACAGAGCGCGGCCTGGCCAGAA |
| | TGCAATAACGGGAGGCGCTGTGGCTGATTTCGATAACCTGTTCGATGCTGCCATTGCCC |
| | GCGCCGATGAAACGATACGCGGGTACATGGGAACGTCAGCCACCATTACATCCGGTGA |
| | GCAGTCAGGTGCGGTGATACGTGGTGTTTTTGATGACCCTGAAAATATCAGCTATGCCG |
| | GACAGGGCGTGCGCGTTGAAGGCTCCAGCCCGTCCCTGTTTGTCCGGACTGATGAGGT |
| | GCGGCAGCTGCGGCGTGGAGACACGCTGACCATCGGTGAGGAAAATTTCTGGGTAGAT |
| | CGGGTTTCGCCGGATGATGGCGGAAGTTGTCATCTCTGGCTTGGACGGGGCGTACCGC |
| | CTGCCGTTAACCGTCGCCGCTGAAAGGGGGATGTATGGCCATAAAAGGTCTTGAGCAG |
| | GCCGTTGAAAACCTCAGCCGTATCAGCAAAACGGCGGTGCCTGGTGCCGCCGCAATGG |

| | |
|---|---|
| | CCATTAACCGCGTTGCTTCATCCGCGATATCGCAGTCGGCGTCACAGGTTGCCCGTGAG<br>ACAAAGGTACGCCGGAAACTGGTAAAGGAAAGGGCCAGGCTGAAAAGGGCCACGGTCA<br>AAAATCCGCAGGCCAGAATCAAAGTTAACCGGGGGGATTTGCCCGTAATCAAGCTGGGT<br>AATGCGCGGGTTGTCCTTTCGCGCCGCAGGCGTCGTAAAAAGGGGCAGCGTTCATCCC<br>TGAAAGGTGGCGGCAGCGTGCTTGTGGTGGGTAACCGTCGTATTCCCGGCGCGTTTAT<br>TCAGCAACTGAAAAATGGCCGGTGGCATGTCATGCAGCGTGTGGCTGGGAAAAACCGT<br>TACCCCATTGATGTGGTGAAAATCCCGATGGCGGTGCCGCTGACCACGGCGTTTAAACA<br>AAATATTGAGCGGATACGGCGTGAACGTCTTCCGAAAGAGCTGGGCTATGCGCTGCAG<br>CATCAACTGAGGATGGTAATAAAGCGATGAAACATACTGAACTCCGTGCAGCCGTACTG<br>GATGCACTGGAGAAGCATGACACCGGGGCGACGTTTTTTGATGGTCGCCCCGCTGTTTT<br>TGATGAGGCGGATTTTCCGGCAGTTGCCGTTTATCTCACCGGCGCTGAATACACGGGCG<br>AAGAGCTGGACAGCGATACCTGGCAGGCGGAGCTGCATATCGAAGTTTTCCTGCCTGC<br>TCAGGTGCCGGATTCAGAGCTGGATGCGTGGATGGAGTCCCGGATTTATCCGGTGATG<br>AGCGATATCCCGGCACTGTCAGATTTGATCACCAGTATGGTGGCCAGCGGCTATGACTA<br>CCGGCGCGACGATGATGCGGGCTTGTGGAGTTCAGCCGATCTGACTTATGTCATTACCT<br>ATGAAATGTGAGGACGCTATGCCTGTACCAAATCCTACAATGCCGGTGAAAGGTGCCGG<br>GACCACCCTGTGGGTTTATAAGGGGAGCGGTGACCCTTACGCGAATCCGCTTTCAGAC<br>GTTGACTGGTCGCGTCTGGCAAAAGTTAAAGACCTGACGCCCGGCGAACTGACCGCTG<br>AGTCCTATGACGACAGCTATCTCGATGATGAAGATGCAGACTGGACTGCGACCGGGCA<br>GGGGCAGAAATCTGCCGGAGATACCAGCTTCACGCTGGCGTGGATGCCCGGAGAGCA<br>GGGGCAGCAGGCGCTGCTGGCGTGGTTTAATGAAGGCGATACCCGTGCCTATAAAATC<br>CGCTTCCCGAACGGCACGGTCGATGTGTTCCGTGGCTGGGTCAGCAGTATCGGTAAGG<br>CGGTGACGGCGAAGGAAGTGATCACCCGCACGGTGAAAGTCACCAATGTGGGACGTCC<br>GTCGATGGCAGAAGATCGCAGCACGGTAACAGCGGCAACCGGCATGACCGTGACGCCT<br>GCCAGCACCTCGGTGGTGAAAGGGCAGAGCACCACGCTGACCGTGGCCTTCCAGCCG<br>GAGGGCGTAACCGACAAGAGCTTTCGTGCGGTGTCTGCGGATAAAACAAAAGCCACCG<br>TGTCGGTCAGTGGTATGACCATCACCGTGAACGGCGTTGCTGCAGGCAAGGTCAACATT<br>CCGGTTGTATCCGGTAATGGTGAGTTTGCTGCGGTTGCAGAAATTACCGTCACCGCCAG<br>TTAATCCGGAGAGTCAGCGATGTTCCTGAAAACCGAATCATTTGAACATAACGGTGTGAC<br>CGTCACGCTTTCTGAACTGTCAGCCCTGCAGCGCATTGAGCATCTCGCCCTGATGAAAC<br>GGCAGGCAGAACAGGCGGAGTCAGACAGCAACCGGAAGTTTACTGTGGAAGACGCCAT<br>CAGAACCGGCGCGTTTCTGGTGGCGATGTCCCTGTGGCATAACCATCCGCAGAAGACG<br>CAGATGCCGTCCATGAATGAAGCCGTTAAACAGATTGAGCAGGAAGTGCTTACCACCTG<br>GCCCACGGAGGCAATTTCTCATGCTGAAAACGTGGTGTACCGGCTGTCTGGTATGTATG<br>AGTTTGTGGTGAATAATGCCCCTGAACAGACAGAGGACGCCGGGCCCGCAGAGCCTGT<br>TTCTGCGGGAAAGTGTTCGACGGTGAGCTGAGTTTTGCCCTGAAACTGGCGCGTGAGAT<br>GGGGCGACCCGACTGGCGTGCCATGCTTGCCGGGATGTCATCCACGGAGTATGCCGAC<br>TGGCACCGCTTTTACAGTACCCATTATTTTCATGATGTTCTGCTGGATATGCACTTTTCCG<br>GGCTGACGTACACCGTGCTCAGCCTGTTTTTCAGCGATCCGGATATGCATCCGCTGGAT<br>TTCAGTCTGCTGAACCGGCGCGAGGCTGACGAAGAGCCTGAAGATGATGTGCTGATGC<br>AGAAAGCGGCAGGGCTTGCCGGAGGTGTCCGCTTTGGCCCGGACGGGAATGAAGTTAT<br>CCCCGCTTCCCCGGATGTGGCGGACATGACGGAGGATGACGTAATGCTGATGACAGTA<br>TCAGAAGGGATCGCAGGAGGAGTCCGGTATGGCTGAACCGGTAGGCGATCTGGTCGTT<br>GATTTGAGTCTGGATGCGGCCAGATTTGACGAGCAGATGGCCAGAGTCAGGCGTCATTT<br>TTCTGGTACGGAAAGTGATGCGAAAAAAACAGCGGCAGTCGTTGAACAGTCGCTGAGCC<br>GACAGGCGCTGGCTGCACAGAAAGCGGGGATTTCCGTCGGGCAGTATAAAGCCGCCAT<br>GCGTATGCTGCCTGCACAGTTCACCGACGTGGCCACGCAGCTTGCAGGCGGGCAAAGT<br>CCGTGGCTGATCCTGCTGCAACAGGGGGGGCAGGTGAAGGACTCCTTCGGCGGGATG<br>ATCCCCATGTTCAGGGGGCTTGCCGGTGCGATCACCCTGCCGATGGTGGGGGCCACCT<br>CGCTGGCGGTGGCGACCGGTGCGCTGGCGTATGCCTGGTATCAGGGCAACTCAACCCT<br>GTCCGATTTCAACAAAACGCTGGTCCTTTCCGGCAATCAGGCGGGACTGACGGCAGATC<br>GTATGCTGGTCCTGTCCAGAGCCGGGCAGGCGGCAGGGCTGACGTTTAACCAGACCAG<br>CGAGTCACTCAGCGCACTGGTTAAGGCGGGGGTAAGCGGTGAGGCTCAGATTGCGTCC<br>ATCAGCCAGAGTGTGGCGCGTTTCTCCTCTGCATCCGGCGTGGAGGTGGACAAGGTCG<br>CTGAAGCCTTCGGGAAGCTGACCACAGACCCGACGTCGGGGCTGACGGCGATGGCTC<br>GCCAGTTCCATAACGTGTCGGCGGAGCAGATTGCGTATGTTGCTCAGTTGCAGCGTTCC<br>GGCGATGAAGCCGGGGCATTGCAGGCGGCGAACGAGGCCGCAACGAAAGGGTTTGAT<br>GACCAGACCCGCCGCCTGAAAGAGAACATGGGCACGCTGGAGACCTGGGCAGACAGG<br>ACTGCGCGGGCATTCAAATCCATGTGGGATGCGGTGCTGGATATTGGTCGTCCTGATAC<br>CGCGCAGGAGATGCTGATTAAGGCAGAGGCTGCGTATAAGAAAGCAGACGACATCTGG |

| | |
|---|---|
| | AATCTGCGCAAGGATGATTATTTTGTTAACGATGAAGCGCGGGCGCGTTACTGGGATGA<br>TCGTGAAAAGGCCCGTCTTGCGCTTGAAGCCGCCCGAAAGAAGGCTGAGCAGCAGACT<br>CAACAGGACAAAAATGCGCAGCAGCAGAGCGATACCGAAGCGTCACGGCTGAAATATA<br>CCGAAGAGGCGCAGAAGGCTTACGAACGGCTGCAGACGCCGCTGGAGAAATATACCGC<br>CCGTCAGGAAGAACTGAACAAGGCACTGAAAGACGGGAAAATCCTGCAGGCGGATTAC<br>AACACGCTGATGGCGGCGGCGAAAAAGGATTATGAAGCGACGCTGAAAAAGCCGAAAC<br>AGTCCAGCGTGAAGGTGTCTGCGGGCGATCGTCAGGAAGACAGTGCTCATGCTGCCCT<br>GCTGACGCTTCAGGCAGAACTCCGGACGCTGGAGAAGCATGCCGGAGCAAATGAGAAA<br>ATCAGCCAGCAGCGCCGGGATTTGTGGAAGGCGGAGAGTCAGTTCGCGGTACTGGAG<br>GAGGCGGCGCAACGTCGCCAGCTGTCTGCACAGGAGAAATCCCTGCTGGCGCATAAAG<br>ATGAGACGCTGGAGTACAAACGCCAGCTGGCTGCACTTGGCGACAAGGTTACGTATCA<br>GGAGCGCCTGAACGCGCTGGCGCAGCAGGCGGATAAATTCGCACAGCAGCAACGGGC<br>AAAACGGGCCGCCATTGATGCGAAAAGCCGGGGGCTGACTGACCGGCAGGCAGAACG<br>GGAAGCCACGGAACAGCGCCTGAAGGAACAGTATGGCGATAATCCGCTGGCGCTGAAT<br>AACGTCATGTCAGAGCAGAAAAAGACCTGGGCGGCTGAAGACCAGCTTCGCGGGAACT<br>GGATGGCAGGCCTGAAGTCCGGCTGGAGTGAGTGGGAAGAGAGCGCCACGGACAGTA<br>TGTCGCAGGTAAAAAGTGCAGCCACGCAGACCTTTGATGGTATTGCACAGAATATGGCG<br>GCGATGCTGACCGGCAGTGAGCAGAACTGGCGCAGCTTCACCCGTTCCGTGCTGTCCA<br>TGATGACAGAAATTCTGCTTAAGCAGGCAATGGTGGGGATTGTCGGGAGTATCGGCAGC<br>GCCATTGGCGGGGCTGTTGGTGGCGGCGCATCCGCGTCAGGCGGTACAGCCATTCAG<br>GCCGCTGCGGCGAAATTCCATTTTGCAACCGGAGGATTTACGGGAACCGGCGGCAAAT<br>ATGAGCCAGCGGGGATTGTTCACCGTGGTGAGTTTGTCTTCACGAAGGAGGCAACCAG<br>CCGGATTGGCGTGGGGAATCTTTACCGGCTGATGCGCGGCTATGCCACCGGCGGTTAT<br>GTCGGTACACCGGGCAGCATGGCAGACAGCCGGTCGCAGGCGTCCGGGACGTTTGAG<br>CAGAATAACCATGTGGTGATTAACAACGACGGCACGAACGGGCAGATAGGTCCGGCTG<br>CTCTGAAGGCGGTGTATGACATGGCCCGCAAGGGTGCCCGTGATGAAATTCAGACACA<br>GATGCGTGATGGTGGCCTGTTCTCCGGAGGTGGACGATGAAGACCTTCCGCTGGAAAG<br>TGAAACCCGGTATGGATGTGGCTTCGGTCCCTTCTGTAAGAAAGGTGCGCTTTGGTGAT<br>GGCTATTCTCAGCGAGCGCCTGCCGGGCTGAATGCCAACCTGAAAACGTACAGCGTGA<br>CGCTTTCTGTCCCCCGTGAGGAGGCCACGGTACTGGAGTCGTTTCTGGAAGAGCACGG<br>GGGCTGGAAATCCTTTCTGTGGACGCCGCCTTATGAGTGGCGGCAGATAAAGGTGACC<br>TGCGCAAAATGGTCGTCGCGGGTCAGTATGCTGCGTGTTGAGTTCAGCGCAGAGTTTGA<br>ACAGGTGGTGAACTGATGCAGGATATCCGGCAGGAAACACTGAATGAATGCACCCGTG<br>CGGAGCAGTCGGCCAGCGTGGTGCTCTGGGAAATCGACCTGACAGAGGTCGGTGGAG<br>AACGTTATTTTTTCTGTAATGAGCAGAACGAAAAAGGTGAGCCGGTCACCTGGCAGGGG<br>CGACAGTATCAGCCGTATCCCATTCAGGGGAGCGGTTTTGAACTGAATGGCAAAGGCAC<br>CAGTACGCGCCCCACGCTGACGGTTTCTAACCTGTACGGTATGGTCACCGGGATGGCG<br>GAAGATATGCAGAGTCTGGTCGGCGGAACGGTGGTCCGGCGTAAGGTTTACGCCCGTT<br>TTCTGGATGCGGTGAACTTCGTCAACGGAAACAGTTACGCCGATCCGGAGCAGGAGGT<br>GATCAGCCGCTGGCGCATTGAGCAGTGCAGCGAACTGAGCGCGGTGAGTGCCTCCTTT<br>GTACTGTCCACGCCGACGGAAACGGATGGCGCTGTTTTTCCGGGACGTATCATGCTGG<br>CCAACACCTGCACCTGGACCTATCGCGGTGACGAGTGCGGTTATAGCGGTCCGGCTGT<br>CGCGGATGAATATGACCAGCCAACGTCCGATATCACGAAGGATAAATGCAGCAAATGCC<br>TGAGCGGTTGTAAGTTCCGCAATAACGTCGGCAACTTTGGCGGCTTCCTTTCCATTAACA<br>AACTTTCGCAGTAAATCCCATGACACAGACAGAATCAGCGATTCTGGCGCACGCCCGGC<br>GATGTGCGCCAGCGGAGTCGTGCGGCTTCGTGGTAAGCACGCCGGAGGGGGAAAGAT<br>ATTTCCCCTGCGTGAATATCTCCGGTGAGCCGGAGGCGTATTTCCGTATGTCGCCGGAA<br>GACTGGCTGCAGGCAGAAATGCAGGGTGAGATTGTGGCGCTGGTCCACAGCCACCCCG<br>GTGGTCTGCCCTGGCTGAGTGAGGCCGACCGGCGGCTGCAGGTGCAGAGTGATTTGC<br>CGTGGTGGCTGGTCTGCCGGGGGACGATTCATAAGTTCCGCTGTGTGCCGCATCTCAC<br>CGGGCGGCGCTTTGAGCACGGTGTGACGGACTGTTACACACTGTTCCGGGC |
